# Directing Cell Fate via Spatiotemporally Patterned Viscoelasticity Driven by Dynamic Crosslink Exchange

**DOI:** 10.64898/2026.11.10.764845

**Authors:** Yuran Wang, Albert C. Liu, Ping Zhou, Tae Yoon Kwon, Lucy T. Norris, Dengyang Lu, Christopher M. Madl

## Abstract

The spatially and temporally varied mechanical and biochemical properties of the extracellular matrix (ECM) are critical regulators of cell function. Despite the emerging importance of ECM viscoelasticity in dictating cell fate, suitable biomaterials systems enabling spatiotemporal control over matrix viscoelasticity to direct cell function in physiologically relevant 3D contexts are lacking. A dynamic covalent hydrogel platform was developed that enables user-defined, *in situ* modulation of stress relaxation kinetics decoupled from bulk network stiffness via self-contained, radical-free photochemistries. By introducing photocaged alkoxyamines into initially rapidly relaxing hydrazone-based networks, the viscoelasticity of the networks can be tuned on demand by converting the hydrazone crosslinks to more stable oxime crosslinks following light exposure. This hydrazone-to-oxime bond exchange drives a dramatic shift in stress relaxation half time from approximately ∼10^2^ s to ∼10^4^ s within a single, continuous matrix, without altering stiffness or introducing cytotoxicity. This platform enabled real-time control over cellular adaptations to spatiotemporally patterned viscoelastic mechanical cues, directing fibroblast morphological transitions in 2D, human neural progenitor cell stemness maintenance in 3D, and neurite outgrowth from encapsulated motor neuron spheroids. This phototunable hydrogel system represents a robust platform for selectively patterning viscoelastic cues to direct mechanosensitive cell fate transitions.

## 1. Introduction

Cells are surrounded by a polymeric meshwork known as the extracellular matrix (ECM). The native ECM presents a spatially heterogeneous and temporally varied combination of biochemical and mechanical cues that dictate crucial cellular behaviors, spanning proliferation and migration to stem cell differentiation. [1], [2], [3] Accordingly, pioneering efforts have developed techniques to spatiotemporally pattern both biochemical [4], [5], [6], [7] and elastic mechanical properties (i.e., stiffness) [8], [9], [10], [11] using light-responsive chemistries. These systems have uncovered mechanisms of cellular mechanical memory,[12], [13] directed morphogenesis of 3D organotypic cultures,[14] and even regulated vascularization in vivo.[7]

An expanding body of work has added ECM viscoelasticity as another critical mechanical regulator of cellular behavior.[2] Beyond controlling cell spreading, migration, and mechanosensing in 2D culture systems,[15], [16], [17] viscoelastic properties such as stress relaxation and creep are potent regulators of cell fate in physiologically relevant 3D contexts. Stress relaxation rate in 3D hydrogels dictates mesenchymal stromal cell differentiation,[18] neural stem cell maintenance and lineage specification,[19], [20], [21] intestinal organoid morphogenesis,[22] and axon extension from neurons.[23] Despite the importance of matrix viscoelasticity in regulating cell fate in 3D contexts, the ability to spatiotemporally pattern physiologically relevant viscoelastic cues to direct cellular function in 3D remains severely limited.

Prior work has laid the foundation for photo-patterning of viscoelasticity in hydrogel materials,[24], [25], [26], [27] though these approaches have largely focused on 2D cell culture, rely on free radical mediated reactions, and are limited to readouts of cellular spreading and mechanosensing. To leverage viscoelasticity to regulate cellular function in 3D contexts, a hydrogel system must (1) use bioselective, radical-free chemistries to exercise spatiotemporal control, (2) be self-contained, not requiring the addition of exogenous chemical agents, (3) exhibit broadly tunable viscoelastic properties over a physiologically relevant range, and (4) remain stable under extended cell culture conditions (**Fig 1A**).

**Figure 1.**
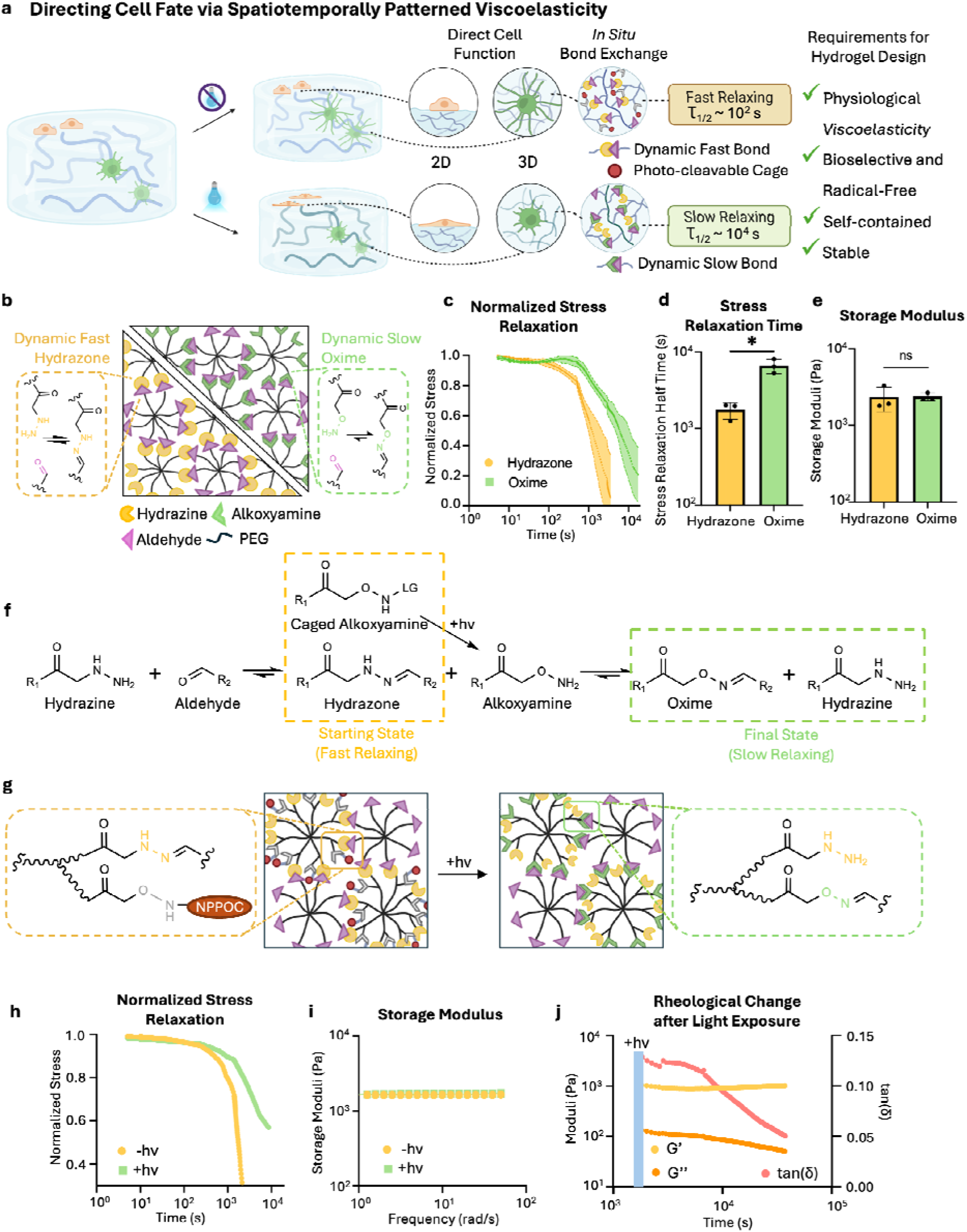
Engineering a hydrogel platform with selective, spatiotemporal control of viscoelasticity via hydrazone/oxime exchange. **(a)** Schematic of hydrogel platform with on-demand tuning of bond exchange kinetics to spatiotemporally pattern viscoelasticity, with design requirements to enable 3D control of cell fate. **(b)** Schematic illustrating 8-arm PEG-based hydrogels with adaptable covalent crosslinks, highlighting that hydrazone bonds exchange rapidly, while oxime bonds exchange much more slowly. Rheological characterization showing **(c)** storage moduli, **(d)** stress relaxation half times, and **(e)** normalized stress relaxation curves of 3.5% (w/v) 8-arm PEG hydrazone and oxime hydrogels. Rheology data are presented as mean ± s.d., n=3. *p<0.05, Welch’s t test. **(f)** Schematic depicting the chemistry and kinetic rationale of a hydrazone/oxime crosslink-based hydrogel capable of achieving on-demand, *in-situ* tuning of stress relaxation without altering network stiffness. **(g)** Schematic depicting PEG-based hydrogel system with on-demand tuning of bond exchange kinetics, with initially hydrazone-based crosslinks being replaced by oxime crosslinks after photo-triggered release of NPPOC protecting groups. After photo-activation, **(h)** stress relaxation time increases by an order of magnitude, while **(i)** storage modulus remains unchanged. **(j)** Representative rheological characterization showing evolution of loss modulus and tan(δ) over time after irradiation.

To achieve these objectives, we engineered a hydrogel system that leverages the distinct bond exchange kinetics and thermodynamic stabilities of hydrazone and oxime dynamic covalent crosslinks to enable user-defined, on-demand modulation of stress relaxation timescale. This system utilizes photomediated oxime chemistry,[28], [29] wherein photoactivation reveals alkoxyamine groups that participate in dynamic crosslink exchange and introduce slower-relaxing oxime bonds within a pre-established hydrazone-based hydrogel network. The resulting transition in matrix relaxation kinetics occurs without altering the overall network architecture, maintaining consistent stiffness and preventing unintended changes in hydrogel swelling. Not only does this system capture viscoelasticity-induced cellular morphological transitions in 2D contexts, it also enables spatiotemporal control over diverse and physiologically relevant cellular functions in 3D, including maintenance of human neural progenitor cell stemness and differentiation capacity and guidance of neurite outgrowth from human motor neuron spheroids. Thus, this platform goes beyond elucidating molecular mechanisms of cellular mechanosensing and enables user-directed growth of engineered cell and tissue constructs.

## 2. Results

### 2.1. Designing Hydrogels with On-Demand Tuning of Physiological Stress Relaxation Timescales via Dynamic Covalent Bond Exchange

Dynamic covalent chemistries based on hydrazone and oxime bonds enable fabrication of viscoelastic hydrogels spanning a broad and physiologically relevant range of stress relaxation times.[23], [28], [30], [31] Hydrazone and oxime crosslinks exhibit markedly different thermodynamic stability and exchange kinetics, with hydrazones displaying lower equilibrium constants and faster dissociation rates (K_eq_ ∼ 10^4^ - 10^5^ M^-1^; k_off_ ∼ 10^-3^ - 10^-4^ s^-1^) compared to oximes (K_eq_ ∼ 10^8^ - 10^9^ M^-1^; k_off_ ∼ 10^-7^ - 10^-9^ s^-1^)[32], [33], [34] (**Figure 1b, Supplementary Table 1**). These discrepancies in kinetic parameters and hydrolytic stability govern the viscoelastic mechanical properties of the resulting polymer networks, while sufficiently high thermodynamic equilibrium constants afford stable crosslink density and thus stiffness. We therefore postulated that hydrogels in which initially rapidly relaxing hydrazone crosslinks were replaced by slowly relaxing oxime crosslinks would enable on-demand tuning of relaxation timescales without altering stiffness.

To validate this approach, we first synthesized purely hydrazone- and oxime-crosslinked multi-arm poly(ethylene glycol) (PEG) hydrogels (**Figure 1b-e**) and characterized their viscoelastic behavior via oscillatory shear rheology. Maintaining a constant crosslinking density yielded hydrogels with indistinguishable shear storage moduli (i.e. the same bulk stiffness; **Figure 1e**). In contrast, stress relaxation tests revealed that oxime-crosslinked hydrogels undergo stress relaxation approximately one order of magnitude slower than hydrazone-based hydrogels (**Figure 1c,d**). Together, these results establish that matrix stress relaxation rates can be decoupled from hydrogel stiffness through dynamic covalent crosslink chemistry, motivating the development of a biomaterial platform capable of *in situ*, on-demand tuning of viscoelasticity.

Reasoning that oximes serve as not only slower relaxing, but also thermodynamically favored, crosslinks, we envisioned a hydrogel initially formed through hydrazone crosslinking while incorporating photocaged alkoxyamines as latent oxime precursors (**Figure 1f,g**). Upon photoactivation, the liberated alkoxyamines would drive dynamic crosslink exchange toward oxime formation, thereby slowing hydrogel stress relaxation. The hydrazone-to-oxime exchange is achieved by sequential hydrazone hydrolysis and oxime formation, and the rates and stabilities of these reactions govern the direction and speed of the transition. To validate the feasibility of this approach, we utilized reported thermodynamic and kinetic constants to predict the timescale of this exchange (see Supporting Information)[32], [33], [34], [35], [36]. Our numerical simulations were consistent with bond exchange occurring over the course of ∼10 hours (**Figure S1**), which is a reasonable timescale for cell culture applications. Thus, we proceeded to design a hydrogel system that contained both hydrazone crosslinks and photo-caged alkoxyamines.

To implement this design, each arm of a 4-arm PEG was sequentially functionalized via amide coupling and copper(I)-catalyzed azide-alkyne cycloaddition (CuAAC) to present both hydrazine and photocaged alkoxyamine end groups, respectively (**Figure S2**). This branched architecture was designed to maintain stoichiometric equivalence between the hydrazines and alkoxyamines. 2-(2-Nitrophenyl)-propyloxycarbonyl (NPPOC) was selected as the alkoxyamine photocaging group, as it is a cytocompatible alkoxyamine photo-protecting group that undergoes efficient cleavage upon exposure to 365 nm light.[28] The multi-step synthesis resulted in the desired bifunctional product, PEG-[4Hz/4HNO-NPPOC] (**Figures S3-7**) in good yield (52% over 5 steps). The bifunctional PEG-[4Hz/4HNO-NPPOC] polymers were subsequently crosslinked with 8-arm aldehyde PEGs to yield viscoelastic hydrogels with initially rapid stress relaxation (τ_1/2_∼1750 s, **Figure 1h**). Following 5 min of exposure to 365 nm light (12 mW/cm^2^) and 12 hours of equilibration, the stress relaxation timescale increased by an order of magnitude (τ_1/2_∼10^4^ s, **Figure 1h**). At the same time, the storage moduli of the hydrogels remained unchanged, confirming that stress relaxation time can be dynamically tuned independently of stiffness (**Figure 1i**). Tracking the evolution of the viscoelastic properties of the hydrogels post-irradiation revealed that the loss moduli and tan(δ) decreased over the course of 10 hours (**Figure 1j**), consistent with our numerical predictions (**Figure S1**), while the storage moduli remained roughly constant. Together, these results demonstrate that photo-triggered hydrazone to oxime conversion is an attractive approach to spatiotemporally pattern stress relaxation timescale to direct cell fate.

### 2.2. Fabrication of Photoresponsive Engineered Protein Hydrogels for Spatiotemporal Control of Stress Relaxation Time

To establish a platform with on-demand tuning of matrix stress relaxation kinetics capable of directing cellular functions, we developed a two-component hydrogel system composed of recombinant elastin-like proteins (ELPs) and multi-arm PEG crosslinkers (**Figure 2a,b**). While PEG was selected as the primary crosslinking species due to its established biocompatibility and capacity for robust, quantitative chemical functionalization, it lacks intrinsic bioactive cues required to support cell-ECM interactions. In contrast, ELPs can be engineered to integrate biomimetic functionality into the polymer backbones. The ELP selected for the present system features bioactive domains containing integrin-binding RGD sequences and susceptibility to proteolytic degradation separated by elastin-like structural domains.[1], [19] Specific lysine residues located within the elastin-like domains present sites for chemical modification to enable subsequent crosslinking without interfering with bioactivity. To obtain hydrogels with on-demand tuning of stress relaxation, the initial hybrid ELP-PEG networks are crosslinked via hydrazone bonds formed from reactions between aldehydes on the ELPs and hydrazines on the PEG crosslinkers. Meanwhile, competitive alkoxyamine functionalities are presented on the PEG crosslinkers but are initially rendered unreactive via a photo-cleavable protecting group. Upon targeted light exposure to deprotect the alkoxyamines, dynamic bond exchange drives a selective conversion toward thermodynamically favored, more stable oxime crosslinks.

**Figure 2.**
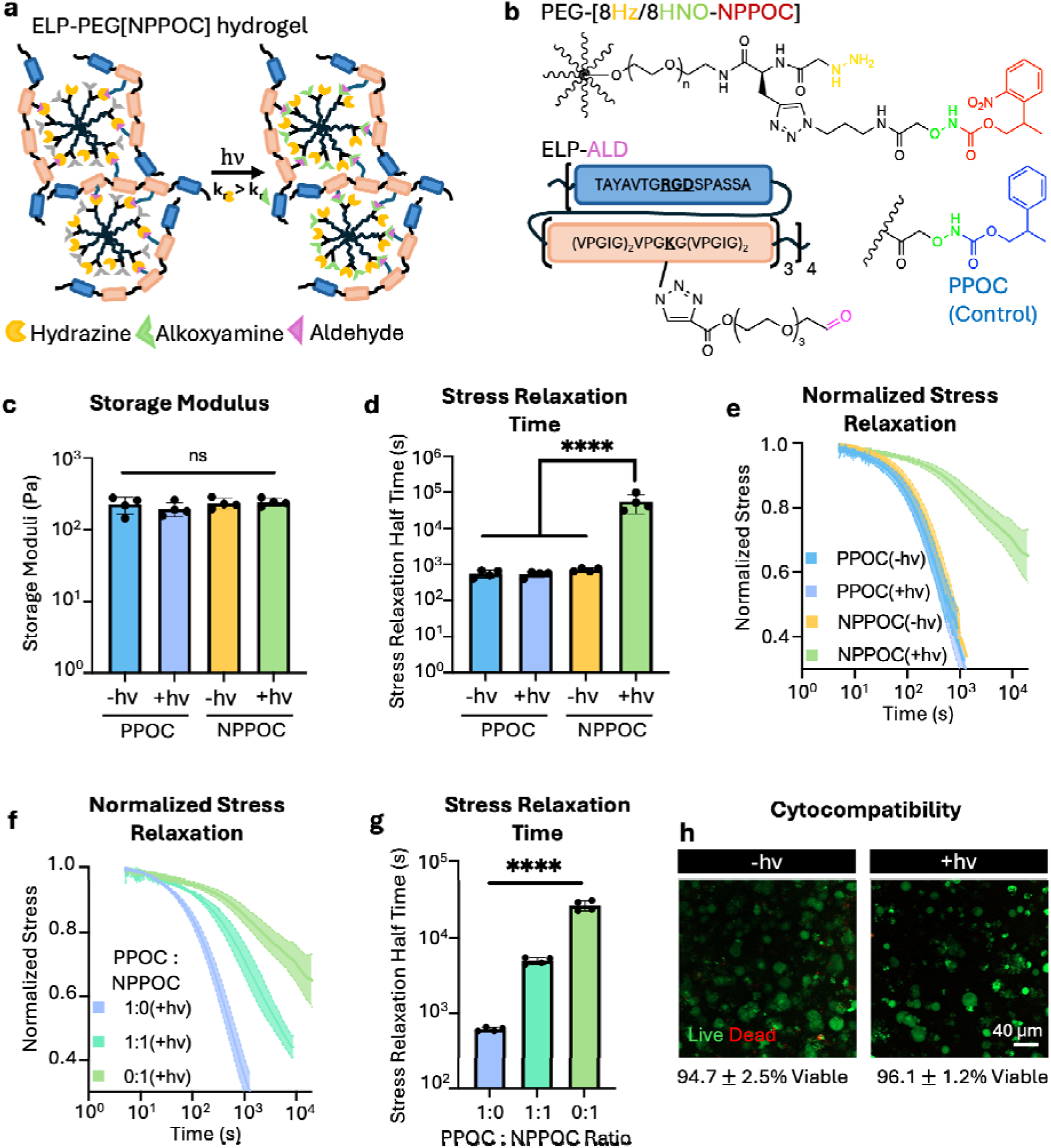
Engineering biocompatible hydrogels with selective, on-demand tuning of stress relaxation. **(a)** Schematic illustrating the light-induced transformation from hydrazone to oxime crosslinks in hybrid ELP-PEG networks. **(b)** Chemical structures of hydrogel components: PEG-hydrazine-alkoxyamine-NPPOC (PEG-[8Hz/8HNO-NPPOC]), the analogous photo-inert control PPOC-protected PEG material, and ELP-aldehyde (ELP-ALD). Rheological characterization showing **(c)** storage moduli, **(d)** stress relaxation half time, and **(e)** normalized stress relaxation curves of 3% (w/v) ELP-PEG hydrogels, demonstrating that NPPOC hydrogels relax significantly slower after light exposure without altering network stiffness, while both stress relaxation and stiffness remain unchanged in control PPOC hydrogels. **(f, g)** Graded tunability of stress relaxation achieved by blending NPPOC- and PPOC-protected PEGs at 1:1 ratio. **(h)** Representative confocal fluorescence images and quantified cell viability of NPCs encapsulated within phototunable ELP-PEG hydrogels, imaged 1 day post-light exposure (scale bar = 40 μm). Data (c–h) are presented as mean ± s.d. (n = 4, ****p < 0.0001, lognormal two-way ANOVA with Tukey *post hoc* testing).

To prepare the hydrazone and NPPOC-caged dual-functionality crosslinkers, a similar amide coupling and CuAAC approach was followed as above. The five-step synthesis resulted in the desired bifunctional product, PEG-[8Hz/8HNO-NPPOC] (**Figures S8-12**). To establish a photochemically inert control material to isolate potential confounding effects introduced by light exposure, we synthesized a variant crosslinking macromer utilizing a structurally analogous 2-phenylpropyloxycarbonyl (PPOC) protecting group. The PPOC group lacks the photosensitive nitro moiety of the NPPOC group, rendering the control PEG-[8Hz/8HNO-PPOC] material incapable of light-mediated deprotection (**Figure S13**). Both PEG crosslinkers were prepared in good yield (55-60% over 5 steps) with a 1:1 ratio of hydrazines to protected alkoxyamines, as characterized by ^1^H NMR (**Figures S14,15**).

The functionalization of ELPs with pendant aldehyde groups presented a separate synthetic challenge. While primary amines are the most tractable sites for chemical modification, their inherent reactivity with aldehydes complicates direct functionalization. Thus, existing dynamic covalent hydrogel systems incorporating ELPs typically confine the aldehyde functionality to non-protein components while modifying the ELP with the corresponding hydrazine groups,[37], [38], [39] limiting the architectural versatility of the resulting hydrogel networks. Other strategies to introduce aldehydes to recombinant proteins, such as enzymatic modification[40] and non-canonical amino acid incorporation,[41] are limited to one or a few sites in the proteins and are thus not suitable for crosslinking reactions. Therefore, to functionalize the ELP component with aldehydes, lysine residues were first converted into azides via aqueous diazo transfer,[42] followed by CuAAC with a heterobifunctional linker possessing both a terminal alkyne and an aliphatic aldehyde (**Figure S16**). The alkyne terminus of the linker presents a propiolate group, as the electron-withdrawing ester group accelerates the reaction with azides,[43] thereby requiring lower copper loading and preserving the reactivity of the aldehyde moiety (see Supporting Information). The product, ELP-ALD, was obtained in good yield (80% over two steps) with complete conversion of primary amines to aldehydes based on ^1^H NMR (**Figures S17,18**).

### 2.3. Phototriggered Crosslink Exchange Enables Selective, Physiological Stress Relaxation Tuning

To validate that the hybrid ELP-PEG system afforded selective and physiologically relevant tuning of stress relaxation timescale, rheological characterization was performed on hydrogels crosslinked with either photoreactive PEG-[8Hz/8HNO-NPPOC] or the photo-inert control PEG-[8Hz/8HNO-PPOC] at a 1:1 molar ratio of reactive groups. Across all experimental groups, the critical gelation point (G’>G”) was achieved within 2 min following precursor mixing, and 30 min was sufficient to reach a stable plateau storage modulus (**Figure S19**). These rapid gelation kinetics underscore the applicability of this hydrogel platform as both a 2D cell culture substrate and a 3D scaffold for cell encapsulation. All hydrogels exhibited comparable storage moduli (G’∼250 Pa) (**Figure 2c**), demonstrating consistent stiffness across formulations prior to light exposure.

Following 365 nm irradiation (5 min, 12 mW/cm^2^), NPPOC-containing hydrogels exhibited dramatic changes in stress relaxation behavior, with an approximately 50-fold increase in relaxation half time (τ_1/2_), from ∼600 s before irradiation to over 30,000 s after irradiation (**Figure 2d, e**), spanning relaxation timescales similar to native tissues.[2], [3], [18], [20] By tracking changes in hydrogel loss modulus, we validated that crosslink exchange occurred over ∼10 hours, as predicted based on thermodynamic and kinetic constants (**Figure S1**). Importantly, photo-inert PPOC control hydrogels displayed no significant change in relaxation behavior following light exposure, consistent with the observed changes in relaxation behavior in NPPOC-containing hydrogels arising from photo-uncaging of alkoxyamines, rather than nonspecific effects of irradiation. As the overall crosslink density should be conserved after bond exchange, the stiffness of the networks should remain constant. Consistent with this mechanism, the storage moduli of the NPPOC gels remained unchanged after light exposure (**Figure 2c**), indicating that photo-deprotection did not significantly alter the elastic properties of hydrogel matrices. Further demonstrating decoupled tuning of stress relaxation, the hydrogels did not exhibit appreciable changes in swelling as a function of irradiation (**Figure S20**).

To fine tune the extent of stress relaxation modulation, the amount of hydrazone-to-oxime exchange could be further controlled by varying the ratio of photoreactive to photo-inert PEG crosslinkers. Hydrogels formulated with an intermediate 1:1 molar ratio of NPPOC- and PPOC-protected PEGs exhibited an intermediate relaxation half time of approximately 5000 s after light exposure, demonstrating graded tuning of network viscoelasticity (**Figure 2f, g**). To confirm that this photochemical approach to tuning relaxation behavior is cytocompatible, human neural progenitor cells (NPCs) were encapsulated in the NPPOC hydrogels as a representative therapeutically relevant cell type, and cell viability was assessed with and without light exposure via Live/Dead staining. These cytocompatibility assays confirmed that the phototunable hydrogels maintain high cell viability (∼95%) for encapsulated cells (**Figure 2h**). Together, these results establish the hybrid ELP-PEG hydrogel platform as a cytocompatible biomaterial system that enables precise, light-mediated control of matrix stress relaxation while preserving bulk stiffness, providing a tunable framework to investigate and control cellular responses to dynamic mechanical cues.

### 2.4. Spatiotemporal Tuning of Stress Relaxation Regulates Cell Morphology in 2D

Cytoskeletal dynamics respond to substrate viscoelasticity, with substrate stress relaxation behavior regulating processes such as cell spreading and migration.[15], [34], [44], [45], [46] Thus, to assess whether cells respond to spatiotemporally patterned relaxation times, we cultured MRC5 human lung fibroblasts stably expressing the fluorescent actin reporter LifeAct-iRFP670 on our phototunable ELP-PEG hydrogels. To confirm that the hydrogels access cell-relevant relaxation timescales, fibroblasts were seeded onto 2D hydrogel substrates that were either previously exposed to 365 nm light (to yield slowly relaxing materials) or left unexposed (to retain rapid relaxation timescale) (**Figure 3a**). Slowly relaxing samples were irradiated 1-hour post-mixing to ensure completion of initial gelation. Prior to cell seeding, all hydrogel formulations were incubated in PBS at 37 °C for 24 hours to allow the dynamic networks to equilibrate. Fluorescence imaging and subsequent morphometric quantification revealed that fibroblasts cultured on slowly relaxing substrates adopted a highly elongated, spindle-shaped morphology, exhibiting significantly higher aspect ratio (i.e., elongation) and lower form factor (i.e., increase in perimeter irregularity driven by localized membrane deformations) compared to cells on rapidly relaxing substrates (**Figure 3b-d**). This elongated morphology is consistent with previous observations that slower-relaxing matrices better preserve cytoskeletal tension by resisting stress dissipation to promote focal adhesion maturation and support more spread and elongated cellular phenotypes.[2], [18] These results confirm that cells can sense and respond to the stress relaxation timescales accessed by our hydrogel platform.

**Figure 3.**
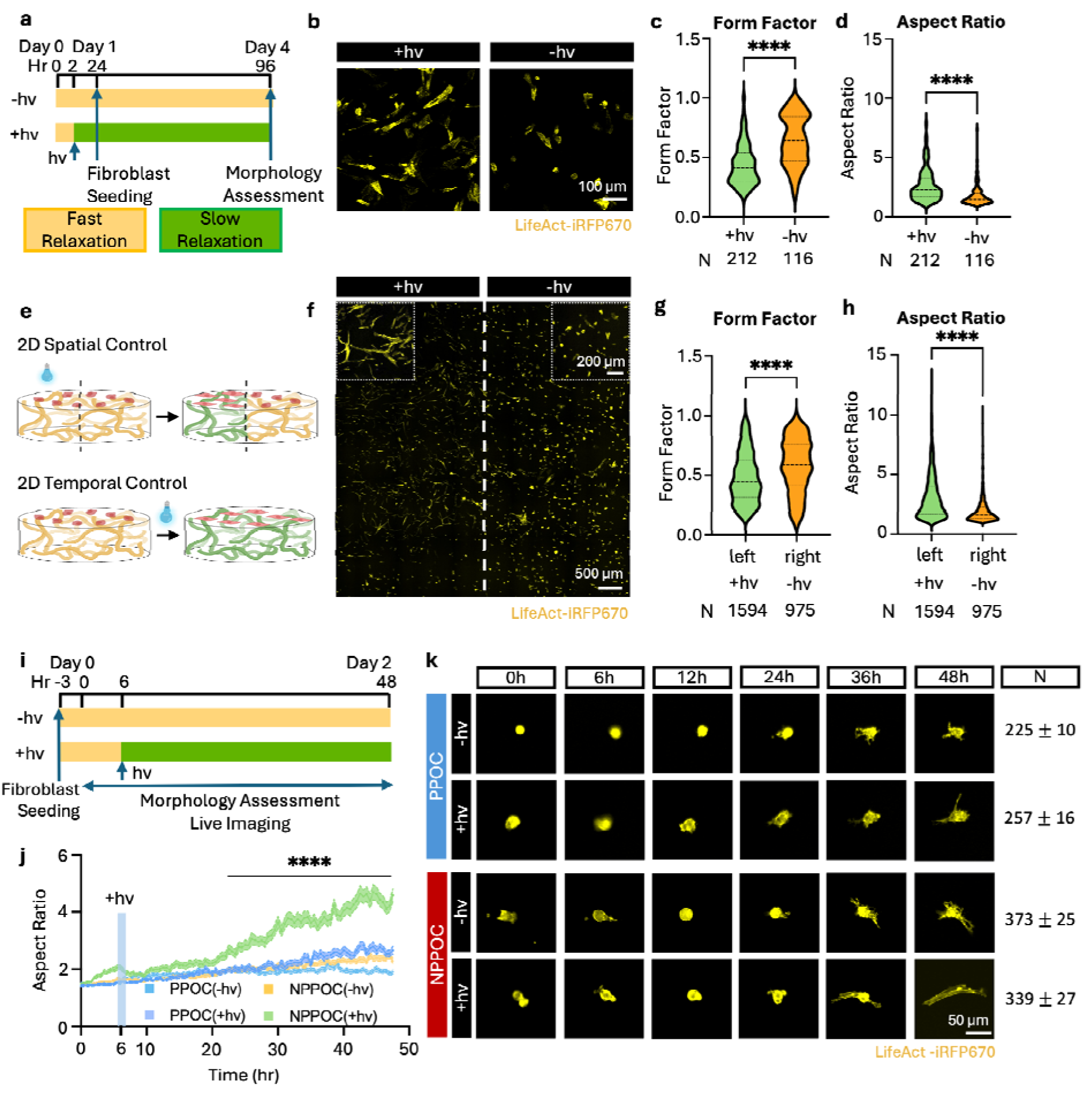
Hydrogels with photo-triggered bond exchange enable spatiotemporal control over network viscoelasticity to modulate cell morphology. **(a)** Experimental timeline for evaluating fibroblast morphological differences on pre-treated hydrogel substrates. **(b)** Representative widefield fluorescence images and quantification of **(c)** form factor and **(d)** aspect ratio of LifeAct-iRFP670 MRC5 fibroblasts cultured on rapidly relaxing (-hv) and slowly relaxing (+hv) 2D hydrogel substrates at day 3 post-seeding. **(e)** Schematics depicting the experimental designs for spatial and temporal effects on fibroblast morphology via phototunable hydrogels. **(f)** Representative widefield fluorescence images and quantification of **(g)** form factor and **(h)** aspect ratio of fibroblasts cultured on spatially patterned NPPOC-protected ELP-PEG hydrogel substrates presenting distinct mechanical zones: a slowly relaxing left half (+hv) and a rapidly relaxing right half (-hv). **(i)** Experimental timeline for live-cell imaging to monitor the real-time morphological response of fibroblasts to *in situ* modulation of substrate stress relaxation. **(j)** Quantified time-course of the aspect ratio and **(k)** representative time-lapse widefield fluorescence images of fibroblast morphology over time in response to light-triggered changes in substrate relaxation time. Data in (c, d, g, h) are presented as violin plots (the black dashed lines represent median values and dotted lines represent quartiles) and analyzed via unpaired t-test (****p < 0.0001). Data in (j) are presented as mean ± s.e.m. and analyzed via two-way ANOVA with Tukey *post hoc* testing (****p < 0.0001).

Next, to demonstrate the spatial tunability of our platform, we selectively irradiated regions of 2D NPPOC-protected ELP-PEG hydrogel substrates to generate localized changes in substrate stress relaxation behavior (**Figure 3e**). Across a continuous 2D hydrogel substrate, the left half was exposed to 365 nm light while the right half was shielded from light, resulting in spatially restricted hydrazone-to-oxime conversion. Fibroblasts cultured across this patterned interface exhibited distinct morphological phenotypes based on their local substrate conditions (**Figure 3f-h**). Cells within light-exposed regions adopted the morphology associated with slowly relaxing substrates, whereas cells in non-irradiated regions maintained the phenotype observed on faster-relaxing substrates. These results demonstrate that the ELP-PEG platform enables spatially patterned control of matrix stress relaxation, allowing cells to respond to programmed heterogeneous mechanical properties across a continuous bulk material.

Beyond spatial patterning, the *in situ* tunability of this platform uniquely enables the real-time observation of cellular adaptation following temporally controlled changes in substrate viscoelasticity. To investigate dynamic cellular responses, fibroblasts were seeded onto equilibrated 2D NPPOC- or PPOC-protected hydrogel substrates and allowed to adhere and stabilize for 3 hours prior to live imaging (**Figure 3i**). For the first 6 hours of imaging, samples were observed under their initial, rapidly relaxing conditions to establish baseline morphologies, after which subsets of the NPPOC- and PPOC-protected samples were either irradiated with 365 nm light *in situ* or left unexposed. Live cell imaging continued for a total of 48 hours to capture cellular adaptation following the light-triggered change in stress relaxation behavior. The fibroblasts demonstrated clear morphological adaptation in response to the temporal shift in relaxation time (**Figure 3j,k**, **Supplementary Videos 1–4**). Across the three persistently fast-relaxing control conditions, including irradiated PPOC-protected materials, fibroblasts maintained a rounded morphology with only fine protrusions. In contrast, fibroblasts cultured on light-activated, initially NPPOC-protected substrates progressively transitioned towards a spindle-like morphology. Morphometric quantification revealed that by 15 hours post-irradiation, fibroblasts on the light-activated, slowly relaxing substrates began to adopt a spindle-like shape, exhibiting significantly higher elongation compared to the three fast-relaxing controls. This morphological divergence continued to widen as the experiment progressed, with cellular spreading and elongation stabilizing after ∼42 hours of imaging. This real-time morphological evolution corroborates the temporal tunability of the NPPOC-protected hydrogel networks. Furthermore, the temporal delay in morphological adaptation aligns with the underlying kinetics of the hydrazone-to-oxime bond exchange and the requisite timescales for cytoskeletal remodeling. Collectively, these findings validate that the NPPOC-protected ELP-PEG platform affords spatiotemporal control over matrix stress relaxation, enabling the study of cellular responses to spatially heterogeneous and dynamically evolving viscoelastic environments.

### 2.5. Temporal Modulation of Stress Relaxation Behavior Directs Neural Progenitor Cell Fate in 3D

Having demonstrated that the phototunable ELP-PEG platform enables dynamic regulation of cellular behavior in response to changes in stress relaxation timescale in 2D culture, we next evaluated its utility for controlling cell behavior in 3D microenvironments. Neural progenitor cells (NPCs) provide an ideal model for this purpose because their self-renewal and differentiation are highly sensitive to matrix viscoelasticity.[19], [20], [37], [47], [48], [49] We have previously demonstrated that rapid stress relaxation promotes NPC stemness by permitting cell-mediated physical (i.e., viscoplastic) matrix remodeling to enable cell-cell contact, whereas culture in purely elastic matrices without other modes of network remodeling results in a rapid loss of stemness.[19], [37] Importantly, matrix remodeling by NPCs is inherently temporal. When NPCs are encapsulated as single cells, they transiently lose expression of stemness markers within 1 day and then subsequently regain expression beginning by day 3 as the cells remodel the material and reestablish cell-cell contacts.[19]

Therefore, we hypothesized that varying the time post-encapsulation at which the microenvironment is transitioned to a slowly relaxing state would regulate the cells’ ability to establish cell-cell contacts and maintain stemness. Human NPCs were encapsulated at a density of 3×10^7^ cells/mL within phototunable NPPOC-protected or photo-inert control PPOC-protected hydrogels and maintained in neural stem cell maintenance medium for 7 days (**Figure 4a, b**). Hydrogels were formulated to have an initial storage modulus (G’) of ∼ 250 Pa, within the physiological range of healthy brain tissue[30] and consistent with our prior work.[37] To investigate the effects of altering stress relaxation, and thus physical remodelability, at different times, samples were irradiated with 365 nm light at day 1 or day 3 post-encapsulation or maintained in an unexposed, rapidly relaxing state for the full 7 days of culture. This strategy enabled precise control over the duration for which encapsulated cells experienced a permissive, remodelable matrix before transitioning to a slower-relaxing microenvironment.

**Figure 4.**
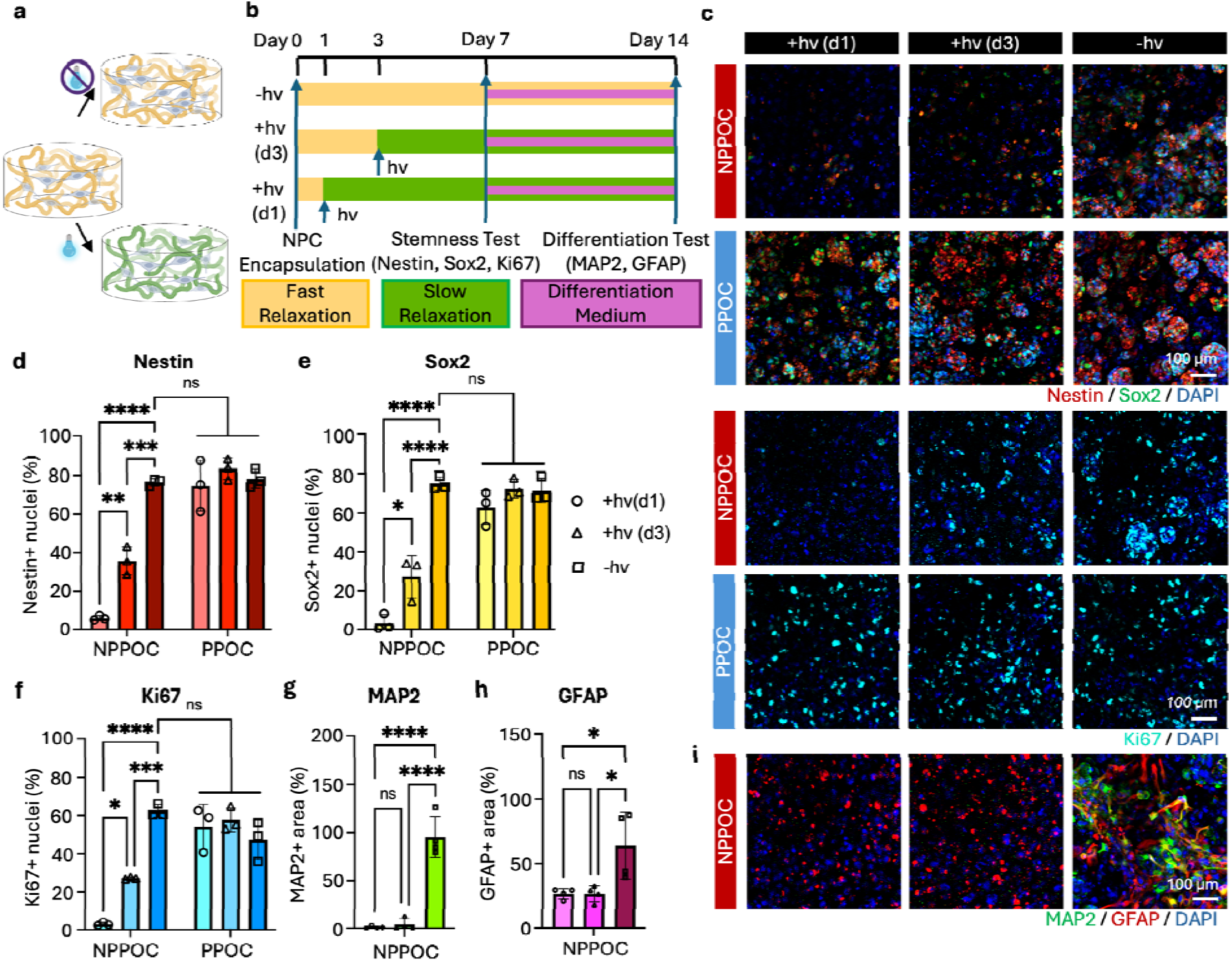
Temporal tuning of stress relaxation in 3D regulates human neural progenitor cell stemness. **(a)** Schematic illustrating NPCs encapsulated within hydrogels with phototunable stress relaxation. **(b)** Experimental timeline for evaluating the effects of temporally varied stress relaxation on NPC stemness (maintenance phase, days 0–7) and subsequent lineage commitment (differentiation phase, days 7–14). **(c)** Representative maximum-intensity projection confocal micrographs of stemness markers nestin (red) and Sox2 (green) and proliferation marker Ki67 (cyan) after 7 days of culture, counterstained with DAPI (blue). Quantification of the stemness markers **(d)** nestin and **(e)** Sox2 and the proliferation marker **(f)** Ki67 at day 7. Quantified expression of the differentiation markers **(g)** MAP2 and **(h)** GFAP from **(i)** confocal micrographs taken at day 14 of culture: MAP2 (green, neurons) and GFAP (red, astrocytes), counterstained with DAPI (blue). Data are presented as mean ± s.d. (n = 3; *p < 0.05, **p < 0.01, ***p < 0.001, ****p < 0.0001; two-way ANOVA with Tukey post-hoc test for (d-f); one-way ANOVA with Bonferroni post-hoc test for (g) and (h))

After 7 days of culture, clear differences in NPC phenotype emerged, with the duration of culture under rapidly relaxing conditions dictating the level of stemness maintenance. As expected, 7 days of culture in rapidly relaxing, unexposed NPPOC-protected hydrogels, as well as in photo-inert PPOC-protected controls, both with and without light exposure, resulted in robust maintenance of NPC stemness, as indicated by expression of the stemness markers nestin and Sox2 and the proliferation marker Ki67 (**Figure 4c-f**). In these conditions, the NPCs were able to self-organize into clusters and form cell-cell contacts. In contrast, transitioning the material to a slowly relaxing state 1 day post-encapsulation resulted in a marked loss of stemness. The NPCs largely remained as isolated single cells and lost expression of nestin, Sox2, and Ki67. Maintaining the NPCs in a rapidly relaxing environment for 3 days prior to irradiation resulted in an intermediate phenotype. Fewer and smaller cell clusters were observed, with overall lower fractions of cells expressing nestin, Sox2, and Ki67 compared to persistently rapidly relaxing controls. These results support the hypothesis that sufficient remodeling time is needed to form cell-cell contacts and maintain stemness.

As differentiation capacity is the ultimate functional readout of NPC stemness, the encapsulated cells were also subjected to conditions favoring differentiation (**Figure 4b**). Following 7 days of culture in maintenance medium, encapsulated NPCs were transitioned to differentiation medium lacking exogenous growth factors. Lineage commitment was evaluated by immunofluorescence staining for MAP2 (Microtubule-Associated Protein 2), which is enriched in the dendrites of mature neurons, and GFAP (Glial Fibrillary Acidic Protein), a canonical astrocyte marker (**Figure 4g-i**). Immunofluorescence analysis revealed robust neural differentiation only in the continuously rapidly relaxing condition, whereas slowing stress relaxation at either day 1 or day 3 post-encapsulation resulted in no detectable differentiation (**Figure 4g**). A similar effect was also observed for astrocyte differentiation (**Figure 4h**). Notably, although the condition irradiated at day 3 retained partial progenitor characteristics during the stemness maintenance period, this was insufficient to preserve subsequent differentiation potential. These results indicate that the transition to a slowly relaxing matrix impairs neuronal differentiation, consistent with previous reports demonstrating the necessity of matrix remodeling via stress relaxation for NPC stemness maintenance.[19], [20], [37] Together, these results demonstrate that the phototunable ELP-PEG platform enables dynamic regulation of matrix viscoelasticity in three-dimensional cultures, providing a tool to leverage temporally evolving mechanical cues to regulate cell fate.

### 2.6. Directing Neurite Outgrowth via Spatiotemporal Patterning of Stress Relaxation

Having demonstrated the ability to selectively modulate the viscoplastic remodelability of 3D hydrogel matrices by dynamically regulating stress relaxation behavior, we finally sought to demonstrate that the phototunable hydrogel system can regulate therapeutically meaningful cell behaviors in a spatiotemporally defined manner. Successful neural tissue engineering requires proper guidance of axons to their targets. As neurite extension in 3D is also enabled by viscoplastic remodeling,[23] we reasoned that by spatiotemporally restricting the rapidly relaxing regions of the hydrogels, we could direct neurite outgrowth. Motor neuron (MN) spheroids were selected as a model system, as regulating muscle reinnervation would be broadly useful in regenerative medicine.

Human induced pluripotent stem cell (hiPSC)-derived motor neuron spheroids[50] were encapsulated in the phototunable NPPOC-protected hydrogels and cultured in MN medium (**Figure 5a**). Spheroids were allowed to mature *in situ* until initial neurite extension was observed (typically 5 days), after which a subset of samples were irradiated with 365 nm light to induce the transition to a slowly relaxing network. All samples were cultured for a total duration of 14 days prior to fixation and immunofluorescence characterization (**Figure 5b**). Within the continuously rapidly relaxing networks, spheroids exhibited extensive multidirectional neurite extension, whereas transition to a slowly relaxing state significantly reduced neurite outgrowth, characterized by decreased total neurite length and neurite number (**Figures 5c,d**; **S21**). These results demonstrate that dynamic modulation of stress relaxation behavior can be used to regulate neurite outgrowth.

**Figure 5.**
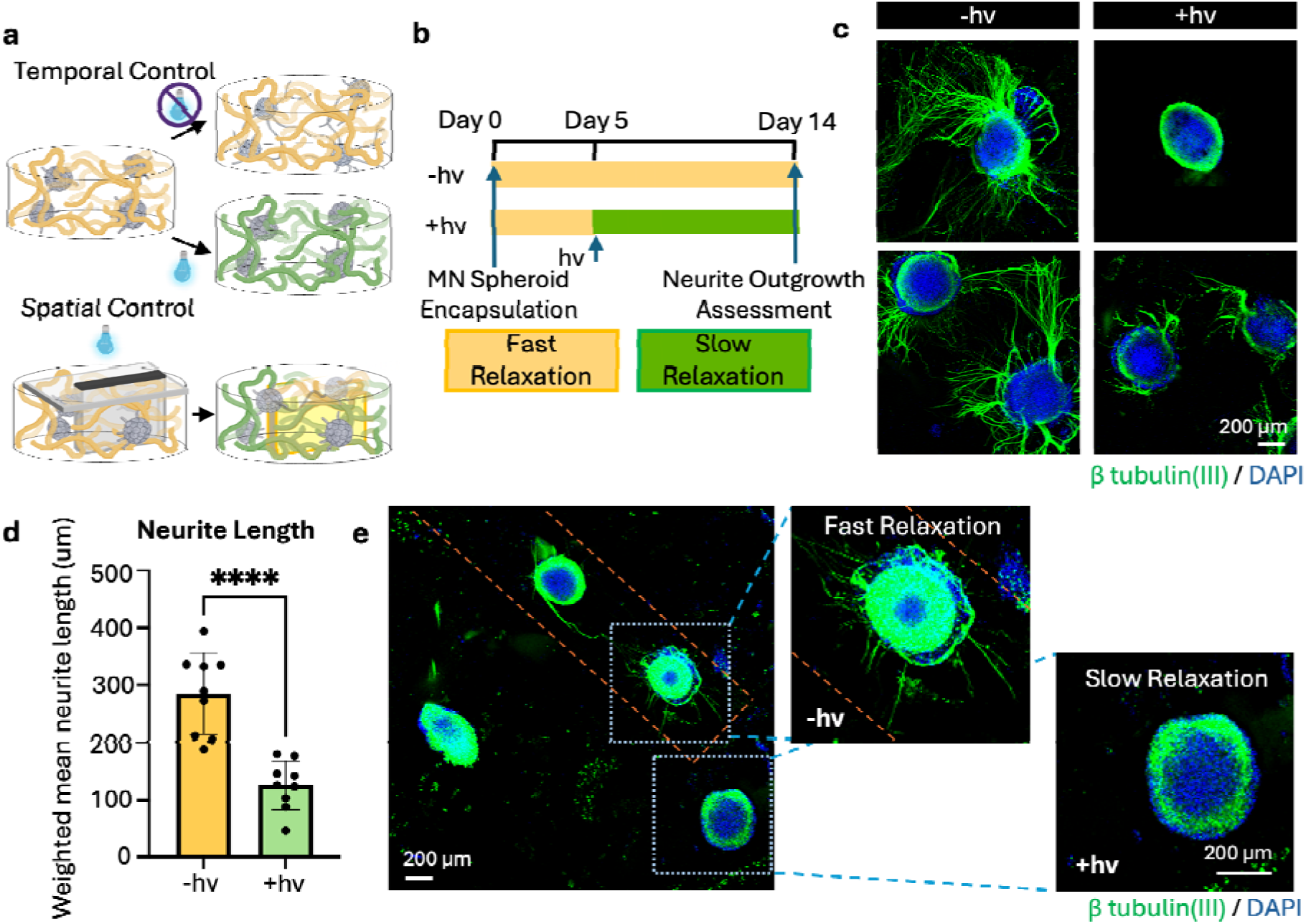
Spatiotemporal patterning of stress relaxation regulates neurite outgrowth from motor neuron spheroids. **(a)** Schematic illustration of hiPSC-derived MN spheroid 3D encapsulation and spatial photo-patterning approach. **(b)** Experimental timeline for evaluating the effects of temporally varied stress relaxation on MN spheroid neurite outgrowth. **(c, d)** Representative immunofluorescence maximum intensity projections and corresponding neurite length quantifications of MN spheroids after 14 days of culture. Spheroids within untreated hydrogels exhibit robust, isotropic neurite outgrowth, whereas irradiated samples exhibit significantly inhibited neurite outgrowth. n = 9, 3 individual hydrogel samples with total 9 spheroids for each condition were quantified; data are presented as mean ± s.d. (unpaired t-test, *p < 0.05, **p < 0.01, ***p < 0.001, ****p < 0.0001, **(e)** 3D viscoelastic photopatterning demonstrating spatial control over neurite outgrowth. Orange dashed lines denote boundary of light exposure.

As the direction of neurite outgrowth is a key determinant of innervation, we sought to exert spatial, in addition to temporal, control over viscoplastic remodelability in 3D to direct neurite outgrowth (**Figure 5a**). Spatial patterning was achieved using a custom photomask that shielded a narrow channel-like region of the hydrogel from irradiation, while surrounding regions were exposed to light and underwent transition to the slowly relaxing state (**Figure S22**). This generated localized regions with distinct relaxation behaviors within a continuous hydrogel, with the protected region remaining fast-relaxing and permissive for neurite outgrowth. Motor neuron spheroids positioned within the protected rapidly relaxing region exhibited neurite extension predominantly within the unexposed area, whereas spheroids located in the surrounding slow-relaxing matrix displayed limited neurite outgrowth (**Figure 5e**). These results demonstrate the feasibility of three-dimensional patterning of stress relaxation behavior for spatial regulation of neurite outgrowth.

## 3. Discussion

The ECM presents cells with heterogeneous mechanical environments that evolve during tissue development, aging and disease. [51], [52] Among the various mechanical properties sensed by cells, matrix viscoelasticity is increasingly appreciated as a key regulator of cell fate.[2], [3], [15], [22] Therefore, biomaterial platforms with an ability to modify viscoelastic mechanical properties after cells have established interactions with the matrix are required whether the goal is engineering tissue function or investigating cellular responses to dynamically varying biophysical cues in aging and disease. To achieve this on-demand control, we leveraged photo-triggered dynamic covalent crosslink exchange to modulate stress relaxation time while maintaining consistent stiffness. By permitting spatiotemporal modulation of stress relaxation over a physiologically relevant range, this hydrogel platform enables both fundamental studies of cell-ECM mechanobiology as well as dynamic engineering of cellular microenvironments to direct cell fate.

Our approach overcomes several limitations of prior systems that utilize photochemistry to tune relaxation rate. [24], [25], [26], [27] A key advantage is the use of bioselective hydrazone and oxime dynamic covalent crosslinks and cytocompatible photo-deprotection of alkoxyamines.[28], [29] In contrast to systems that turn viscous dissipation “on” or “off”[24], [26], the hydrazone to oxime transition spans a physiologically relevant range of stress relaxation times, from ∼10^2^ s to ∼10^4^ s. Furthermore, we also achieved graded tuning of network viscoelasticity by blending photoresponsive NPPOC and photoinert PPOC materials in defined ratios. This approach provides access to a continuous spectrum of viscoelastic states, offering opportunities to mimic the relaxation properties of different tissues, subtle biological gradients, or distinct stages of age-related changes in matrix mechanics. Beyond limitations in viscoelastic tunability exhibited by prior systems, reliance on radical-mediated reactions presents a particular concern for applications involving sensitive cell populations. Photoinitiator exposure alone, without irradiation, can cause substantial cell death,[53] and subsequent radical generation can cause off-target damage to cells and biomolecules, potentially compromising cell viability and complicating interpretation of cell responses to matrix mechanics [54], [55], [56], [57]. Our bioselective photocaged system enables post-gelation modulation of viscoelasticity without relying on free radicals or introducing additional reagents, avoiding potential sources of chemical interference with cell–matrix interactions.

The native ECM is protein-rich, and cell-ECM interactions rely on the binding of cell surface receptors to proteinaceous ECM components and active remodeling of the ECM by cell-secreted proteases. Thus, to mimic the complex 3D microenvironment of the native ECM, engineered biomaterials should also contain cell-interactive protein components. By engineering tunable viscoelasticity into recombinant ELP-based hydrogels, we have extended the ability to spatiotemporally pattern stress relaxation timescale to protein hydrogels. To incorporate the ELP component, we developed a robust strategy to incorporate multiple aldehyde moieties at selected locations in the protein to serve as hydrazine- and alkoxyamine-reactive sites. This approach is the inverse of prior strategies to form hydrazone-crosslinked ELP hydrogels, which pair hydrazine-functionalized proteins with aldehyde-bearing macromolecular crosslinkers.[23], [47], [49] Our strategy uses selective chemical approaches to overcome limitations of enzymatic or genetic code expansion-based approaches[40], [41] to install multiple aldehydes in a single protein, broadening the materials available for use in dynamic covalent hydrogels.

Fast-to-slow relaxation transitions elicited distinct cellular responses in 2D and 3D cultures. Slower relaxation promoted fibroblast elongation on 2D substrates, whereas in 3D matrices it restricted NPC clustering and motor neuron neurite extension. These findings suggest that the effects of matrix relaxation depend on cellular identity and culture context. Using a common ELP–PEG platform across these experiments reduces differences associated with comparing chemically distinct material systems and provides a basis for investigating this context dependence. The demonstrated compatibility with multiple cell types further supports the platform’s utility for studying diverse cellular responses to evolving matrix mechanics. Spatial control over cell behavior within a continuous hydrogel was demonstrated using patterned irradiation to direct fibroblast morphological transitions and neurite outgrowth. Beyond individual cell populations, spatial control of relaxation could enable the design of multicellular constructs with regions tailored to distinct mechanical requirements, directing cell-cell interactions and tissue organization.

We envision that this platform will expand the biomaterial toolkit for mechanobiology and tissue engineering, with the experiments presented here providing practical examples of its use across cell types and culture configurations. The capacity to modulate stress relaxation after initiating cell culture offers opportunities to investigate how evolving mechanical environments shape cellular behavior and to guide tissue development through spatially and temporally controlled matrix cues.

## 4. Experimental section

### 4.1. General

All chemicals were purchased from Fisher Scientific, Sigma-Aldrich, or BLDpharm, unless otherwise noted, and used as received. Poly(ethylene glycol) (PEG) precursors were purchased from JenKem Technology. Water-sensitive reactions were conducted using commercially available anhydrous solvents. Hydrazine- and alkoxyamine-functionalized multi-arm PEGs were synthesized following known routes, as described in the Supporting Information (**Figures S23-26**). PEG-aldehyde (8-arm, 20 kDa) was synthesized by Swern oxidation following a known procedure,[32] as previously reported by our lab (**Figure S27**).[37] ^1^H NMR spectra were recorded on Bruker NEO 400, 500 or 600 spectrometers at 400, 500 or 600 MHz, respectively. Chemical shifts were reported in parts per million (ppm) and were referenced to the residual solvent peak (CDCl_3_, δ^1^H = 7.26 ppm, DMSO-*d*_6_, δ^1^H = 2.50 ppm, and D₂O, δ ^1^H = 4.79 ppm). The following abbreviations were used s (singlet), d (doublet), dd (double-doublet), t (triplet), and m (multiplet). A collimated Thorlabs 365 nm mounted LED (M365L3) was used for irradiation, unless otherwise noted. Irradiance was measured using a UV-Meter Silver Line Full UV radiometer. Fluorescence microscopy was performed on a Zeiss LSM 900 laser scanning confocal microscope equipped with a broadband (385–730 nm) widefield epifluorescence light source and a Photometrics Prime BSI camera. No unexpected or unusually high safety hazards were encountered during these studies.

### 4.2. Synthesis of NPPOC-HNO-N_3_

2-((((2-(2-nitrophenyl)propoxy)carbonyl)amino)oxy)acetic acid (NPPOC-HNO-COOH) was synthesized as described in the Supporting Information, following a known synthetic route (**Figure S28-30**).[28] In a round bottom flask purged with argon, NPPOC-HNO-COOH (298 mg, 1 mmol, 1 eq), *N*-(3-dimethylaminopropyl)-*N*-ethylcarbodiimide (EDC, 201.8 mg, 1.3 mmol, 1.3 eq) and 4-dimethylaminopyridine (DMAP, 24.44 mg, 0.2 mmol, 0.2 eq) were dissolved in dichloromethane (DCM, 5 mL). 3-azidopropylamine (130 mg, 1.3 mmol,1.3 eq) was added to the solution, and the reaction was allowed to proceed overnight at room temperature. The solvent was removed under reduced pressure, and the residue was purified by silica gel chromatography (hexane: ethyl acetate = 1:4) to give the desired product as a light yellow oil (201 mg, 53% yield, **Figure S31**) ^1^H NMR (400 MHz, CDCl_3_) δ ppm 7.75 (dd, *J* = 8.1, 1.4 Hz, 1H), 7.58 (m, 1H), 7.42 (m, 2H), 4.36 (dd, *J* = 10.6, 6.0 Hz, 1H), 4.31 (s, 2H), 4.22 (dd, *J* = 10.5, 8.2 Hz, 1H), 3.75 (m, *J* = 13.8, 6.9 Hz, 1H), 3.38 (m, 4H), 1.83 (m, *J* = 6.8 Hz, 2H), 1.37 (d, *J* = 7.0 Hz, 3H).

### 4.3. Synthesis of PPOC-HNO-N_3_

The photo-inert PPOC-HNO-N_3_ precursor was prepared from 2-phenylpropan-1-ol following the same protocol as for NPPOC-HNO-N_3_, as described in the Supporting Information (125.7 mg, 50% yield, **Figures S32-34**). ^1^H NMR (400 MHz, CDCl_3_) δ ppm 8.05 (s, 1H), 7.64 (s, 1H), 7.33 (m, 2H), 7.25 (m, *J* = 5.9, 1.3 Hz, 2H), 7.22(s, 1H), 4.31 (s, 2H), 4.28 (dd, *J* = 7.2, 2.4 Hz, 2H), 3.38 (m, 4H), 3.11 (m, *J* = 7.1 Hz, 1H), 1.83 (m, *J* = 6.8 Hz, 2H), 1.30 (d, *J* = 7.1 Hz, 3H).

### 4.4. Synthesis of 8-arm 20 kDa PEG-hydrazine-alkoxyamine(NPPOC) (PEG-[8Hz/8HNO-NPPOC])

The synthesis of PEG-[8Hz/8HNO-NPPOC] was accomplished in 5 steps (**Supporting Figure S2**). First, the polymer end groups were functionalized with propargylglycine to present both amine and alkyne moieties. Boc-propargyl-Gly-OH (143 mg, 0.672 mmol, 4.2 eq to -NH_2_) and HATU (244 mg, 0.64 mmol, 4 eq) were dissolved in anhydrous *N,N-*dimethylformamide (DMF, 4 mL) in a two-neck round bottom flask. Upon full dissolution, *N,N*-diisopropylethylamine (DIEA, 280 μL, 1.6 mmol, 10 eq) was added to the solution, and the reaction was allowed to proceed for 10 min. Meanwhile, 8-arm PEG-amine (400 mg, MW ∼ 20 kDa, 0.02 mmol, 0.16 mmol -NH_2_, 1 eq) was dissolved in anhydrous DMF (4 mL) in a separate flask. The activated Boc-propargyl-Gly-OH solution was added dropwise to the stirring PEG solution and allowed to react overnight at room temperature under argon. The following day, the Boc-protected PEG product was precipitated by addition to 80 mL ice-cold diethyl ether. The collected crude PEG product was dissolved in DCM (4 mL). While stirring, triisopropylsilane (200 μL), water (200 μL), and trifluoroacetic acid (4 mL) were added to the Boc-protected PEG solution. The reaction was allowed to proceed for 4 h at room temperature. The deprotected PEG product was collected by precipitation in 80 mL ice-cold diethyl ether. The collected PEG product was dissolved in MilliQ water and dialyzed against MilliQ water for 2 days (MWCO 2 kDa, 4 x 4L, 4°C). After purification, the PEG solution was frozen and lyophilized to afford the product PEG-[8NH_2_/8alkyne] as a white solid (390 mg, 0.0195 mmol, 97.5% yield after 2 steps). NMR characterization is presented in the Supporting Information (**Figures S8, 9**).

Next, the hydrazine functionality was installed via amide coupling to the free amines. Tri-Boc-hydrazinoacetic acid (263 mg, 0.672 mmol, 4.2 eq to -NH_2_) was reacted with HATU (244 mg, 0.64 mmol, 4 eq) and DIEA (280 μL, 1.6 mmol, 10 eq) in anhydrous DMF (4 mL) for 10 min. The activated tri-Boc-hydrazinoacetic acid solution was added dropwise to pre-dissolved PEG-[8NH_2_/8alkyne] (400 mg, MW ∼ 20 kDa, 0.02 mmol, 0.16 mmol -NH_2_, 1 eq) in DMF (8 mL), and the reaction was allowed to proceed overnight at room temperature. The reaction mixture was added to ice-cold diethyl ether (80 mL) to precipitate the PEG. The precipitated PEG product was then dissolved in MilliQ water and dialyzed against MilliQ water (MWCO 2 kDa, 4 x 4L, 4°C), followed by lyophilization to obtain the PEG-[8Hz-Boc_3_/8alkyne] product as a white solid (383 mg, 0.0192 mmol, 95.8% yield). NMR characterization is presented in the Supporting Information (**Figure S10**).

A CuAAC approach was taken to add the NPPOC-protected alkoxyamine moieties. PEG-[8Hz-Boc_3_/8alkyne] (200 mg, MW ∼ 20 kDa, 0.01 mmol, 0.08 mmol -NH_2_, 1 eq) was dissolved in PBS (5 mL) and degassed with argon for 30 min. Copper (II) sulfate and ascorbic acid were dissolved in MilliQ water at concentrations of 12.8 mg/mL and 158.5 mg/mL, respectively, in separate vials. NPPOC-HNO-N_3_ (45.6 mg, 0.12mmol, 1.5 eq) was dissolved in a minimal amount of DMSO (∼100 μL) in another vial. Ascorbic acid (100 μL, 0.08 mmol, 1 eq), copper sulfate (100 μL, 0.008 mmol, 0.1 eq), and the entire NPPOC-HNO-N_3_ solution were added sequentially and slowly to the PEG solution. The reaction was allowed to proceed overnight at room temperature. The following day, the reaction solution was dialyzed against MilliQ water (MWCO 2 kDa, 4 x 4L, 4°C). The dialyzed solution was frozen and lyophilized to obtain the product, PEG-[8Hz-Boc_3_/8HNO-NPPOC], as a white solid (181 mg, 0.009 mmol, 90.5% yield). NMR characterization is presented in the Supporting Information (**Figure S11**).

In the final step, phosphoric acid was used for mild deprotection of the Boc groups to unmask the hydrazine moieties. After complete dissolution of PEG-[8Hz-Boc_3_/8HNO-NPPOC] (100 mg, MW ∼ 20 kDa, 0.005 mmol, 0.04 mmol -NH_2_, 1 eq) in DCM (3 mL), 100 μL of 85% phosphoric acid was added to the stirring PEG solution. The solution was vigorously stirred for 3 hours. The resulting PEG product separated from the solution and stuck to the vial. This sticky crude product was dissolved in MilliQ water and dialyzed against MilliQ water (MWCO 2 kDa, 4 x 4L, 4°C), followed by freezing and lyophilization to afford the final product, PEG-[8Hz/8HNO-NPPOC], as a white solid (70 mg, 0.0035 mmol, 70% yield, ∼60% overall yield after 5 steps). NMR characterization is presented in the Supporting Information (**Figure S12**).

### 4.5. Synthesis of 8-arm 20 kDa PEG-hydrazine-alkoxyamine(PPOC) (PEG-[8Hz/8HNO-PPOC])

The photo-inert PEG-[8Hz/8HNO-PPOC] precursor was prepared from PPOC-HNO-N_3_ following the same protocol as for PEG-[8Hz/8HNO-NPPOC], as described in the Supporting Information (176 mg, 59% overall yield after 5 steps). NMR characterization is presented in the Supporting Information (**Figures S14, 15**).

### 4.6. Expression and Characterization of ELPs

The ELPs used in this study were developed by Straley and Heilshorn[58] and recloned by our lab as previously described.[37], [59] Plasmids containing RGD-ELP were transformed into BL21(DE3)pLysS *Escherichia coli* (Thermo Fisher Scientific) and expressed under the control of the T7 promoter. Bacteria were cultured in Terrific Broth at 37 °C to an OD600 of 0.8, and ELP expression was induced by the addition of 1 mM isopropyl β-D-1-thiogalactopyranoside (IPTG). The culture temperature was reduced to 32 °C, and the protein was allowed to express for 7 h before collecting the bacteria by centrifugation. Bacterial pellets were resuspended in TEN buffer (10 mm Tris, 1 mm EDTA, and 100 mm NaCl, pH 8.0) and lysed using repetitive freeze-thaw cycles. DNAse I and phenylmethylsulphonyl fluoride (PMSF) were added once to the raw lysate to remove genomic DNA and inhibit proteases. The thawed lysates were adjusted to pH 9.0 using NaOH and purified through inverse temperature cycling at 37 °C and 4 °C.[60] The resulting ELP solution was desalted by dialyzing against MilliQ water (MWCO 2 kDa, 4 × 4.5 L, 4 °C). The dialyzed ELP solutions were frozen and lyophilized to obtain a white solid. Protein purity was confirmed post-expression by SDS-PAGE.

### 4.7. Synthesis of ELP-aldehyde (ELP-ALD)

CuAAC was used to functionalize ELPs with aldehyde moieties. Primary amines on the ELPs (14 in total) were first converted to azides via aqueous diazo transfer[61], as described in the Supporting Information (**Figure S16**). The aldehyde-containing linker, 2-(2-(2-(2-hydroxyethoxy)ethoxy)ethoxy) ethyl propiolate, was synthesized as described in the Supporting Information (**Figures S35, 36**). In a 25 mL round bottom flask, ELP-azide (50 mg, MW ∼ 38 kDa, 0.0013 mmol, 0.0182 mmol azide, 1 eq) was dissolved in 10 mL water. After complete dissolution, the solution was cooled to 0°C. 2-(2-(2-(2-hydroxyethoxy)ethoxy)ethoxy) ethyl propiolate (40 mg, 0.16 mmol, 9 eq to azide) was dissolved in 500 μL DMSO. Then, copper (II) sulfate and ascorbic acid were dissolved in MilliQ water at concentrations of 13.1 mg/mL and 288 mg/mL, respectively, in separate vials. Ascorbic acid (100 μL, 0.18 mmol, 10 eq to azide), copper sulfate (100 μL, 0.009 mmol, 0.5 eq to azide), and the entire 2-(2-(2-(2-hydroxyethoxy)ethoxy)ethoxy) ethyl propiolate solution were added sequentially and slowly to the ELP solution. The reaction was allowed to proceed overnight at room temperature. The following day, the reaction solution was dialyzed against MilliQ water (MWCO 2 kDa, 4 x 4 L, 4°C). The dialyzed solution was frozen and lyophilized to obtain the product, ELP-ALD, as a light yellow solid (45 mg, 0.0012 mmol, 90% yield). NMR characterization is presented in the Supporting Information (**Figures S17, 18**).

### 4.8. Hydrogel fabrication

To fabricate hydrogels, PEG precursor solutions were dissolved in PBS at room temperature for 5 min, and ELP precursor solutions were dissolved in PBS at 4 °C overnight. Both PEG and ELP solutions were subsequently maintained on ice until used. The concentrations of the precursor solutions were chosen to maintain a 1:1 molar ratio of reactive groups. Purely PEG-based hydrogels were prepared by mixing the two PEG precursor solutions in a 1:1 (v:v) ratio to achieve a final polymer content of 3.5% (w/v). ELP–PEG hydrogels were prepared by mixing the two precursor solutions in a 1:1 (v:v) ratio. The ELP-PEG hydrogels used for rheological characterization and 3D cell culture were comprised of final concentrations of 1.5% (w/v) ELP (38 kDa, 14 functional groups per molecule, fully functionalized based on NMR) and 1.95% (w/v) PEG (20 kDa, 8 functional groups per molecule if fully functionalized, 70% functionalized based on NMR). To achieve a suitable storage modulus to observe viscoelasticity-induced changes in cell morphology, the formulation for the 2D cell culture studies consisted of final polymer concentrations of 2.5% (w/v) ELP and 3.25% (w/v) PEG. To tune the extent of relaxation time tuning by blending NPPOC- and PPOC-protected PEGs, equal molar amounts of PEG-[8Hz/8HNO-NPPOC] and PEG-[8Hz/8HNO-PPOC] were pre-mixed before mixing with ELP solutions, resulting in a 2:1:1 volume ratio of ELP:PEG(NPPOC):PEG(PPOC).

### 4.9. Rheological Characterization

Rheological characterization of the purely PEG hydrogels for proof-of-principle was performed using a Rheometrics ARES-RFS3 strain-controlled rheometer. Rheological characterization of the photoreactive and photo-inert ELP-PEG hydrogel samples was performed on a stress-controlled rheometer (HR-30, TA Instruments) using an 8-mm sand-blasted parallel plate geometry, connected to a UV curing accessory for 365 nm light exposure. Polymer solutions were crosslinked *in situ* on the plate at room temperature with a solvent trap to prevent dehydration. For non-light exposed samples, the tests consisted of 1 hour time sweeps for gelation, frequency and strain sweeps to ensure operation within the linear viscoelastic regime, and a final stress relaxation test. All stress relaxation measurements were performed within the predetermined linear viscoelastic region (LVR) for each polymer network (**Figures S37-39**). For light-treated samples, the gels were exposed to 365 nm light at 12 mW/cm^2^ maximum intensity through the transparent bottom plate for 300 seconds after the 1 hour gelation time sweep for complete gelation. After light exposure, a 10-hour time sweep was added to allow crosslink exchange, followed by the frequency sweep, strain sweep, and stress relaxation tests. Time sweeps were performed at 3% strain and 1 Hz at room temperature. Oscillatory frequency sweeps were performed at 3% strain from 0.01 Hz to 100 Hz frequency. Strain sweeps were performed at 1 Hz frequency from 0.01% to 10% strain. Shear stress relaxation tests were performed by applying 3% strain over 5 seconds, and the stress decay was measured over time under constant strain. For samples relaxing too slowly for the relaxation half time to be directly measured, a double Maxwell model was fit to the data to calculate the relaxation half times (see Supporting Information and **Figure S40**).

### 4.10. 2D Fibroblast Culture and Analysis

Human fetal lung fibroblasts (MRC5) were purchased from the American Type Culture Collection (ATCC, CCL-171). MRC5s were cultured in high glucose DMEM supplemented with 10% fetal bovine serum (FBS), and 1% penicillin/streptomycin. To generate the stable reporter cell line, cells were lentivirally transduced to constitutively express the fluorescent actin reporter LifeAct-iRFP670 (Addgene plasmid #84385)[62]. Successfully transduced cells were enriched by fluorescence activated cell sorting (FACS). To fabricate hydrogels for 2D cell culture, 10 μL of premixed hydrogel solution (1:1 v/v PEG to ELP) was added to the bottom of a 24-well plate, and then a 12 mm circular cover class was added on the top of the hydrogel precursor solution droplet. The mixture was allowed to gel for 30 min at room temperature, resulting in a 12 mm circular gel with a thickness of ∼50 μm. PBS (1 mL) was then added into the well. For pre-treated gels, 365 nm light (50 mW/cm^2^, 90 seconds) was applied 1 hour after gelation. The plates containing hydrogels submerged in PBS was equilibrated in a tissue culture incubator overnight. Cells were seeded on the hydrogel substrates the following day at a density of 24,000 cells/well. Cells were imaged using the widefield mode on the Zeiss LSM 900 microscope setup equipped with an environmental control chamber (37 °C, 5% CO_2_). Live-cell time-lapse images were collected at fixed intervals (every half hour) for a total 48 hours. Cell morphology was quantified via CellProfiler. Extracted geometric features included major and minor axis length (to assess cell elongation) and form factor (to assess cell circularity).

### 4.11. 3D Neural Progenitor Cell Culture and Analysis

Human neural progenitor cells (ReNcell CX, SCC007) were purchased from Millipore Sigma. NPCs were cultured and expanded in poly-L-ornithine- and laminin-coated tissue culture flasks in maintenance medium (ReNcell medium supplemented with 20 ng/mL FGF-2 and 20 ng/mL EGF). Prior to encapsulation, NPCs were detached with Accutase, collected by centrifugation, and counted using a hemocytometer. The cells were subsequently resuspended in the PEG precursor solution at a density of 30,000 cells/μL. The ELP precursor solution was then added to the cell suspension, mixed by pipetting, and added to custom 4 mm diameter silicone molds plasma bonded to glass coverslips. The molds were transferred to 24 well plates and allowed to gel in a tissue culture incubator for 30 min before NPC maintenance medium was added. The resulting 3D hydrogel samples were ∼300 μm thick with cells distributed throughout. At 1 or 3 days post-encapsulation, 365 nm light (50 mW/cm^2^) was applied for 90 s to the respective samples. For NPC differentiation, after 7 d in the maintenance medium, the cells were cultured for an additional 7 d in the base medium without FGF-2 and EGF to induce differentiation. To quantify changes in cell fate, cells were immunostained with markers indicative of stemness maintenance and differentiation and imaged via confocal microscopy. Cells positive for nestin, Sox2 and Ki67 were manually counted in FIJI ImageJ software and reported as percentage of total cells counted based on DAPI signal. MAP2 and GFAP expression were reported as percent area positive for the respective marker, as calculated with FIJI ImageJ software.

### 4.12. 3D Culture and Analysis of Motor Neuron Spheroids

Motor neuron spheroids were derived from human iPSCs following published protocols[50], as described in the Supporting Information (**Figure S41**). Spheroids suspended in 6 μL of PBS were transferred to the cell culture molds described above. Then, 9 uL of premixed 5% (w/v) ELP and 6.5% (w/v) PEG solution was added to the mold and mixed gently with the PBS spheroid suspension to achieve the final hydrogel concentration of 1.5% (w/v) ELP and 1.95% (w/v) PEG. Two to five spheroids were encapsulated per hydrogel. Samples were allowed to gel in a tissue culture incubator for 30 min before motor neuron medium was added to the culture plate. The encapsulated spheroids were cultured for 5 days before exposure to 365nm light (50 mW/cm^2^, 90 seconds). For spatial photopatterning, custom quartz photomasks with a single channel-like opaque region were used, as detailed in the Supporting Information (**Figures S22, 42**). Neurite outgrowth from the spheroids was quantified by staining the samples for the neuronal marker β-tubulin III and analyzing the resulting confocal microscopy images using Sholl analysis via the Neuroanatomy plugin in FIJI ImageJ software,[63], [64] as detailed in the Supporting Information.

### 4.13. Cell Viability

Cell viability was assessed for 3D encapsulated NPCs. Cells were encapsulated and cultured as described above. For light-treated samples, 365 nm light (50 mW/cm^2^, 90 seconds) was applied 24 hours post-encapsulation. 24 hours later, the maintenance medium was changed and supplemented with 1 µM calcein-AM and 1 µM ethidium-homodimer. The samples were incubated at 37 °C for 45 min, washed twice with maintenance medium, and imaged via confocal microscopy. Cells positive for calcein-AM and ethidium-homodimer were manually counted in FIJI ImageJ software and reported as percentage of total cells.

### 4.14. Immunofluorescence Analysis

To assess cell fate outcomes by fluorescence microscopy, samples were fixed with 4% paraformaldehyde in PBS for 30 min at 37 °C, permeabilized with 0.25% Triton X-100 in PBS (PBST) for 1 hour, and blocked with 5% bovine serum albumin (BSA) and 5% goat serum (GS) in PBST for 3 hours at room temperature. Samples were subsequently stained with relevant antibodies (**Supporting Table S2**) in PBST containing 2.5% BSA and 2.5% GS on an orbital shaker at 4 °C overnight. The following day, the samples were washed three times with PBST, followed by incubation with corresponding secondary antibodies (**Supporting Table S3**) and DAPI (1:1000 dilution) on an orbital shaker at 4 °C overnight. Following 3 additional PBST washes at room temperature, the stained samples were imaged by confocal microscopy.

### 4.15. Statistical Analysis

Statistical analysis was performed using GraphPad Prism 10. Comparisons between two experimental groups were analyzed using a lognormal unpaired *t*-test. Comparisons among more than two experimental groups with a single varying parameter were performed using one-way analysis of variance (ANOVA) with Bonferroni *post hoc* testing. Comparisons among more than two experimental groups with two varying parameters were analyzed by a two-way ANOVA with Tukey *post hoc* testing. n.s. = not significant (*p* > 0.05), \**p <* 0.05, \*\**p <* 0.01, \*\*\**p <* 0.001, \*\*\*\**p <* 0.0001.

## Acknowledgements

The authors thank Prof. Kyle Vining (University of Pennsylvania) for use of the HR-30 rheometer and the radiometer, and Prof. Mark G. Allen (University of Pennsylvania) for the manufacturing of the photomask. The plasmid pLenti Lifeact-iRFP670 BlastR was a gift from Ghassan Mouneimne (via Addgene). The authors gratefully acknowledge the use of facilities and instrumentation supported by U.S. National Science Foundation (NSF) through the University of Pennsylvania Materials Research Science and Engineering Center (MRSEC) (DMR-2309043). This work was carried out in part at the Singh Center for Nanotechnology, which is supported by the NSF National Nanotechnology Coordinated Infrastructure Program under grant NNCI-2025608. T.Y.K. was supported by a Postdoctoral Fellowship Program (Nurturing Next-generation Researchers) granted by the National Research Foundation of Korea (RS-2024-00407821). Funding for this work was provided by the U.S. National Institutes of Health (NIH, R35 GM154913 to C.M.M.), a seed grant from the Center for Engineering Mechanobiology (CEMB), an NSF Science and Technology Center (CMMI 1548571), and the University of Pennsylvania Synergy Grant program.

## Conflicts of Interest

The authors declare no conflicts of interest.

## Data Availability Statement

All source data will be made publicly available at the time of publication.

## Notes

### Competing Interest Statement

The authors have declared no competing interest.

