## Supporting Information for "Directing Cell Fate via Spatiotemporally Patterned Viscoelasticity Driven by Dynamic Crosslink Exchange"

### Supporting Methods

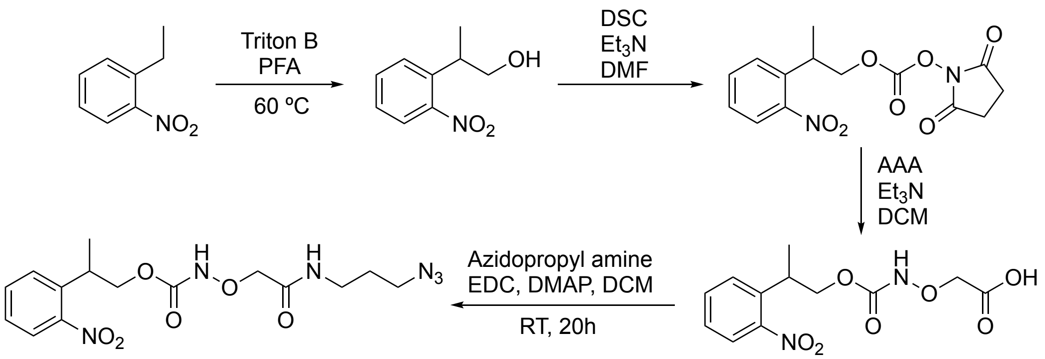

#### Synthesis of 2-(2-nitrophenyl)propan-1-ol

1-ethyl-2-nitrobenzene (18.48 g, 1 eq), paraformaldehyde (4.36 g, 1.19 eq), and a 40% w/w solution of benzyltrimethylammonium hydroxide in methanol (1 eq, 50 mL) were added to a round bottom flask and purged with argon. The reaction mixture was heated to reflux (~95 ℃) and allowed to react overnight. The solvent was removed under reduced pressure, and 200 mL 1M HCl solution were added with stirring. Ethyl acetate (200 mL) was added with vigorous stirring to partition the mixture. The organic phase was collected, and the aqueous phase was extracted with ethyl acetate (2x100 mL) The combined organic phases were dried over magnesium sulfate, filtered, and concentrated by rotary evaporation. The residue was purified by silica gel chromatography (hexane: ethyl acetate = 4:1 to 3:1) to afford the product as a red oil (11.76 g, 54.2% yield). Detailed NMR characterization is reported in **Figure S28**. $^1$H NMR (500 MHz, $CDCl_3$): δ ppm 7.74 (dd, J = 8.1, 1.2 Hz, 1H), 7.57 (m, 1H), 7.49 (dd, J = 7.9, 1.3 Hz, 1H), 7.36 (m,1H), 3.78 (m, 2H), 3.51 (m, J = 6.8 Hz, 1H), 2.05 (s, 1H), 1.33 (d, J = 7.1 Hz, 3H).

#### Synthesis of 2,5-dioxopyrrolidin-1-yl (2-(2-nitrophenyl)propyl) carbonate (NPPOC-OSu)

*N,N'*-disuccinimidyl carbonate (10.13 g, 1.5 eq) and 2-(2-nitrophenyl)propan-1-ol (4.78 g,1 eq) were added to a round bottom flask, purged with argon, and suspended in 125 mL anhydrous acetonitrile. $Et_3N$ (20.2 mL, 5.5 eq) was added dropwise, and the reaction mixture was stirred overnight at room temperature. The solvent was removed under reduced pressure, and the residue was purified by silica gel chromatography (hexane: ethyl acetate = 1:1) to give the desired product as a red oil (7.69 g, 90.5% yield). Detailed NMR characterization is reported in **Figure S29**. $^1$H NMR (500 MHz, CDCl3), δ ppm 7.81(dd, J = 8.1, 1.2 Hz, 1H), 7.6 (m, J = 7.6, 1.1 Hz, 1H), 7.49 (dd, J = 7.9, 1.3 Hz, 1H), 7.40 (m, 1H), 4.50, (m, 2H), 3.78 (m, J = 6.7 Hz, 1H), 2.79 (s, 4H), 1.42 (d, J = 7.1 Hz, 3H)

#### Synthesis of 2-((((2-(2-nitrophenyl)propoxy)carbonyl)amino)oxy)acetic acid (NPPOC-HNO-COOH)

NPPOC-OSu (3.42 g, 1 eq) and aminooxyacetic acid·1/2 HCl (1.16 g, 1 eq) were added to a round bottom flask, purged with argon, and suspended in 75 mL anhydrous dichloromethane (DCM). $Et_3N$ (7.5 mL, 5 eq) was added via syringe, and the reaction was allowed to proceed at room temperature, protected from light, for 20 h. The reaction mixture was concentrated by rotary evaporation, and the residue was taken up in 100 mL water. The aqueous phase was washed with ethyl acetate (2 x 100mL), acidified to pH ~ 1 with HCl, and then extracted with ethyl acetate (3 x 50 mL). The combined organic extracts were dried over $MgSO_4$, filtered, concentrated in vacuo, and purified via silica gel chromatography (ethyl acetate : acetic acid = 100:1). Fractions containing the desired product were concentrated in vacuo. To remove residual acetic acid, the residue was dissolved in dichloromethane (100 mL) and washed with water (3 x 100 mL) and brine (50 mL), dried over $MgSO_4$, filtered, and concentrated in vacuo to afford a yellow oil (1.94 g, 57% yield). Detailed NMR characterization is reported in **Figure S30**. $^1$H NMR (400 MHz, $CDCl_3$) δ ppm 9.87 (br. s, 1H), 8.30 (s, 1H), 7.75 (d, J = 8.1 Hz, 1H), 7.59 (t, J = 7.6 Hz, 1H), 7.47 (d, J = 7.6 Hz, 1H), 7.39 (t, J = 7.8 Hz, 1H), 4.42 (s, 2H), 4.37 (dd, J = 10.5, 6.1 Hz, 1H), 4.29 (dd, J = 10.5, 7.8 Hz, 1H), 3.75 (m, J = 6.9 Hz, 1H), 1.37 (d, J = 6.8 Hz, 3H)

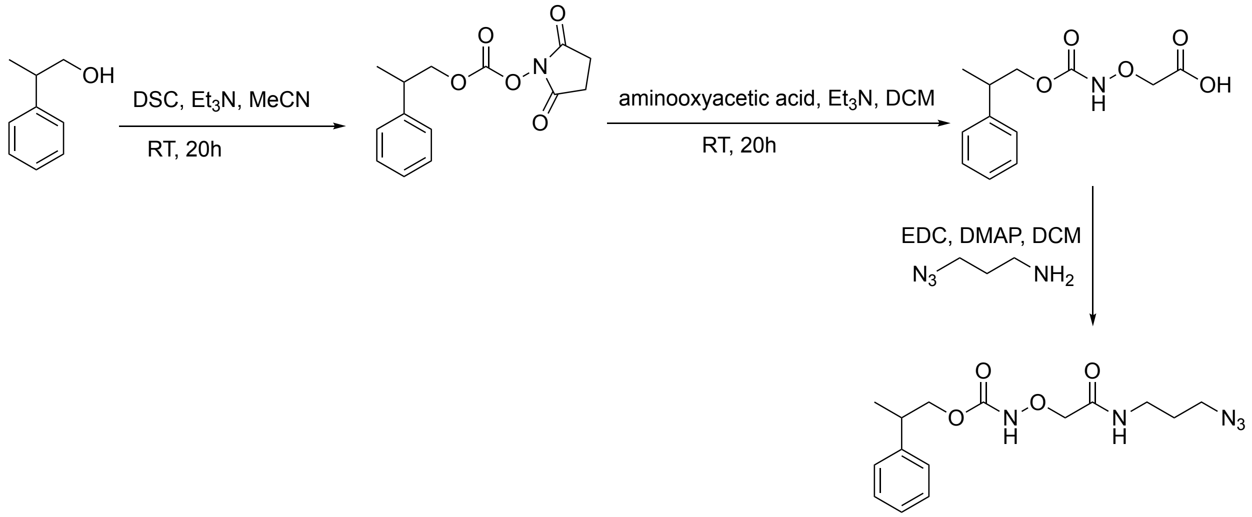

**Synthesis of 2,5-dioxopyrrolidin-1-yl (2-phenylpropyl) carbonate (PPOC-Osu)**

*N,N'*-disuccinimidyl carbonate (1.92 g, 1.5 eq) and 2-phenylpropan-1-ol (681 mg,1 eq) were added to a round bottom flask, purged with argon, and suspended in 125 mL anhydrous acetonitrile. $Et_3N$ (4 mL, 5.5 eq) was added dropwise, and the reaction mixture was stirred overnight at room temperature. The solvent was removed under reduced pressure, and the residue was purified by silica gel chromatography (hexane: ethyl acetate = 1:1) to give the desired product as a red oil (1.01 g, 73% yield). Detailed NMR characterization is reported in **Figure S32**. $^1$H NMR (400 MHz, $CDCl_3$) δ ppm 7.34 (m, *J* = 6.9, 1.0 Hz, 2H), 7.25 (m, 2H),7.22 (s, 1H), 4.38 (m, 2H), 3.21 (m, *J* = 7.1 Hz, 1H), 2.81 (s, 4H), 1.37 (d, *J* = 7.0 Hz, 3H).

**Synthesis of 2-((((2-phenylpropoxy)carbonyl)amino)oxy)acetic acid (PPOC-HNO-COOH)**

PPOC-OSu (556 mg, 1 eq) and aminooxyacetic acid·1/2 HCl (219 mg, 1 eq) were added to a round bottom flask, purged with argon, and suspended in 75 mL anhydrous dichloromethane. $Et_3N$ (1.5 mL, 5 eq) was added via syringe, and the reaction was allowed to proceed at room temperature for 20 h. The reaction mixture was concentrated by rotary evaporation, and the residue was taken up in 30 mL water. The aqueous phase was washed with ethyl acetate (2 x 30mL), acidified to pH ~ 1 with HCl, and then extracted with ethyl acetate (3 x 30 mL). The combined organic extracts were dried over $MgSO_4$, filtered, concentrated in vacuo, and purified via silica gel chromatography (ethyl acetate:acetic acid = 100:1). Fractions containing the desired product were concentrated in vacuo. To remove residual acetic acid, the residue was dissolved in dichloromethane (30 mL) and washed with water (3 x 30 mL) and brine (20 mL), dried over $MgSO_4$, filtered, and concentrated in vacuo to afford a yellow oil (248 mg, 49% yield). Detailed NMR characterization is reported in **Figure S33**. $^1$H NMR (400 MHz, $CDCl_3$) δ ppm 7.92 (s, 1H), 7.35 (m, 2H), 7.25 (m, *J* = 6.0, 1.3 Hz, 2H), 7.22 (s, 1H), 4.47 (s, 2H), 4.33 (m, 2H), 3.13 (m, *J* = 7.0 Hz, 1H), 1.31 (d, 3H).

#### Synthesis of 2-phenylpropyl (2-((3-azidopropyl)amino)-2-oxoethoxy)carbamate (PPOC-HNO-$N_3$)

PPOC-HNO-COOH (190 mg, 1 eq), *N*-(3-dimethylaminopropyl)-*N*-ethylcarbodiimide (EDC, 151.4 mg, 1.3 eq) and 4-dimethylaminopyridine (DMAP, 18.3 mg, 0.2 eq) were dissolved in dichloromethane (5 mL). 3-azidopropylamine (97.5 mg, 1.3 eq) was added to the solution, and the reaction was allowed to proceed overnight at room temperature. The solvent was removed under reduced pressure, and the residue was purified by silica gel chromatography (hexane: ethyl acetate = 1:4) to give the desired product as a light-yellow oil (125.7 mg, 50% yield). Detailed NMR characterization is reported in **Figure S34**. $^1$H NMR (400 MHz, $CDCl_3$) δ ppm 8.05 (s, 1H), 7.64 (s, 1H), 7.33 (m, 2H), 7.25 (m, *J* = 5.9, 1.3 Hz, 2H), 7.22(s, 1H), 4.31 (s, 2H), 4.28 (dd, *J* = 7.2, 2.4 Hz, 2H), 3.38 (m, 4H), 3.11 (m, *J* = 7.1 Hz, 1H), 1.83 (m, *J* = 6.8 Hz, 2H), 1.30 (d, *J* = 7.1 Hz, 3H).

#### Synthesis of 4-arm 20 kDa PEG-hydrazine-alkoxyamine(NPPOC) (PEG-[4Hz/4HNO-NPPOC])

The synthesis of PEG-[4Hz/4HNO-NPPOC] was accomplished in 5 steps following the same protocol as for PEG-[8Hz/8HNO-NPPOC] (**Figure S2**). 4-arm PEG-amine (200 mg, MW ~ 20 kDa, 0.01 mmol, 0.04 mmol -$NH_2$, 1 eq) was functionalized with propargylglycine via amide coupling with Boc-propargyl-Gly-OH (36 mg, 0.168 mmol, 4.2 eq to -$NH_2$), HATU (61 mg, 0.16 mmol, 4 eq) and *N,N*-diisopropylethylamine (DIEA, 70 μL, 0.4 mmol, 10 eq). After deprotection, purification and lyophilization, the product PEG-[4$NH_2$/4alkyne] was obtained as a white solid (189 mg, 0.0095 mmol, 94.5% yield after 2 steps). NMR characterization is presented in the Supporting Information (**Figures S3,4**).

Then, the free amines of PEG-[4$NH_2$/4alkyne] (120mg, MW ~ 20 kDa, 0.006 mmol, 0.024 mmol -$NH_2$, 1 eq) were modified with hydrazine groups by reacting with tri-Boc-hydrazinoacetic acid (9.6 mg, 0.0252 mmol, 4.2 eq to -$NH_2$), HATU (9.4 mg, 0.64 mmol, 4 eq) and DIEA (41.8 μL, 1.6 mmol, 10 eq) overnight. The PEG solution was dialyzed and subsequently lyophilized to obtain the PEG-[4Hz-$Boc_3$/4alkyne] product as a white solid (112 mg, 0.0056 mmol, 93% yield). NMR characterization is presented in the Supporting Information (**Figure S5**).

The NPPOC-protected alkoxyamine moieties were added via CuAAC. PEG-[4Hz-$Boc_3$/4alkyne] (112 mg, MW ~ 20 kDa, 0.0056 mmol, 0.0224 mmol -$NH_2$, 1 eq) dissolved in

PBS was reacted with ascorbic acid (4 mg in 100 μL water solution, 1 eq), copper sulfate (0.56 mg in 100 μL water solution, 0.00224 mmol, 0.1 eq) and NPPOC-HNO-$N_3$ (12.8 mg in 100 μL DMSO, 0.0336mmol, 1.5 eq). The reaction was allowed to proceed overnight at room temperature. The resulting solution was dialyzed against MilliQ water (MWCO 2 kDa, 4 x 4L, 4℃). The dialyzed solution was frozen and lyophilized to obtain the product, PEG-[4Hz-$Boc_3$/4HNO-NPPOC], as a white solid (105 mg, 0.00525 mmol, 94% yield). NMR characterization is presented in **Figure S6**.

To deprotect the Boc groups under mild conditions to selectively unmask the hydrazine moieties on PEG-[4Hz-$Boc_3$/4HNO-NPPOC] (100 mg, MW ~ 20 kDa, 0.005 mmol, 0.02 mmol -$NH_2$, 1 eq), 100 μL of 85% phosphoric acid was added to the stirring PEG solution in 3 mL DCM. After 3 hours, the product was dialyzed against MilliQ water (MWCO 2 kDa, 4 x 4L, 4℃) and lyophilized to obtain the final product, PEG-[4Hz/4HNO-NPPOC], as a white solid (63 mg, 0.0032 mmol, 63% yield, ~52% overall yield after 5 steps). NMR characterization is presented in **Figure S7**.

**Synthesis of 8-arm 20 kDa PEG-hydrazine-alkoxyamine(PPOC) (PEG-[8Hz/8HNO-PPOC])**

The synthesis of PEG-[8Hz/8HNO-PPOC] was accomplished in 5 steps following the same protocol as for PEG-[8Hz/8HNO-NPPOC] (**Figure S2**). The same CuAAC approach was taken to add the PPOC-protected alkoxyamine moieties. PEG-[8Hz-$Boc_3$/8alkyne] (300 mg, MW ~ 20 kDa, 0.015 mmol, 0.12 mmol -$NH_2$, 1 eq) was dissolved in PBS (5 mL) and degassed with argon for 30 min. Copper (II) sulfate and ascorbic acid were dissolved in MilliQ water at concentrations of 19.2 mg/mL and 237.8 mg/mL, respectively, in separate vials. PPOC-HNO-$N_3$ (61.5 mg, 0.18 mmol, 1.5 eq) was dissolved in a minimal amount of DMSO (~100 μL) in another vial. Ascorbic acid (100 μL, 0.08 mmol, 1 eq), copper sulfate (100 μL, 0.008 mmol, 0.1 eq), and the entire PPOC-HNO-$N_3$ solution were added sequentially and slowly to the PEG solution. The reaction was allowed to proceed overnight at room temperature. The following day, the reaction solution was dialyzed against MilliQ water (MWCO 2 kDa, 4 x 4L, 4℃). The dialyzed solution was frozen and lyophilized to obtain the product, PEG-[8Hz-$Boc_3$/8HNO-PPOC], as a white solid (267 mg, 0.0089 mmol, 89% yield). NMR characterization is presented in **Figure S14**.

To deprotect the Boc groups to unmask the hydrazine moieties, phosphoric acid was used. After complete dissolution of PEG-[8Hz-$Boc_3$/8HNO-PPOC] (250 mg, MW ~ 20 kDa, 0.0125 mmol, 0.1 mmol -$NH_2$, 1 eq) in dichloromethane (4 mL), 200 μL of 85% phosphoric acid was added to the stirring PEG solution. The solution was vigorously stirred for 3 hours. The resulting PEG product separated from the solution and stuck to the vial. This sticky crude product was dissolved in MilliQ water and dialyzed against MilliQ water (MWCO 2 kDa, 4 x 4L, 4℃), followed by freezing and lyophilization to afford the final product, PEG-[8Hz/8HNO-PPOC], as a white solid (176 mg, 0.0088 mmol, 88% yield, 59% overall yield after 5 steps). NMR characterization is presented in **Figure S15**.

**Synthesis of ELP-azide**

Azide functional groups were introduced into ELPs via aqueous diazo transfer.[1] 100 mg ELP was dissolved in 4 mL freshly prepared potassium carbonate buffer (10 mg/mL in MilliQ water). 400 μL copper(II) chloride solution (1 mg/mL) was added to the ELP solution, followed by the addition of sufficient 1H-imidazole-1-sulfonyl azide tetrafluoroborate, dissolved in potassium carbonate buffer, to modify all primary amines (14 per ELP). The reaction was allowed to proceed for 24 h at room temperature with constant stirring. The solution was dialyzed against MilliQ water (MWCO 2 kDa, 4 x 4L, 4℃), followed by freezing and lyophilization to afford the final product, ELP-azide, as a light-yellow powder. NMR characterization is presented in **Figure S17**. An alternative approach to synthesize ELP-azide was via amide coupling reactions with azidoacetic acid-NHS ester [2].100 mg ELP was dissolved in 5 mL DMSO. Then, 9.5 mg azidoacetic acid-NHS ester (0.048 mmol, 1.3 eq) was added to the ELP solution, followed by the addition of triethylamine (10 µL, 0.072 mmol, 1.5 eq to amine group). The reaction mixture was stirred at room temperature overnight, followed by dialysis and lyophilization as above to afford the final product ELP-azide. Both approaches were successful, and diazo transfer was selected as the default protocol because the reaction occurs in aqueous solution.

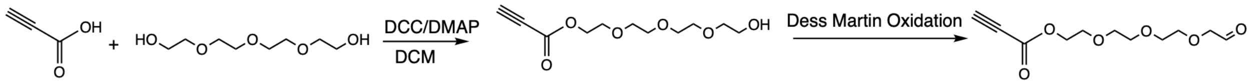

**Synthesis of 2-(2-(2-(2-hydroxyethoxy)ethoxy)ethoxy)ethyl propiolate**

In a round bottom flask, tetraethlyene glycol (TEG, 1 g, 5.15 mmol, 1.1 eq), *N,N'*-dicyclohexylcarbodiimide (966 mg, 4.68 mmol, 1 eq) and 4-(dimethylamino)pyridine (57.18mg, 0.468mmol, 0.1 eq) were dissolved in 10 mL DCM. After complete dissolution, the solution was cooled to 0° C. Then, a solution of propiolic acid (327.87 mg, 4.68 mmol, 1 eq) in 10 mL DCM was added dropwise to the TEG solution over 30 min. The reaction was allowed to proceed for 2 hours on an ice bath before being moved to room temperature for another 5 hours. To purify the crude product, the solution was first filtered through celite and concentrated by rotary evaporation. The residue was purified by silica gel chromatography (DCM : MeOH = 93 : 7) to afford the product, 2-(2-(2-(2-hydroxyethoxy)ethoxy)ethoxy)ethyl propiolate, as a ruby oil (380 mg, 1.54mmol, 30% yield) Detailed NMR characterization is reported in **Figure S35**. $^{1}H$ NMR (400 MHz, $CDCl_3$) δ ppm 4.36 (m, 2H), 3.73 (m, 4H), 3.67 (d, *J* = 2.7 Hz, 8H), 3.62 (m, 2H), 2.90 (s, 1H).

**Synthesis of 2-(2-(2-(2-oxoethoxy)ethoxy)ethoxy)ethyl propiolate**

2-(2-(2-(2-Hydroxyethoxy)ethoxy)ethoxy)ethyl propiolate (100 mg, 0.41 mmol, 1 eq) was dissolved in 5 mL DCM. Dess Martin periodinane (DMP, 350 mg, 0.82 mmol, 2 eq) was dissolved in 10 mL DCM in a separate vial. The DMP solution was added to the linker precursor solution, and the reaction was allowed to proceed overnight, vented to the atmosphere. The solution turned cloudy overnight and formed a white precipitate. To purify the crude product, 10 mL saturated $NaHCO_3$ solution was added to the reaction mixture and vigorously stirred for 5 min to quench the unreacted DMP and byproducts. Then, the organic phase was separated, washed with saturated $NaHCO_3$ solution and brine, and dried over magnesium sulfate. After filtration and concentration by rotary evaporation, the product was obtained as a white solid (58 mg, 0.24 mmol, 58% yield). Detailed NMR characterization is reported in **Figure S36**. $^1H$ NMR (400 MHz, $CDCl_3$) δ ppm 9.74 (s, 1H), 4.36 (m, 2H), 4.17 (d, *J* = 0.8 Hz, 2H), 3.73 (m, *J* = 8.0, 4.2, 2.4 Hz, 6H), 3.67 (s, 4H), 2.90 (s, 1H).

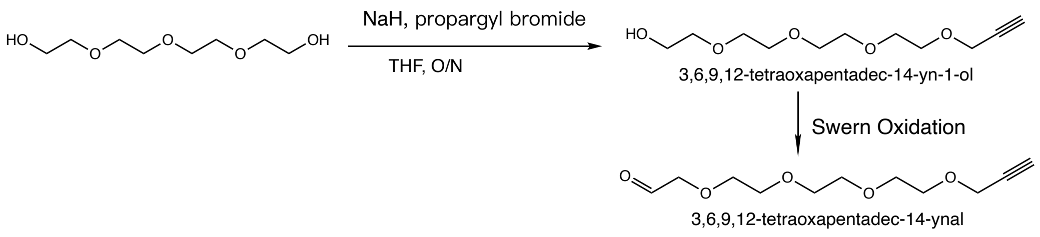

**Synthesis of 3,6,9,12-tetraoxapentadec-14-yn-1-ol**

3,6,9,12-tetraoxapentadec-14-yn-1-ol was synthesized following a published protocol[3]. In a round bottom flask, tetraethlyene glycol (TEG, 4.5 mL, 1.9 eq) was dissolved in 20 mL THF, and then the solution was cooled to 0°C in an ice bath. Sodium hydride (351 mg, 1.1 eq) was added portionwise, and the stirring solution was maintained at room temperature for 1 hour before being re-cooled to 0°C. Propargyl bromide (1 mL, 1 eq) was added dropwise by syringe pump over 40 min. The reaction was allowed to proceed overnight at room temperature. To quench the reaction, 10 mL cold water was added, and the mixture was extracted with DCM (3 x 30 mL), washed with brine (20 mL), and dried over magnesium sulfate. After filtration, the crude product was concentrated by rotary evaporation and purified by silica gel chromatography (DCM : MeOH = 94 : 6). The product was obtained as a colorless oil (1.8 g, 48% yield). Detailed NMR characterization is reported in **Figure S43**. $^1H$ NMR (400 MHz, $CDCl_3$) δ ppm 4.20 (d, J = 2.4 Hz, 1H), 3.61-3.72 (m, 16H), 2.42 (t, J = 2.4 Hz, 1H).

**Synthesis of 3,6,9,12-tetraoxapentadec-14-ynal**

3,6,9,12-tetraoxapentadec-14-ynal was synthesized following a published protocol[4]. A solution of anhydrous DMSO (600 μL, 4 eq) and DCM (10 mL) was cooled to -70°C and oxalyl chloride (375 μL, 2 eq) was added dropwise. After 15 min, a solution of 3,6,9,12-tetraoxapentadec-14-yn-1-ol (500 mg, 1 eq) in DCM (8 mL) was added slowly. After 30 min, DIEA (3.75 mL, 10 eq) was added. The reaction was allowed to proceed for 3 hours, with the mixture gradually warming to room temperature from -70°C. The solution was diluted with 50 mL DCM and washed with 1M HCl (2 x 30 mL) and water (30 mL). The organic layer was dried and concentrated. The crude product was purified by silica gel chromatography (hexane : acetone = 7:3). The product was obtained as a colorless oil (58 mg, 11% yield). Detailed NMR characterization is reported in **Figure S44**. $^1H$ NMR (400 MHz, $CDCl_3$) δ ppm 9.05 (s, 1H), 4.21 (d, J = 2.4 Hz, 1H), 3.64-3.74 (m, 14H), 2.43 (t, J = 2.4 Hz, 1H).

**Unsuccessful synthesis of ELP-ALD from 3,6,9,12-tetraoxapentadec-14-ynal**

We additionally attempted to aldehyde functionalize ELP-azide using the propargyl ether linker, following a similar CuAAC synthetic route. ELP-azide (50 mg, 0.0013 mmol, 0.0182 mmol azide, 1 eq) was dissolved in 10 mL water. After complete dissolution, the solution was cooled to 0°C. Ascorbic acid (28.8 mg in 100 μL MilliQ water, 0.18 mmol, 10 eq to azide), copper sulfate (1.31 mg in 100 μL MilliQ water, 0.009 mmol, 0.5 eq to azide), and the 3,6,9,12-tetraoxapentadec-14-ynal solution (45 mg in 300 μL DMSO) were added sequentially and slowly to the ELP solution. The reaction was allowed to proceed overnight at room temperature. The following day, the reaction solution was dialyzed against MilliQ water (MWCO 2 kDa, 4 x 4 L, 4℃), followed by freezing and lyophilization. From $^{1}H$ NMR characterization, there was no characteristic peak located at a chemical shift of 8-9 ppm (δ). Furthermore, no gelation was observed upon mixing solutions of 3.5% (w/v) PEG-hydrazine and 3% (w/v) of this product. Together, these results indicated unsuccessful aldehyde modification to ELP from the propargyl ether linker, motivating continued use of the more reactive propiolate linker.

**Synthesis of 8 arm 20kDa PEG-hydrazine**

PEG-hydrazine was synthesized by amide coupling as previously reported.[5] In a round bottom flask purged with argon, 8-arm 20 kDa PEG-amine (100 mg, 0.005 mmol) was dissolved in 3 mL anhydrous DMF. In a separate vial, tri-Boc-hydrazinoacetic acid (35.6 mg, 0.168 mmol, 4.2 equiv. per amine) was dissolved in anhydrous DMF (~3 mL) and activated with HATU (60.8 mg, 0.16 mmol, 4.0 equiv. per amine) and DIEA (69.7 uL, 0.4 mmol, 10 equiv. per amine). The tri-Boc-hydrazinoacetic acid reaction mixture was stirred for 10 minutes and then added to the PEG solution. The reaction was allowed to proceed overnight at room temperature. The product was precipitated in ice-cold diethyl ether, dried, and treated with a solution of 1:1 DCM:TFA (4 mL), water (100 μL) and triisopropylsilane (100 μL) for 4 hours to remove the Boc groups. The resulting compound was precipitated in ice-cold diethyl ether, dissolved in MilliQ water, dialyzed against MilliQ water (MWCO 2 kDa, 4 x 4L, 4 °C), and lyophilized to obtain the desired product as a white powder (98 mg, 98% yield). NMR characterization is presented in **Figures S23, 24**.

**Synthesis of 8 arm 20kDa PEG-alkoxyamine**

PEG-alkoxyamine was synthesized following a known synthetic route.[6] In a round bottom flask purged with argon, triphenylphosphine (42 mg, 4 eq), *N*-hydroxyphthalimide (13 mg, 2 eq) and 8 arm 20kDa PEG-OH (100 mg, 1 eq) was dissolved in acetonitrile (3 mL) and THF (0.5 mL). Then, the solution was cooled to 0° C on an ice bath. Diisopropyl azodicarboxylate (24.3 mg, 3 eq), dissolved in THF (0.5 mL), was added dropwise to the reaction mixture. The reaction was allowed to react overnight, with the temperature naturally raised from 0° C to room temperature. The product was precipitated in ice-cold diethyl ether, and the white precipitate was dissolved in a minimal amount of DCM (~1 mL). 100 uL anhydrous hydrazine was added with vigorous stirring to remove the protecting group. The resulting deprotected compound was precipitated in ice-cold diethyl ether, dissolved in MilliQ water, dialyzed against MilliQ water (MWCO 2 kDa, 4 x 4L, 4 °C ), and lyophilized to obtain the desired product as white powder (70 mg, 70% yield). NMR characterization is presented in **Figures S25, 26**.

**Hydrogel Stability Assessment**

Hydrogel precursor solutions comprising 3.9% 8-arm PEG-[8Hz/8HNO-NPPOC] and 3%ELP-ALD were prepared as described in the Hydrogel Fabrication section. To assess hydrogel stability, 40 uL hydrogels (1:1 (v:v) PEG:ELP) were cast in pre-weighed vials and kept at room temperature for 30 min to ensure complete gelation. PBS (1 mL) was added to the hydrogel-containing vials, and the samples were maintained at 37 °C for 1, 3, 7 and 14 days. The initial mass, wet mass, and dry mass of the hydrogels were measured. The wet mass was obtained by weighing the hydrogel after removing the PBS. After measuring the wet mass, the samples were frozen and lyophilized to obatain the dry mass. Hydrogel mass swelling ratios were calculated by dividing the wet mass by the dry mass of the samples (**Figure S20**).

**Stress relaxation curve fitting**

For the light-exposed, NPPOC-protected samples, the relaxation times exceeded the measurable range of the rheometer, necessitating an approximate fitting method to quantify the stress relaxation behavior, i.e. the stress relaxation half time, of the hydrogels. We found that fitting the data using the classic Maxwell model ($\sigma(t) = \sigma_0 e^{-t/\tau}$) oversimplifies the behavior of the ELP-PEG network, which involves contributions from both polymer chain physical entanglements and the dynamic chemical crosslinks. To accurately represent the dual relaxation mechanisms, a second-order exponential decay (double Maxwell model) fit was employed ($\sigma(t) = Ae^{-t/a} + Be^{-t/b}$ (A, B> 0)). As demonstrated in **Figure S40**, this dual relaxation mode approach fits all measured conditions well.

**Predicting and validating the kinetics of hydrazone-to-oxime crosslink exchange**

The transition from the fast-relaxing (hydrazone) state to the slow-relaxing (oxime) state is governed by three sequential processes: NPPOC uncaging, hydrazone hydrolysis (the reverse reaction), and subsequent oxime formation.

The kinetics of NPPOC uncaging are highly dependent on UV intensity and exposure duration. Based on established literature, a 300 second exposure at 20 mW/cm$^2$ was sufficient to achieve complete alkoxyamine uncaging.[7] We confirmed this timescale through UV-Vis spectroscopic analysis of the photolysis kinetics, finding that uncaging is achieved within around 90 seconds at an intensity of 50 mW/cm$^2$ (**Figure S45**). Thus, NPPOC uncaging proceeds rapidly and is unlikely to be the limiting step of the crosslink exchange mechanism.

Once the alkoxyamines are uncaged, bond exchange is governed by two chemical equilibria:

$$hydrazine + aldehyde \underset{k_{r1}}{\overset{k_{f1}}{\rightleftharpoons}} hydrazone$$

$$alkoxyamine + aldehyde \underset{k_{r2}}{\overset{k_{f2}}{\rightleftharpoons}} oxime$$

If we define A = hydrazine, B = aldehyde, C = hydrazone, D = alkoxyamine, and E = oxime, where $k_{f1}$ and $k_{r1}$, $k_{f2}$ and $k_{r2}$ are the forward rate and reverse rate for the hydrazone and oxime reactions, respectively, the system of reactions can be written as:

$$A + B \underset{k_{r1}}{\overset{k_{f1}}{\rightleftharpoons}} C$$

$$D + B \underset{k_{r2}}{\overset{k_{f2}}{\rightleftharpoons}} E$$

The corresponding rate equations are:

$$\frac{d[A]}{dt} = -k_{f1}[A][B] + k_{r1}[C]$$

$$\frac{d[D]}{dt} = -k_{f2}[D][B] + k_{r2}[E]$$

$$\frac{d[B]}{dt} = -k_{f1}[A][B] + k_{r1}[C] - k_{f2}[D][B] + k_{r2}[E]$$

$$\frac{d[C]}{dt} = +k_{f1}[A][B] - k_{r1}[C]$$

$$\frac{d[E]}{dt} = +k_{f2}[D][B] - k_{r2}[E]$$

Reported rate constants for hydrazone and oxime crosslinks vary by several orders of magnitude depending on the reaction conditions and characterization method (**Supporting Table S1**).

Campagna et al.[8] recently reported a study of bonding and exchanging kinetics in a hydrazone and oxime-based PEG hydrogel system, which is the most similar reported configuration to the present system. We thus employed their measured rate constants to solve the rate laws and estimate the timescale of bond exchange kinetics.

In hydrogels prepared from with 3% (w/v) ELP-aldehyde stock solution, the final aldehyde concentration in the gel is:

$$\left(\frac{1}{2}\right)\left(\frac{30\ mg}{mL}\right)\left(\frac{1\ mmol}{38000\ mg}\right)\left(\frac{14\ Aldehyde}{1\ ELP}\right) = 5.5\ mM$$

For a hydrazone-based hydrogel at equilibrium, the active hydrazone bonds can be calculated through thermodynamic constants.[9]

$$[Hydrazine]_{eq} + [Aldehyde]_{eq} \rightleftharpoons [Hydrazone]_{eq}$$

With $K_{eq,hydrazone}$ = 740 $M^{-1}$, and $[Hydrazone]_{eq}$ = x, the active bond concentration is:

$$K_{eq,Ho} = \frac{[Hydrazone]_{eq}}{[Hydrazine]_{eq}[Aldehyde]_{eq}} = \frac{x}{(0.0055 - x)^2} = 740\ M^{-1}$$

$$x = [Hydrazone]_{eq} = 3.367\ mM$$

The hydrazone-to-oxime bond exchange happens strictly via the hydrolytic dissociative pathway. A second-order irreversible analytical approximation is used to estimate the exchange rate constant, assuming an oxime formation rate constant $k_2 = 0.12\ M^{-1}s^{-1}$.

$$\frac{d[Oxime]}{dt} = -\frac{d[Aldehyde]}{dt} = k_{obs,exhange}\ [Aldehyde]^2$$

$$k_{obs,exhange} \approx k_2\left(\frac{[Aldehyde]_{eq}}{[Hydrazone]_0 + [Hydrazone]_{eq}}\right) = 0.12\mathrm{M}^{-1}\mathrm{s}^{-1}\left(\frac{0.0021}{0.0055}\right) = 0.045\ M^{-1}s^{-1}$$

For 90% hydrazone-to-oxime exchange, the estimated time is thus:

$$\frac{1}{[Aldehyde]_t} - \frac{1}{[Aldehyde]_0} = k_{obs,exhange} \cdot t$$

$$\frac{1}{0.00055} - \frac{1}{0.0055} = 0.045 \cdot t$$

$$t \approx 36363\ seconds \approx 10\ hours$$

Therefore, it should take around 10 hours for 90% of the initial hydrazone crosslinks to be converted to oximes. This is consistent with numerical simulations of the system of ODEs (**Figure S1a**). However, the crosslink exchange time is quite sensitive to the rate constants used. For example, using other literature values can result in exchange timescales on the order of days (**Figure S1b**).

Given the broad potential range of exchange timescales, we sought to confirm the timescale of bond exchange in our hydrogel system via rheological characterization. Consistent with the conversion from rapidly relaxing hydrazone to slowly relaxing oxime crosslinks, the loss modulus, and thus tan(δ), decrease over time after light exposure and reach a plateau after ~10 hours (**Figure S1c, d**).

**Differentiation of iPSCs into motor neuron progenitors**

The hiPS cell line WTC11 (Coriell Institute, GM25256) was maintained in feeder-free culture on 6-well cell culture plates coated with Matrigel (Corning) using mTeSR+ medium. Cells were passaged every 3 to 5 days using Versene (gentle cell dissociation) for 5 minutes at 37°C.

Human induced pluripotent stem cells (hiPSCs) were differentiated into motor neuron (MN)-like lineages for spheroid generation, as previously described[10]. On day 0, hiPSC clusters were dissociated into single cells using Accutase for 5 minutes at 37°C. The cells were then reseeded onto Matrigel-coated 6-well plates and cultured in neuronal induction medium supplemented with 10 μM ROCK inhibitor (Y-27632). After 24 hours, the medium was replaced with neuronal induction medium without ROCK inhibitor, and the medium was refreshed every other day until day 6 to induce differentiation into neuroepithelial progenitors (NEPs). On day 6, the cells were passaged using GCDR, replated onto laminin/poly-L-ornithine-coated 6-well plates, and cultured in motor neuron progenitor (MNP) patterning medium for an additional 6 days. The resulting MNPs could be efficiently expanded for up to four passages and were maintained in MNP expansion medium for at least one passage before spheroid generation. All media formulations were prepared using a 1:1 mixture of Neurobasal medium and DMEM/F12 supplemented with GlutaMAX, with detailed compositions provided in **Supporting Table S4**.

**MN spheroid generation**

MNPs were dissociated into single cells using Accutase (5 min, 37°C) and were resuspended in MN induction and maturation medium (1 mL). Cells were counted using a hemocytometer, and 30,000 MNPs were plated in a well of a 96 U-bottom ultra-low attachment plate in 150 µL of MN induction and maturation medium. Plates were centrifuged at 1200 rpm for 10 minutes and were incubated at 37°C overnight.

**Quantification of MN spheroid neurite outgrowth in hydrogels**

Extracted data included radius of every concentric ring ($R_i$), numbers of intersections (counting times of axon crossed the specific radial ring, $N_i$), estimated length of all neurites in the shell of adjacent rings ($L_i$). To visualize the data in spheroid-based, several factors were calculated as follows. The radius of the spheroid ($R_s$) was set manually.

Weighted mean axon length: $R_{weighted} = \frac{\sum(R_i - R_s)N_i}{\sum N_i}$

Total cumulative length: $L_{tot} = \sum L_i$

Normalized average length: $L_{avg} = \frac{\sum L_i}{N_{max}}$

**Spatially patterning viscoelasticity to guide neurite outgrowth**

To spatially control light exposure and guide neurite outgrowth, a photomask was fabricated using a conventional photolithographic process (**Figure S22**). A 500 µm-thick quartz substrate was deposited with 100 nm of Cr by PVD evaporation, coated with AZ4620 photoresist, and patterned to define a central opaque feature consisting of a rectangular strip terminated by a semicircular cap 650 µm in diameter. The patterned photoresist served as the mask for Cr etching and was subsequently stripped. The completed photomask was attached directly beneath a glass-bottom well plate, blocking light in the Cr-patterned region while transmitting it through the surrounding transparent area to expose the hydrogel. The masking performance in the presence of the separation introduced by the well-plate glass was evaluated using the setup illustrated in Figure S39. A 15 mm-diameter, 250 µm-thick quartz disk coated with a 10 µm-thick AZ4620 layer was placed inside the well with the photoresist facing upward as a UV-sensitive surrogate sample. The assembly was illuminated from below in a dark enclosure using 365 nm UV light at a dose of 50 mW/cm$^2$, identical to that used in the cell experiments. The disk was then developed, and the transferred photoresist pattern was examined to determine the UV exposure profile produced by the setup, as shown in **Figure S42**. The measured pattern widths closely matched the designed photomask width, with deviations of no more than 2%, confirming effective UV blocking within the defined region.

### Supporting Figures

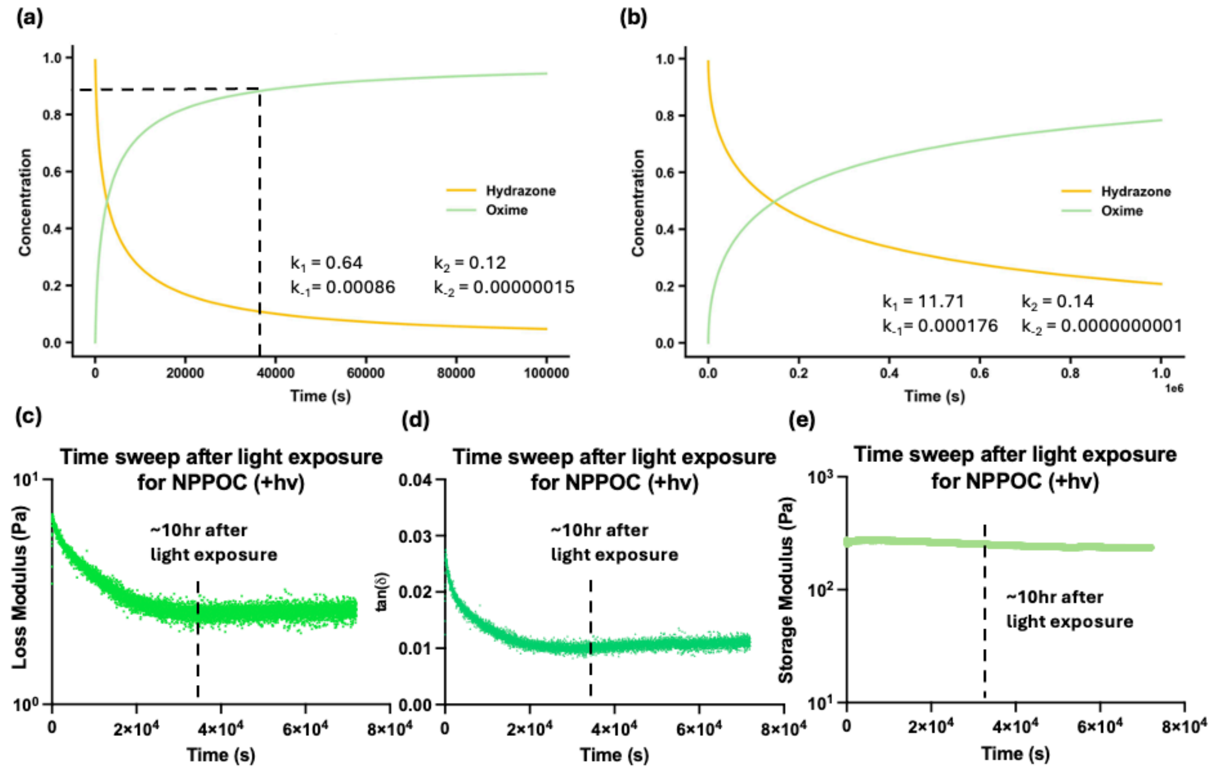

**Figure S1.** ODE computation results with (a) the rate constants retrieved from [8] and (b) other rate constants within published the range for each. Time sweeps of ELP-PEG hydrogels containing NPPOC-protected PEGs after light exposure showing (c) loss modulus, (d) tan(δ), and (e) storage modulus.

**(a)**

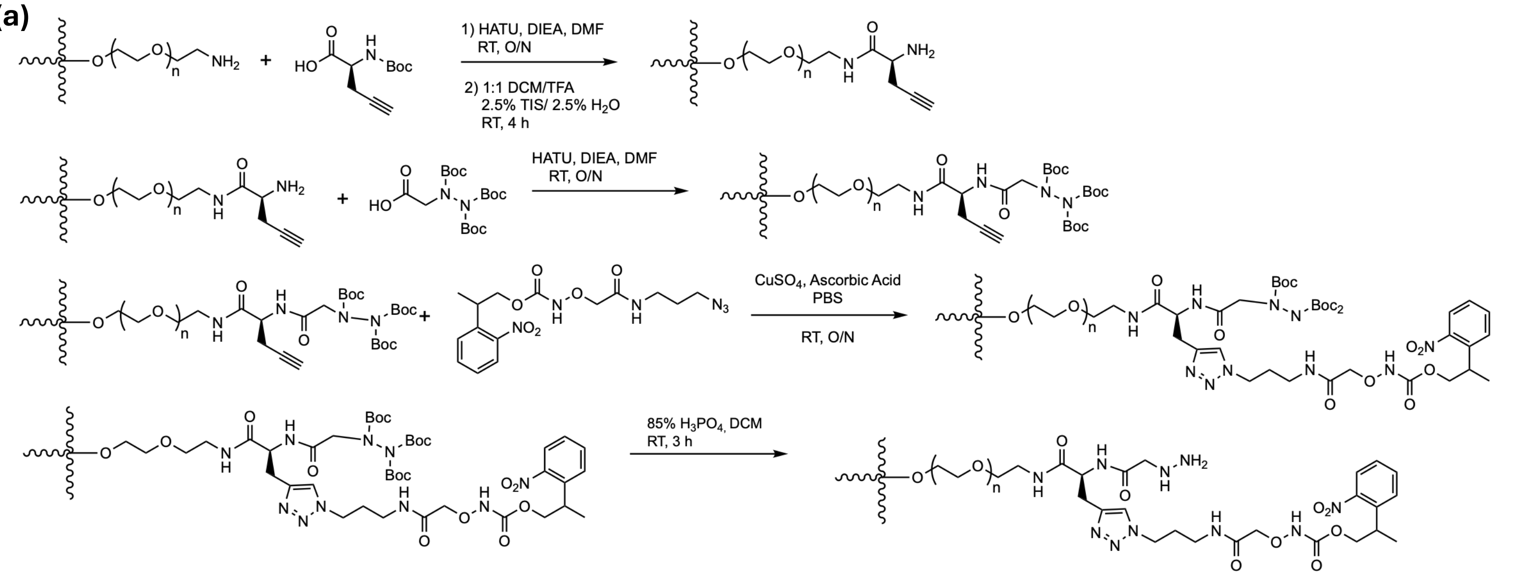

**(b)**

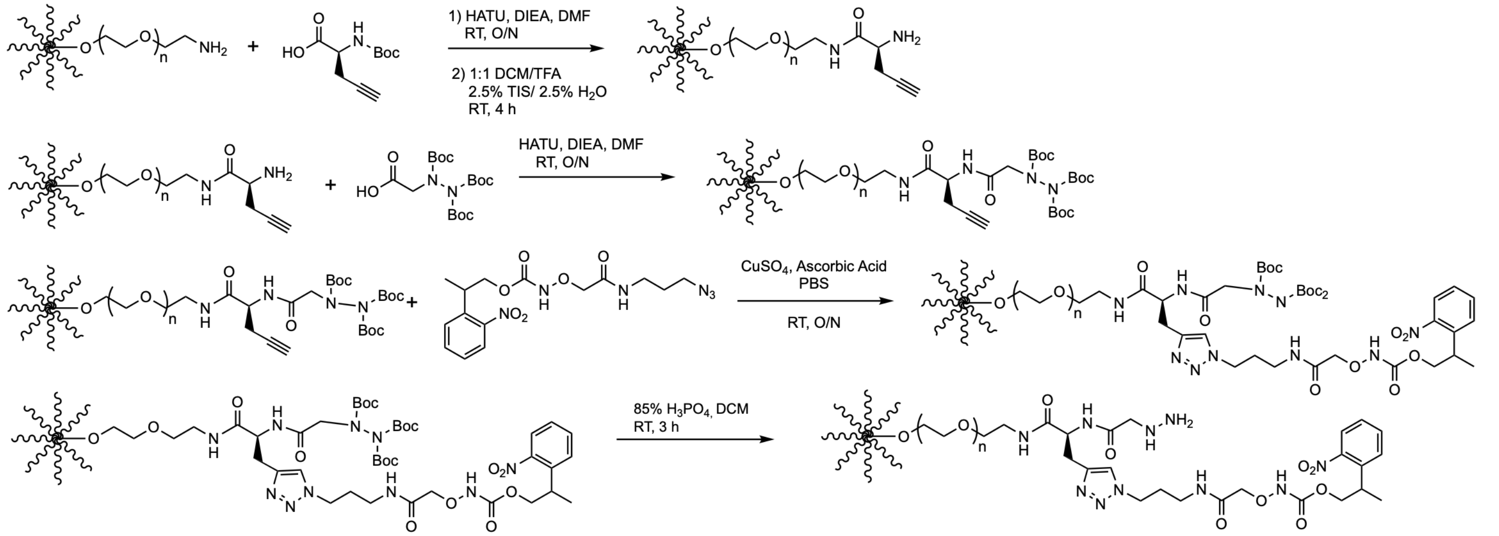

**Figure S2.** Schematics of the synthetic routes for **(a)** 4-arm and **(b)** 8-arm 20 kDa PEG-[8Hz/8HNO-NPPOC]

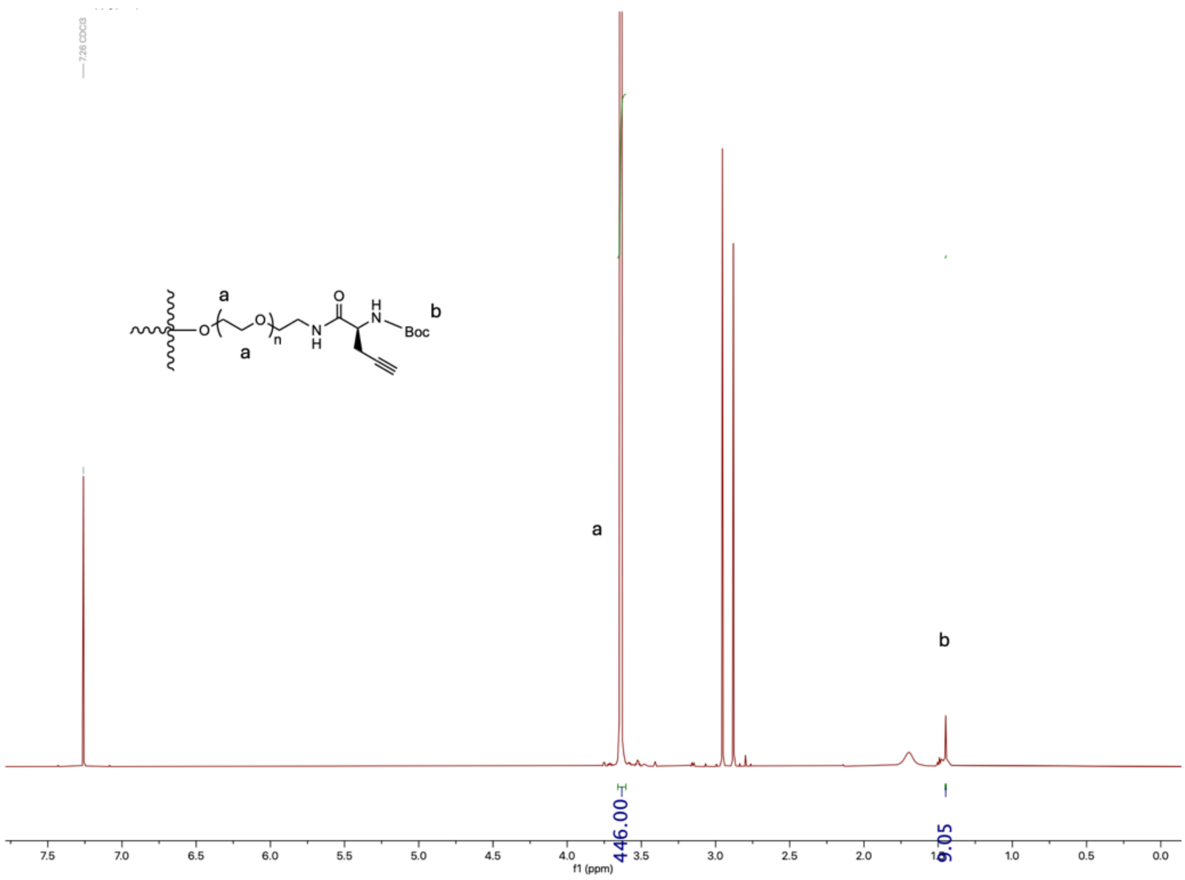

**Figure S3.** $^{1}H$ NMR of PEG-[4NH-Boc/4alkyne] in $CDCl_3$.

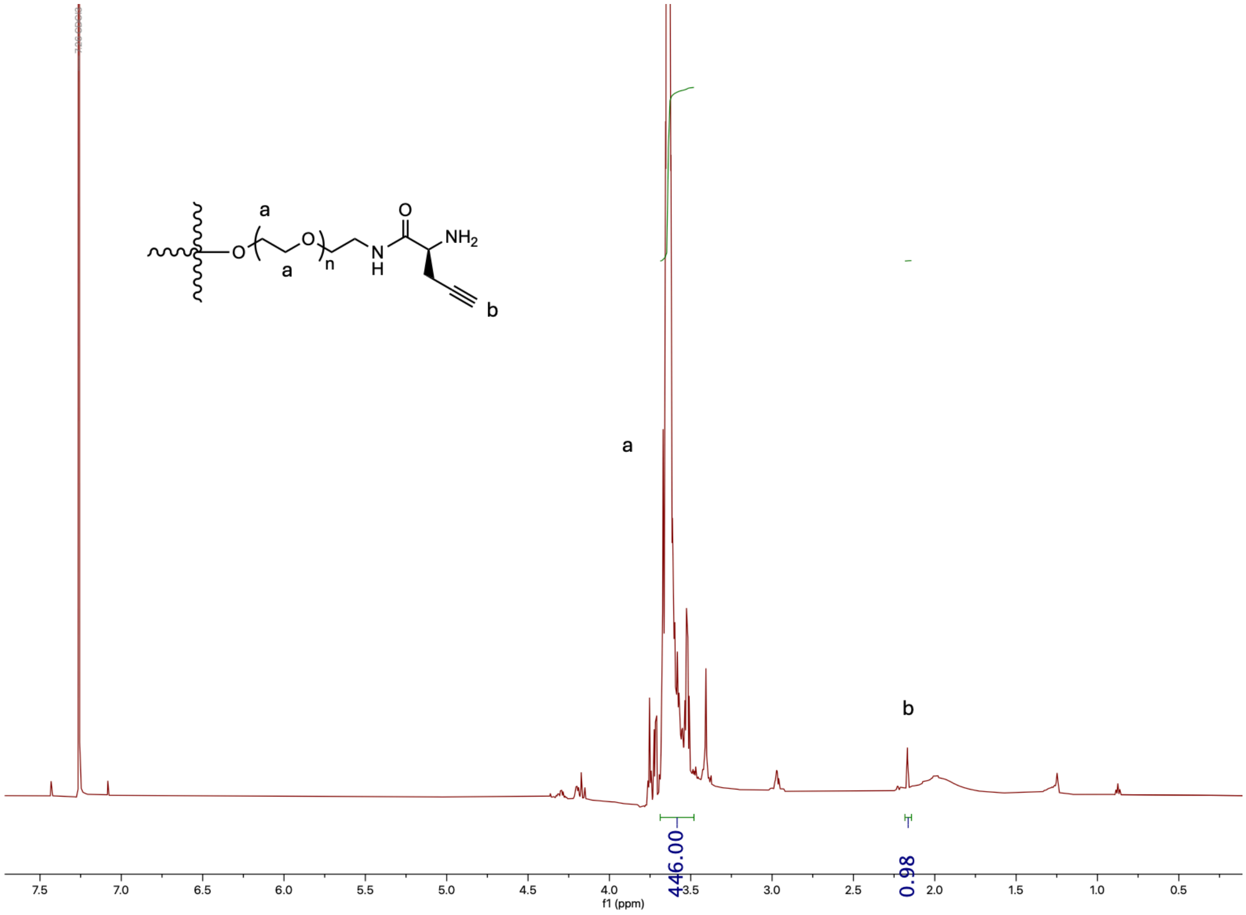

**Figure S4.** $^1$H NMR of PEG-[4NH$_2$/4alkyne] in CDCl$_3$.

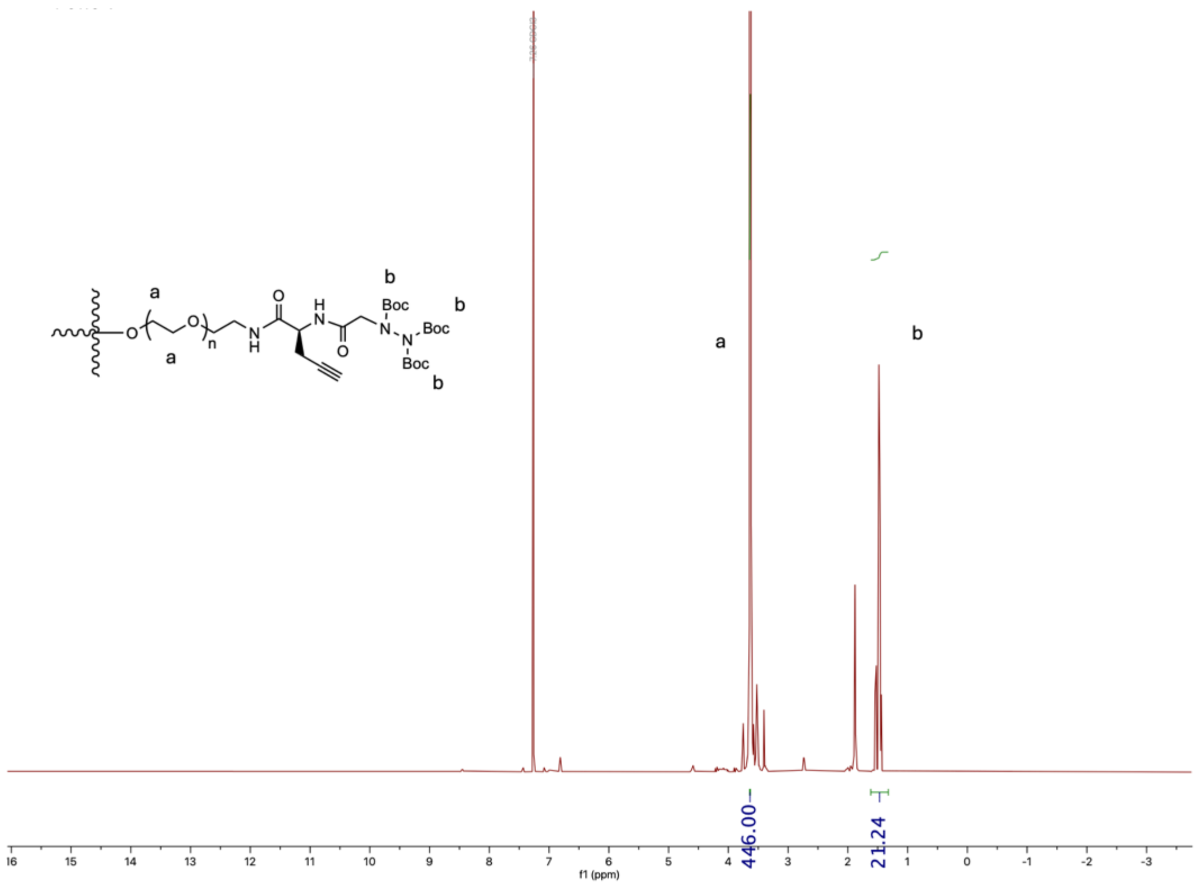

**Figure S5.** $^{1}H$ NMR of PEG-[4Hz-Boc$_3$/4alkyne] in $CDCl_3$.

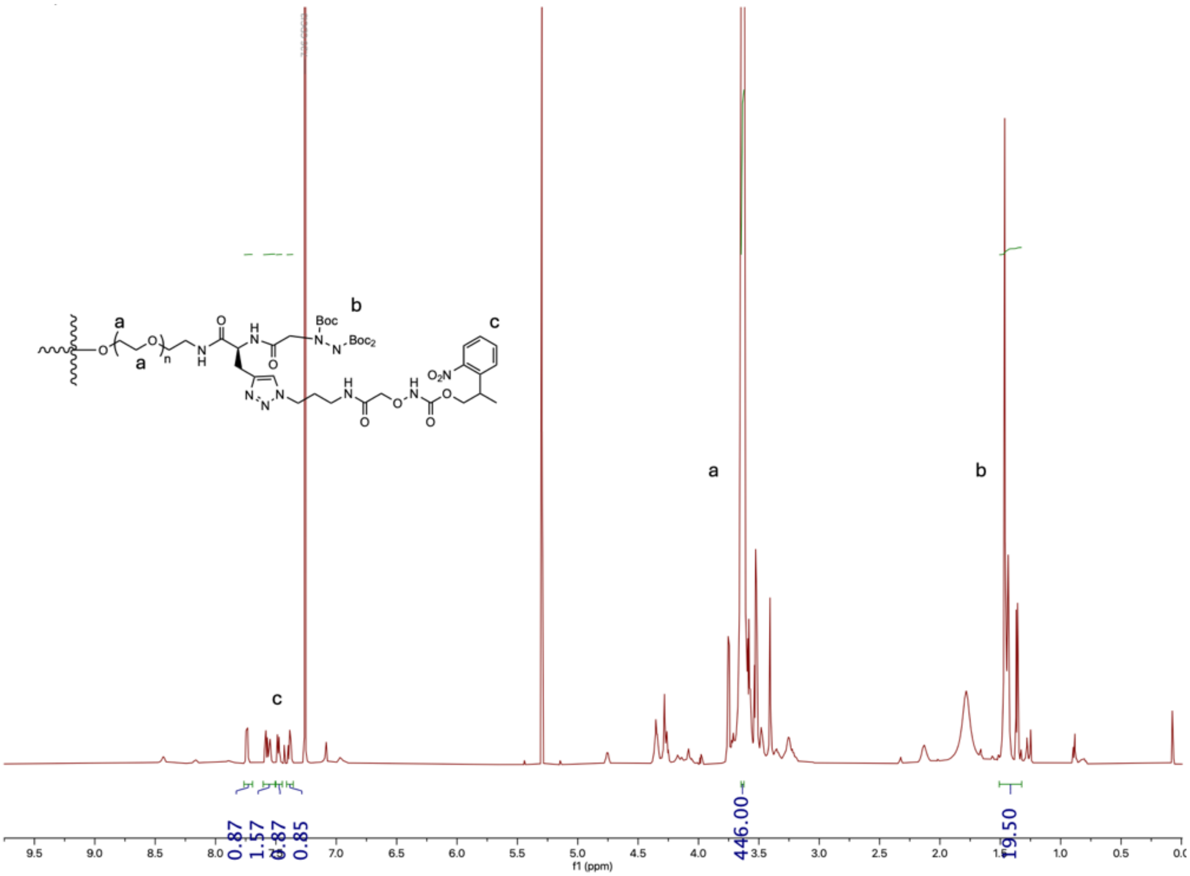

**Figure S6.** [1]H NMR of PEG-[4Hz-Boc$_3$/4HNO-NPPOC] in $CDCl_3$.

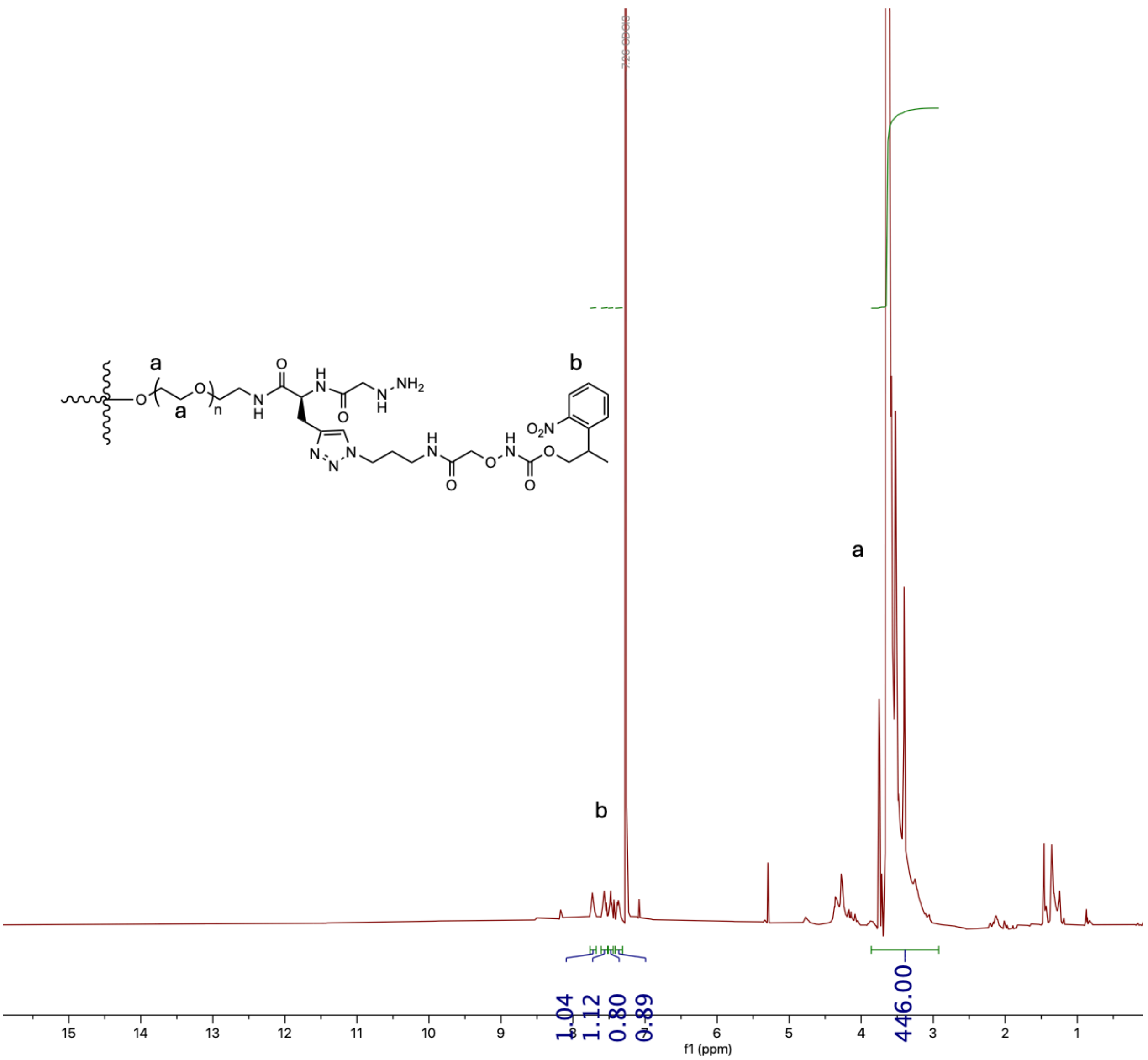

**Figure S7.** [1]H NMR of PEG-[4Hz/4HNO-NPPOC] in DMSO-$d_6$.

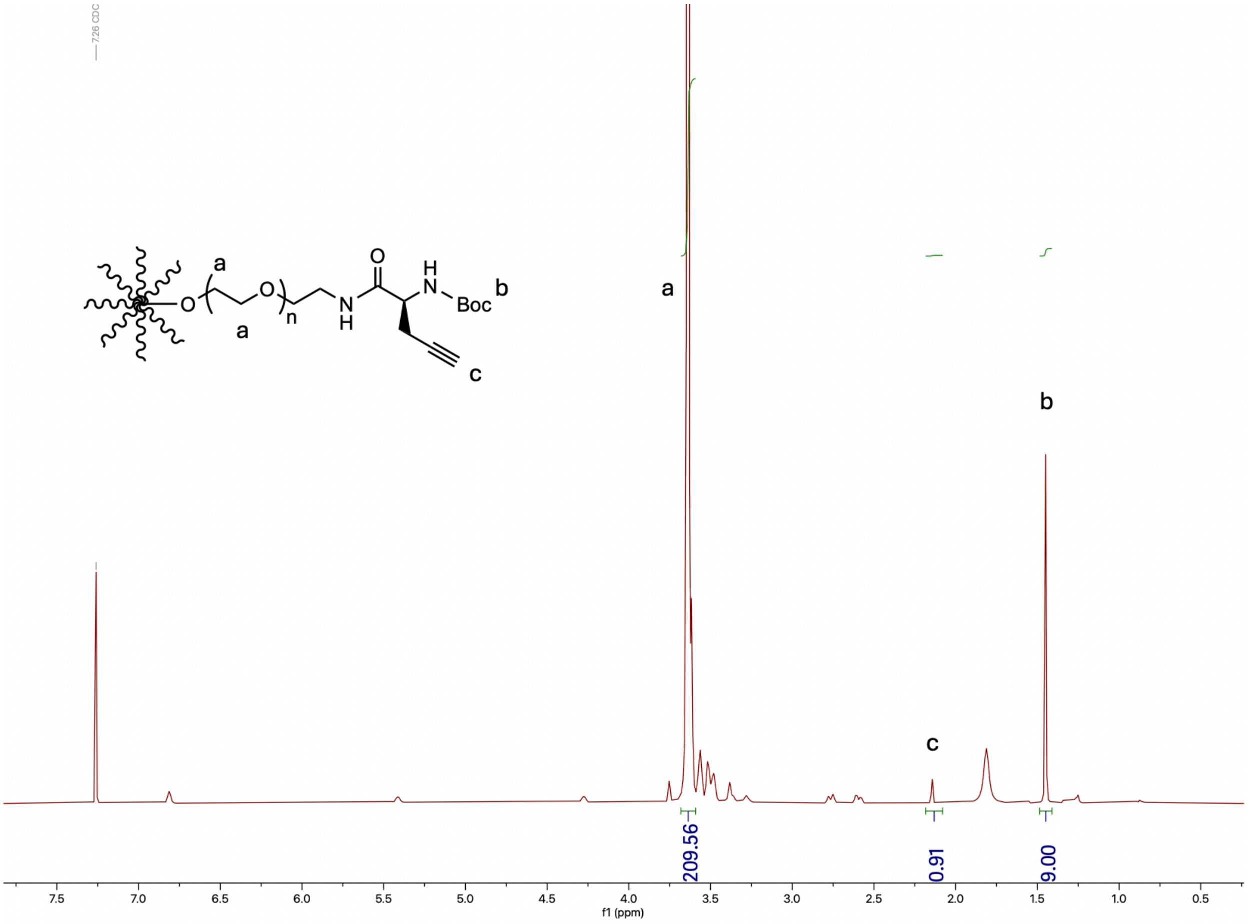

**Figure S8.** $^1$H NMR of PEG-[8NH-Boc/8alkyne] in $CDCl_3$.

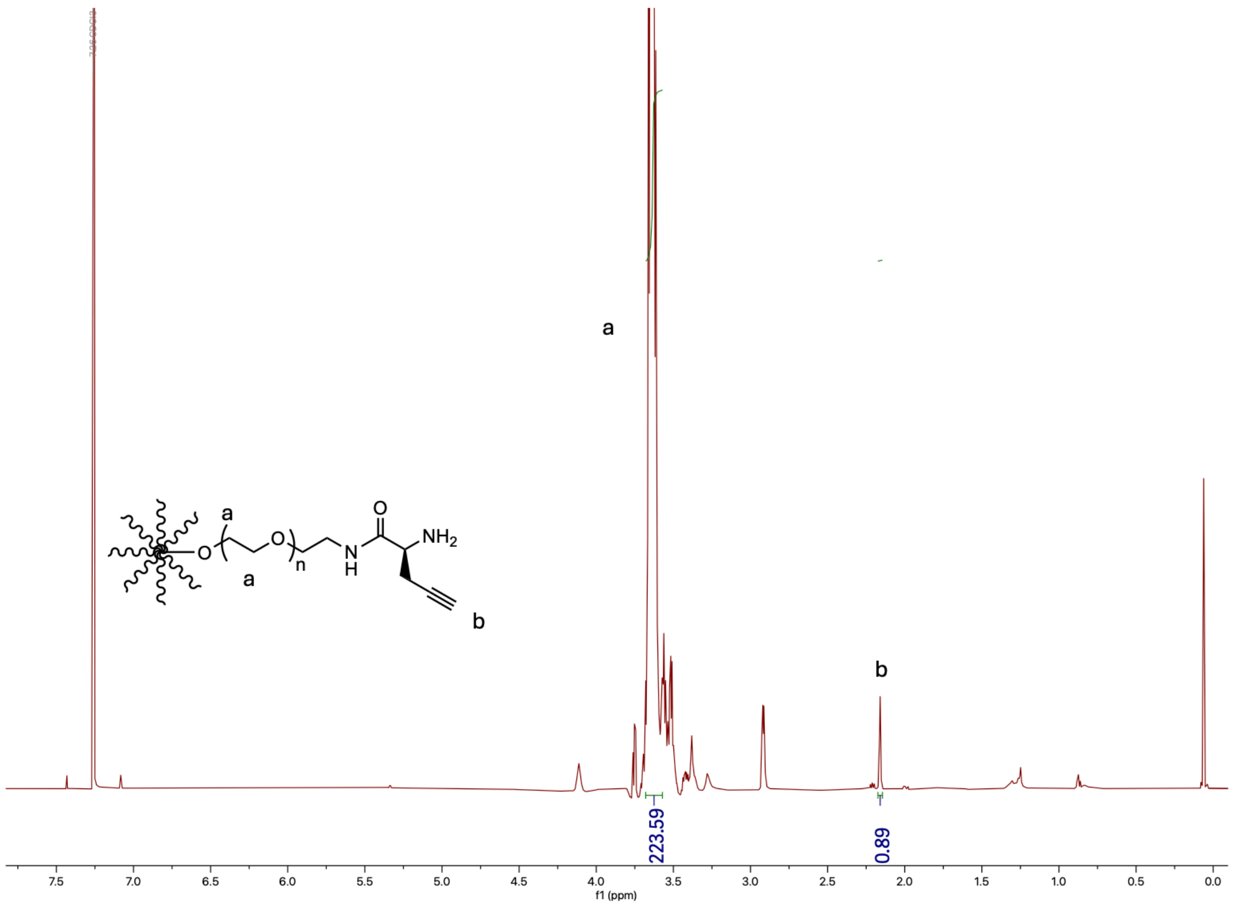

**Figure S9.** $^1$H NMR of PEG-[8NH$_2$/8alkyne] in CDCl$_3$.

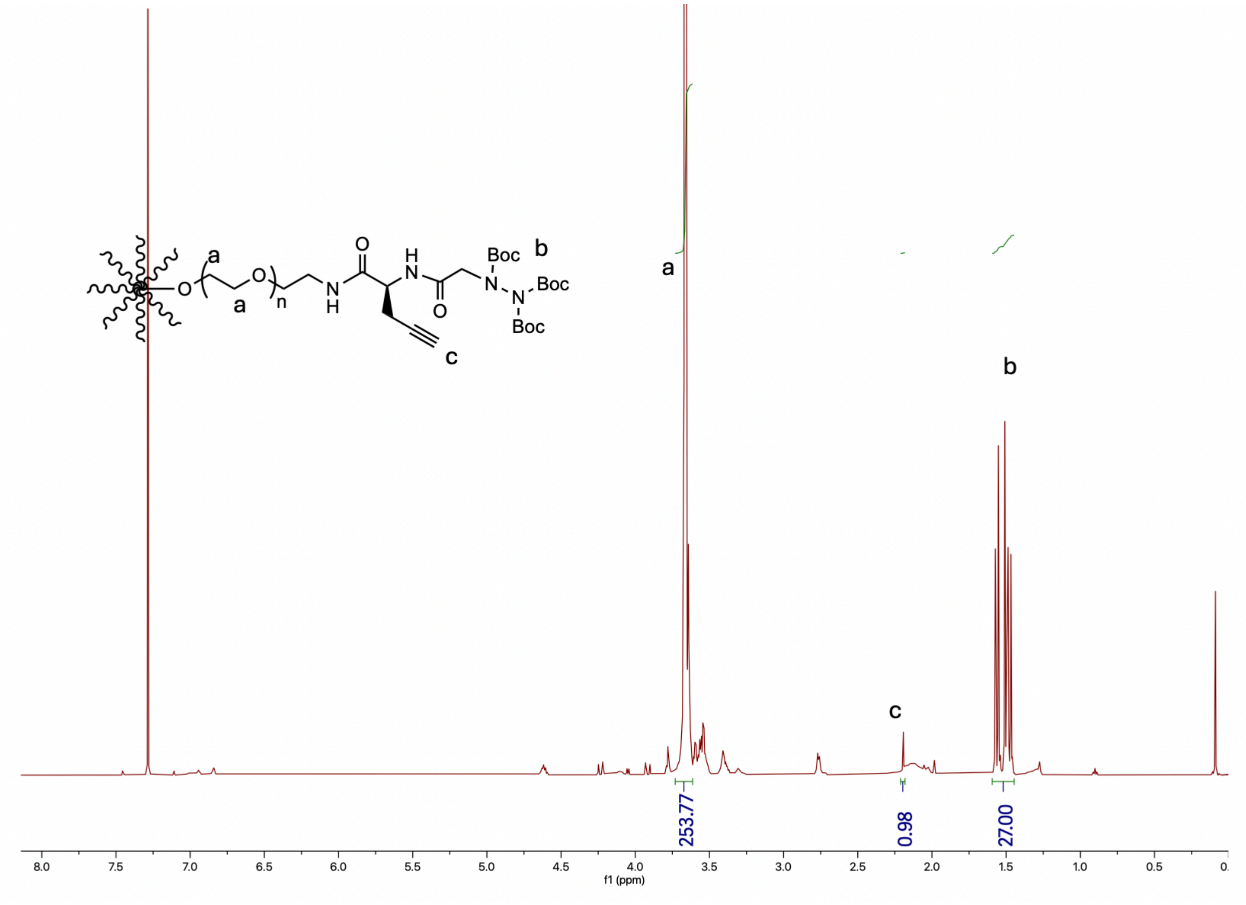

**Figure S10.** $^{1}$H NMR of PEG-[8Hz-Boc$_3$/8alkyne] in $CDCl_3$.

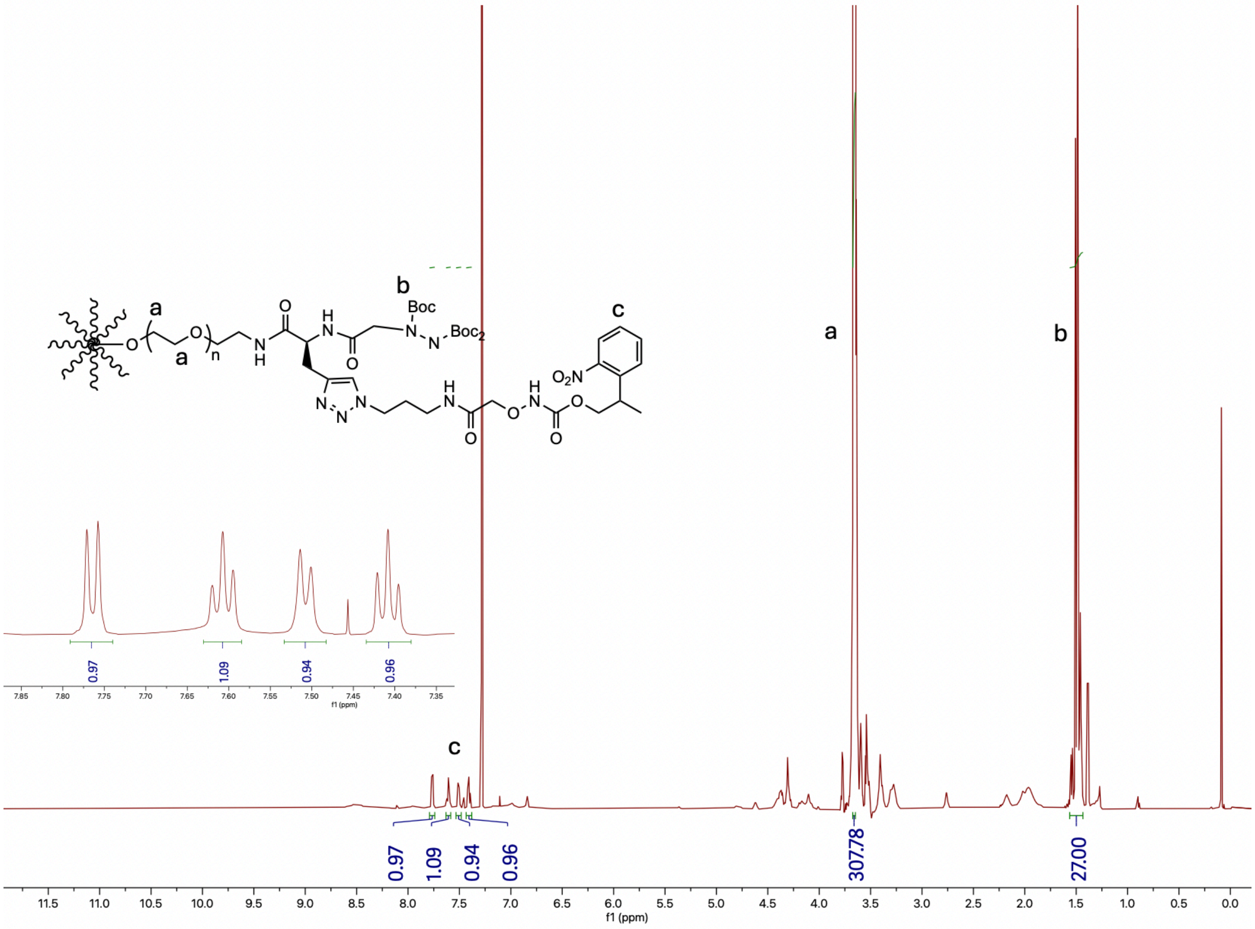

**Figure S11.** $^1$H NMR of PEG-[8Hz-Boc$_3$/8HNO-NPPOC] in $CDCl_3$.

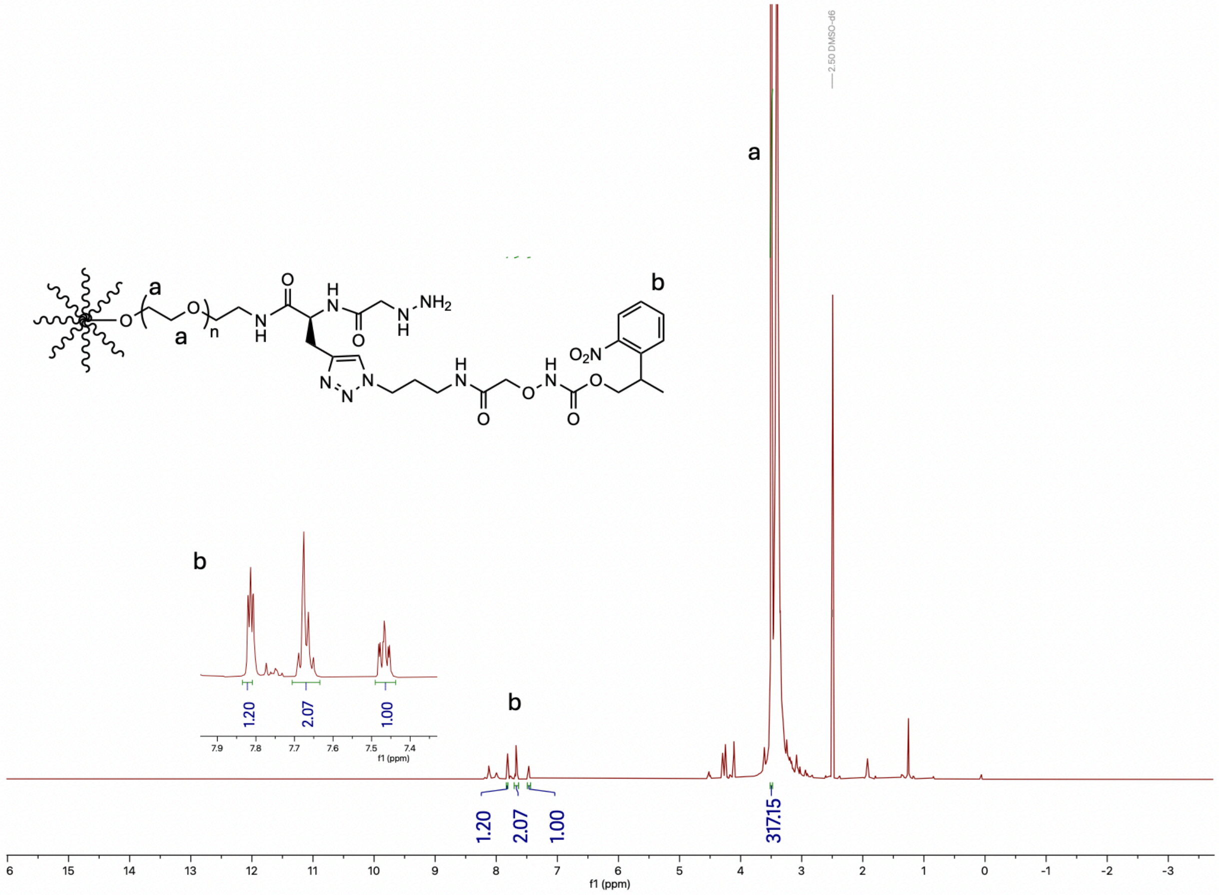

**Figure S12.** [1]H NMR of PEG-[8Hz/8HNO-NPPOC] in DMSO-$d_6$.

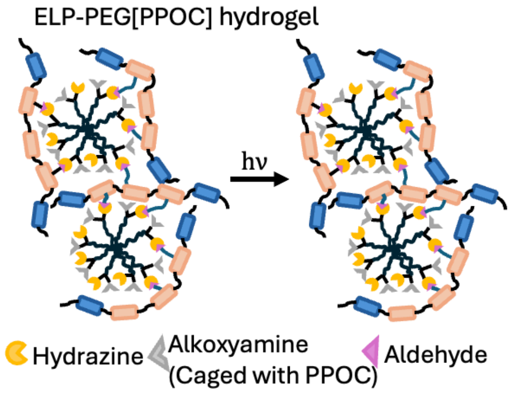

**Figure S13**. Schematic of ELP-PEG[PPOC] hydrogel.

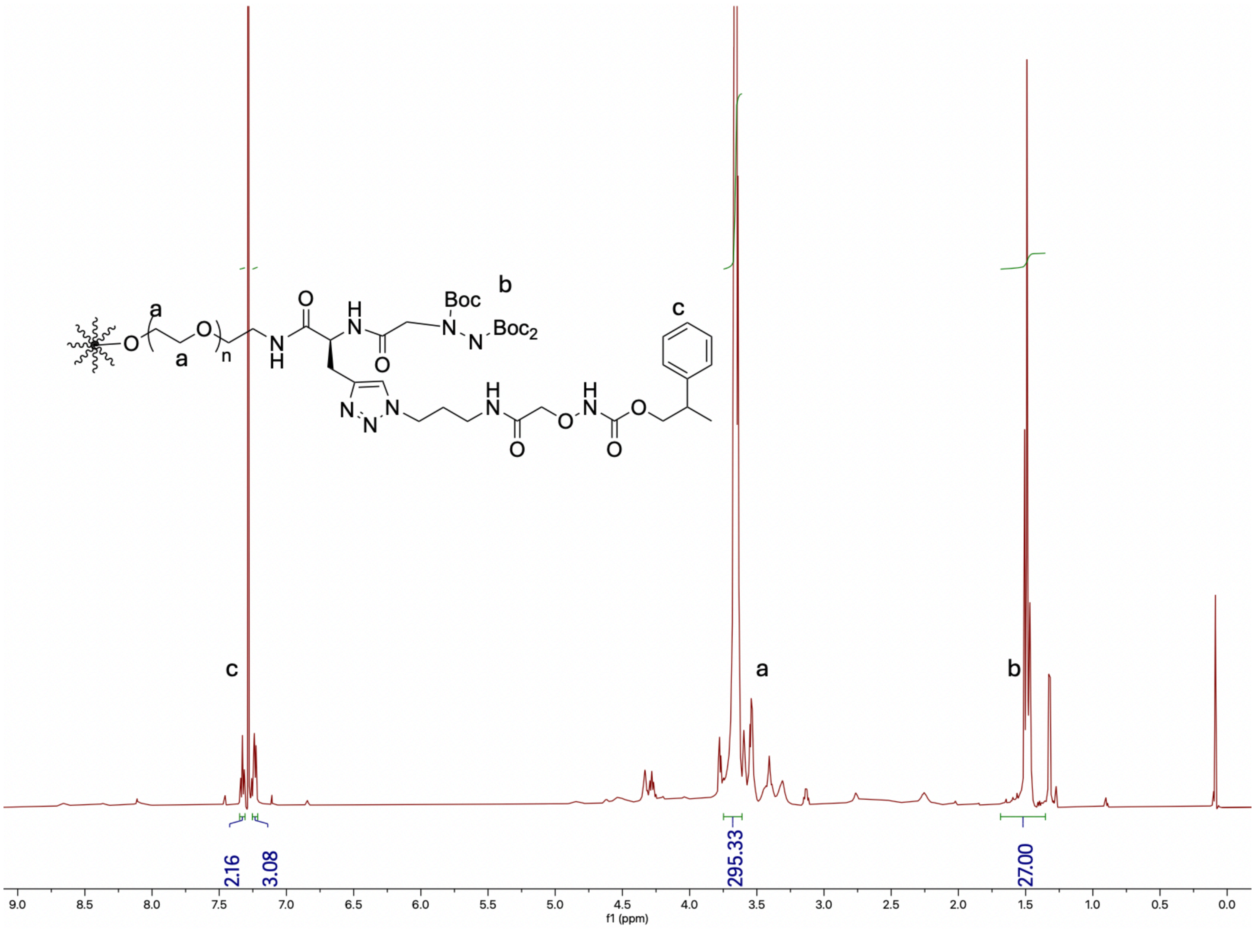

**Figure S14.** $^{1}$H NMR of PEG-[8Hz-Boc$_3$/8HNO-PPOC] in $CDCl_3$.

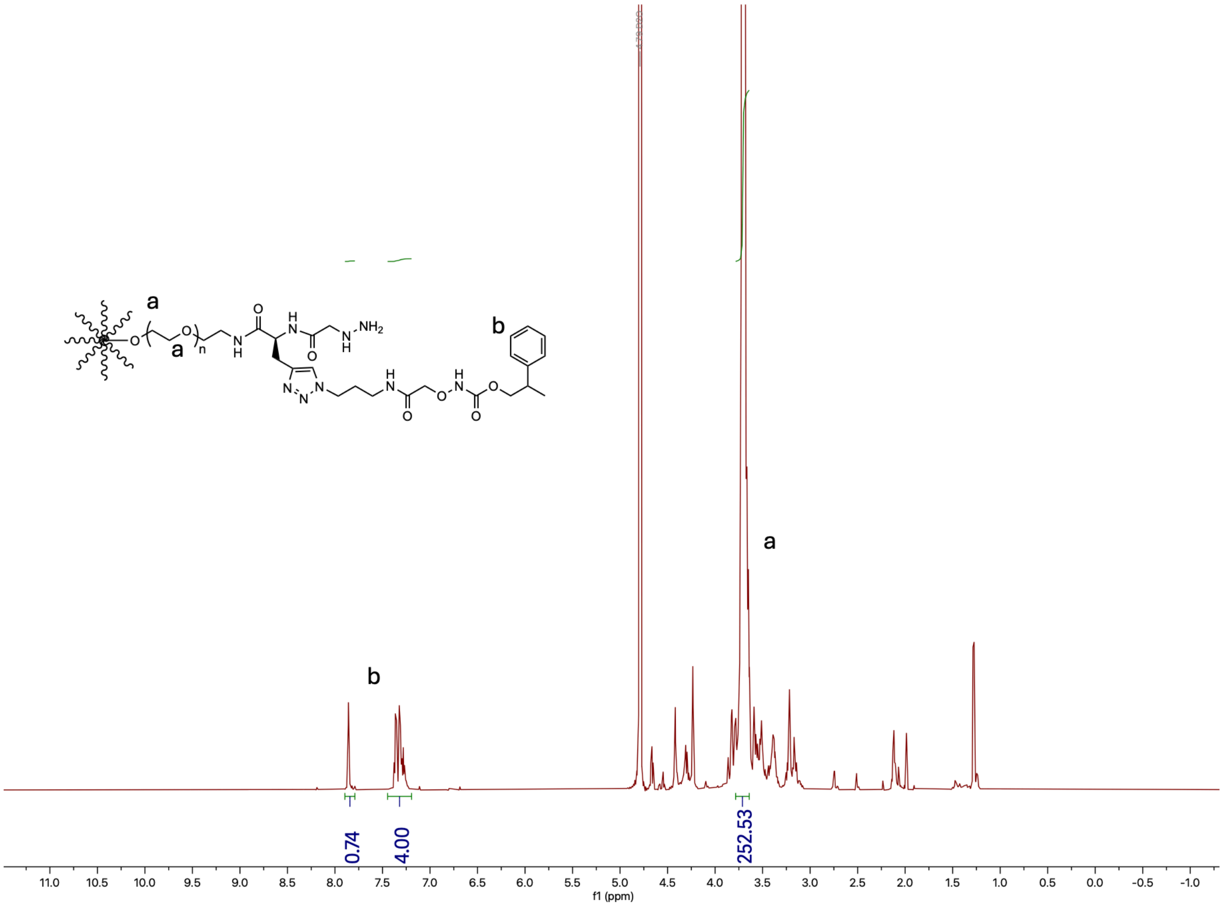

**Figure S15.** $^{1}$H NMR of PEG-[8Hz /8HNO-PPOC] in $CDCl_3$.

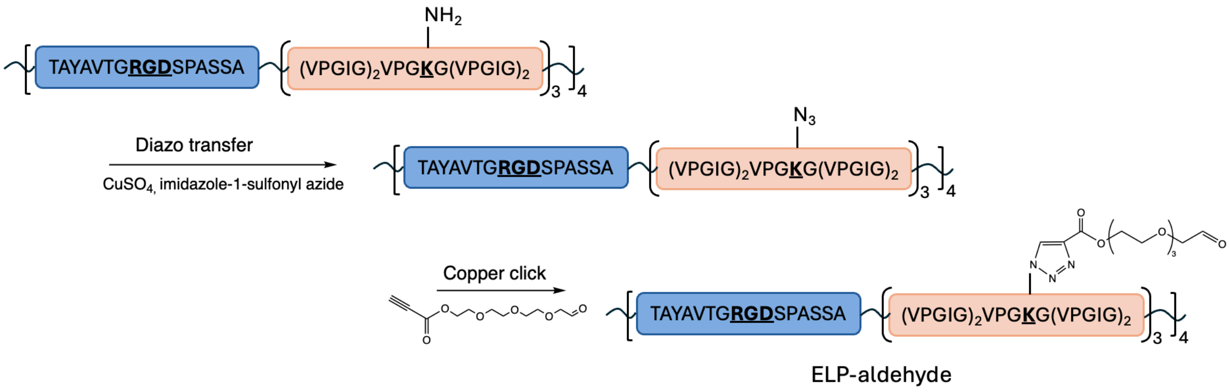

**Figure S16.** Schematic of the synthetic route of ELP-ALD.

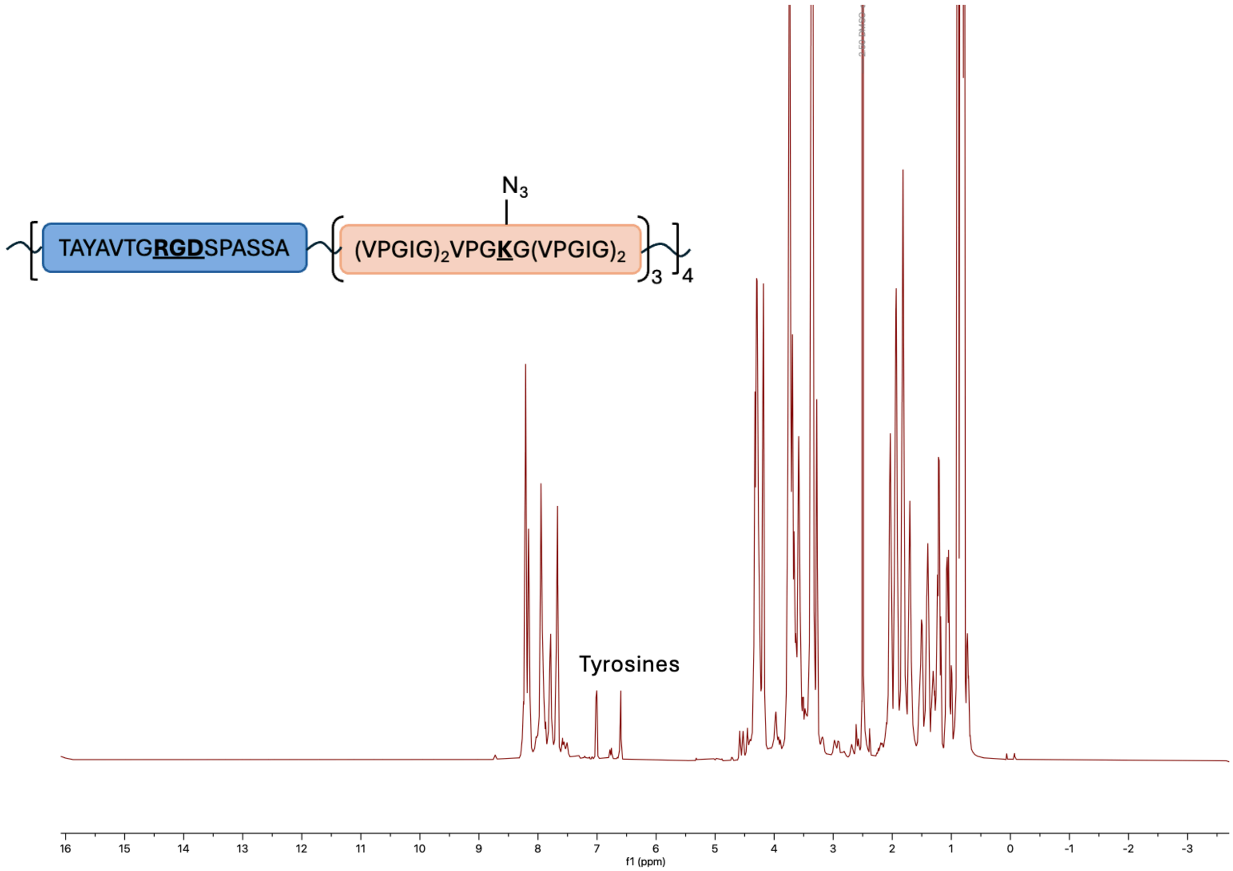

**Figure S17.** $^{1}$H NMR of ELP-azide in DMSO-$d_6$.

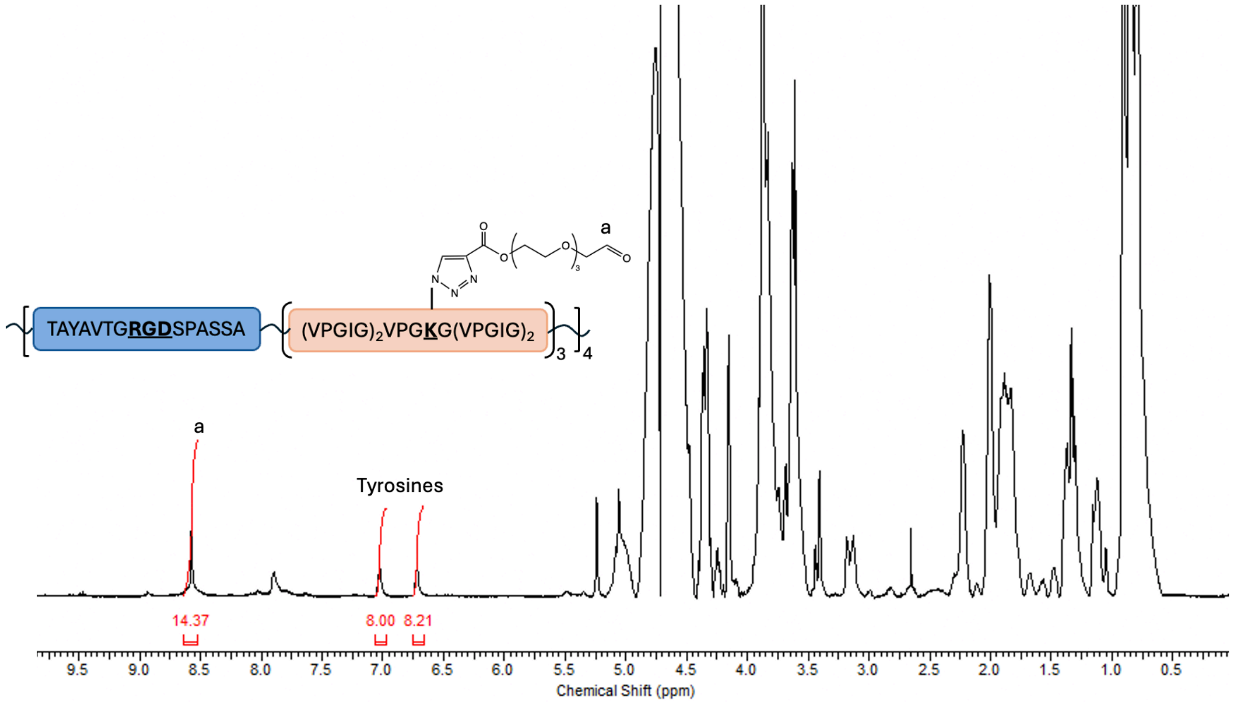

**Figure S18.** $^1$H NMR of ELP-aldehyde (ELP-ALD) in $D_2O$.

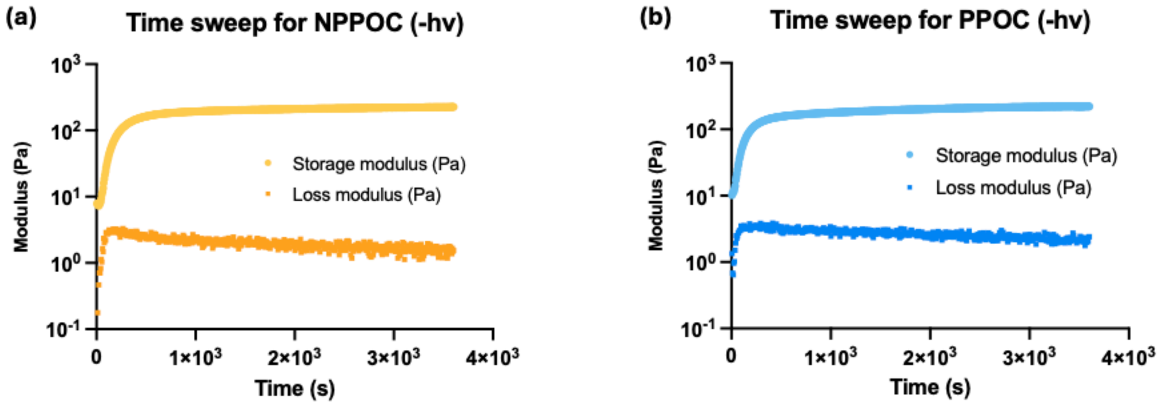

**Figure S19.** Representative time sweeps of ELP-PEG hydrogels gelation for (a) NPPOC- and (b) PPOC-protected PEGs before light treatment

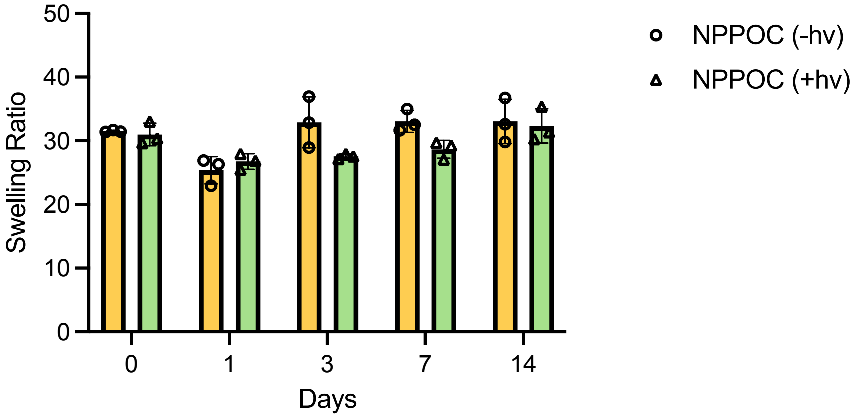

**Figure S20.** Mass swelling ratio as a function of time for hydrogels with final concentrations of 1.5% (w/v) ELP-ALD and 1.95 %(w/v) PEG-[8Hz/8HNO-NPPOC] with or without light. Data are presented as mean ± s.d., n = 3.

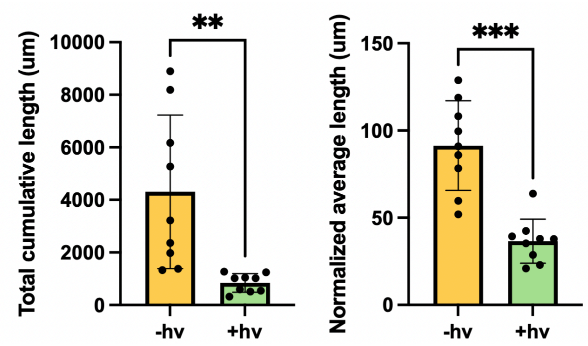

**Figure S21.** Quantified results of total cumulative neurite length and normalized average neurite length untreated and light-treated ELP-PEG hydrogels containing MN spheroids.

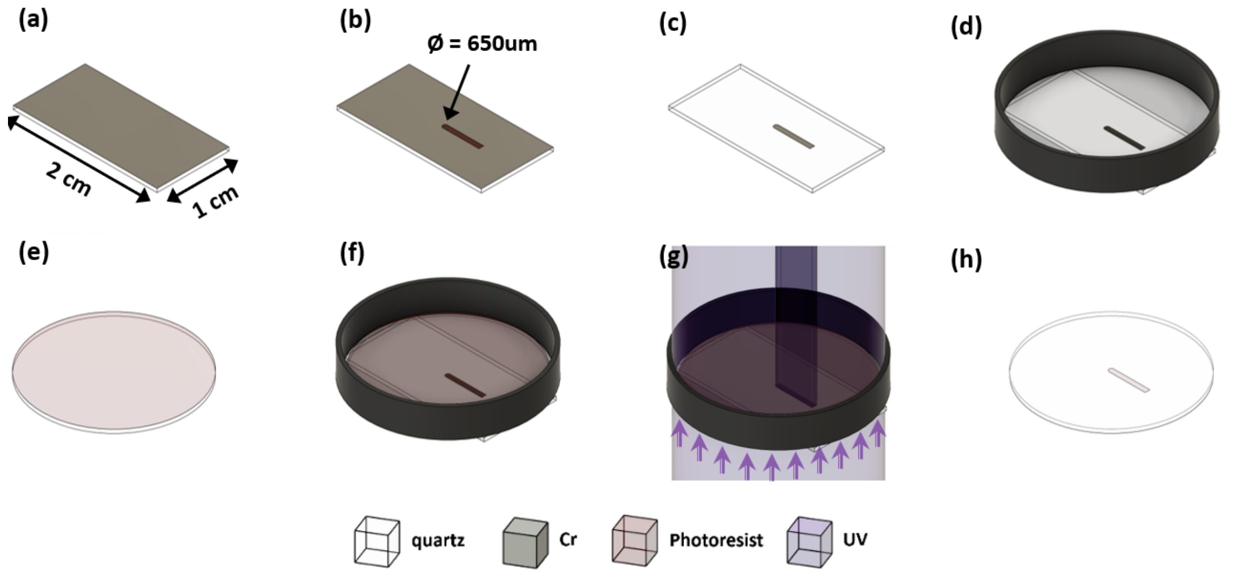

**Figure S22.** Schematics of fabrication and testing of the photomask. (a) A 20 mm × 10 mm Cr-coated quartz substrate. (b) Photoresist patterning of the central masking feature. (c) Cr etching and photoresist removal to form the opaque feature on the transparent quartz substrate. (d) Attachment of the photomask beneath a glass-bottom well plate. (e) A 15 mm-diameter, 250 µm-thick circular quartz disk coated with a 10 µm-thick AZ4620 photoresist layer. (f) Placement of the coated quartz disk inside the well. (g) Bottom-side exposure using 365 nm UV light. (h) Development of the AZ4620 layer.

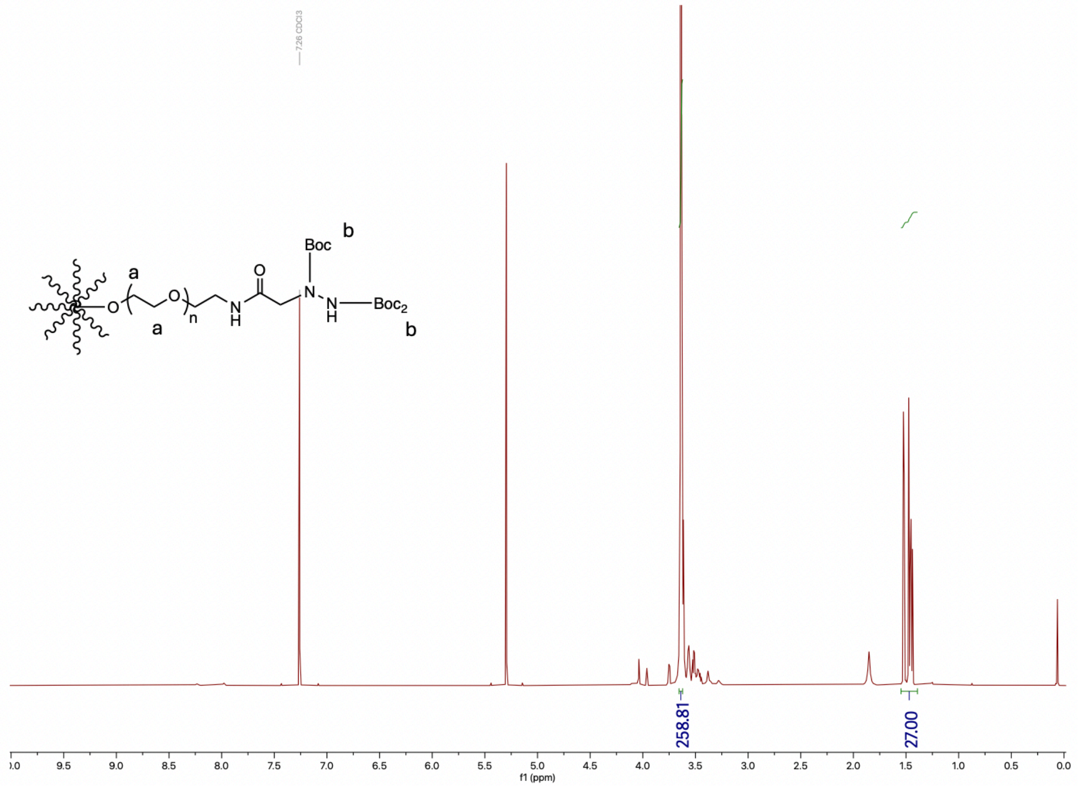

**Figure S23.** [1]H NMR of PEG-hydrazine-$Boc_3$ in $CDCl_3$.

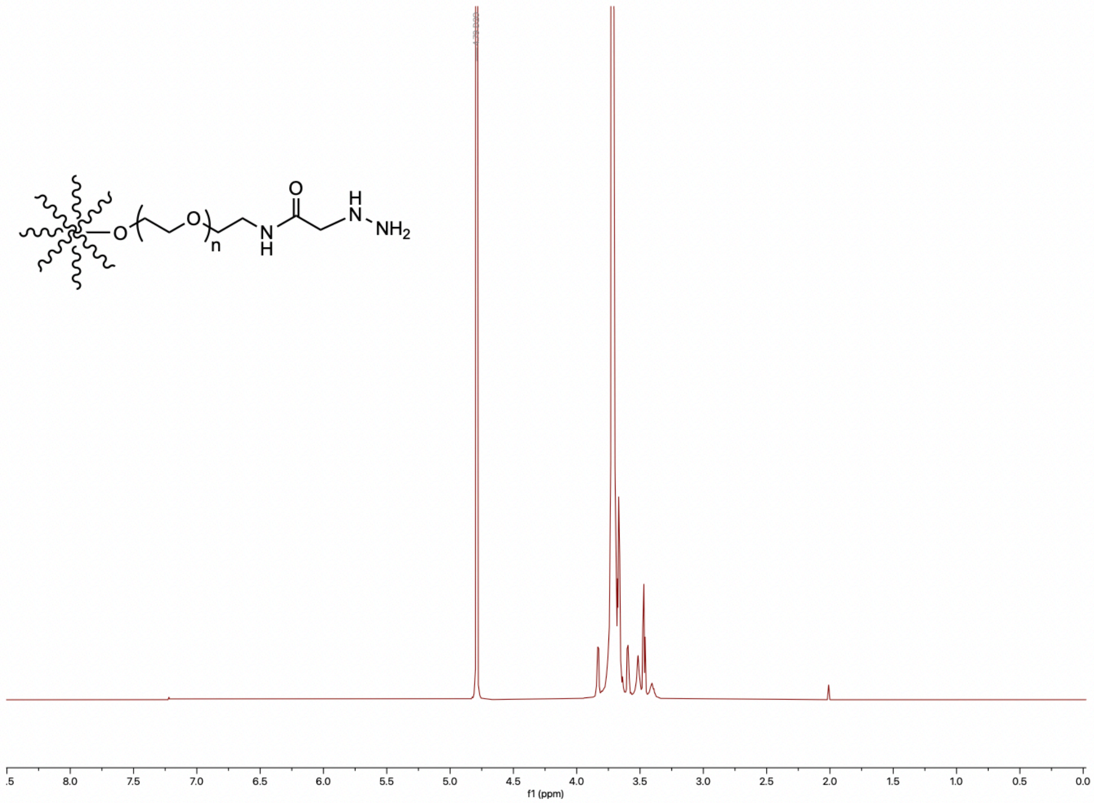

**Figure S24.** [1]H NMR of PEG-hydrazine in $D_2O$.

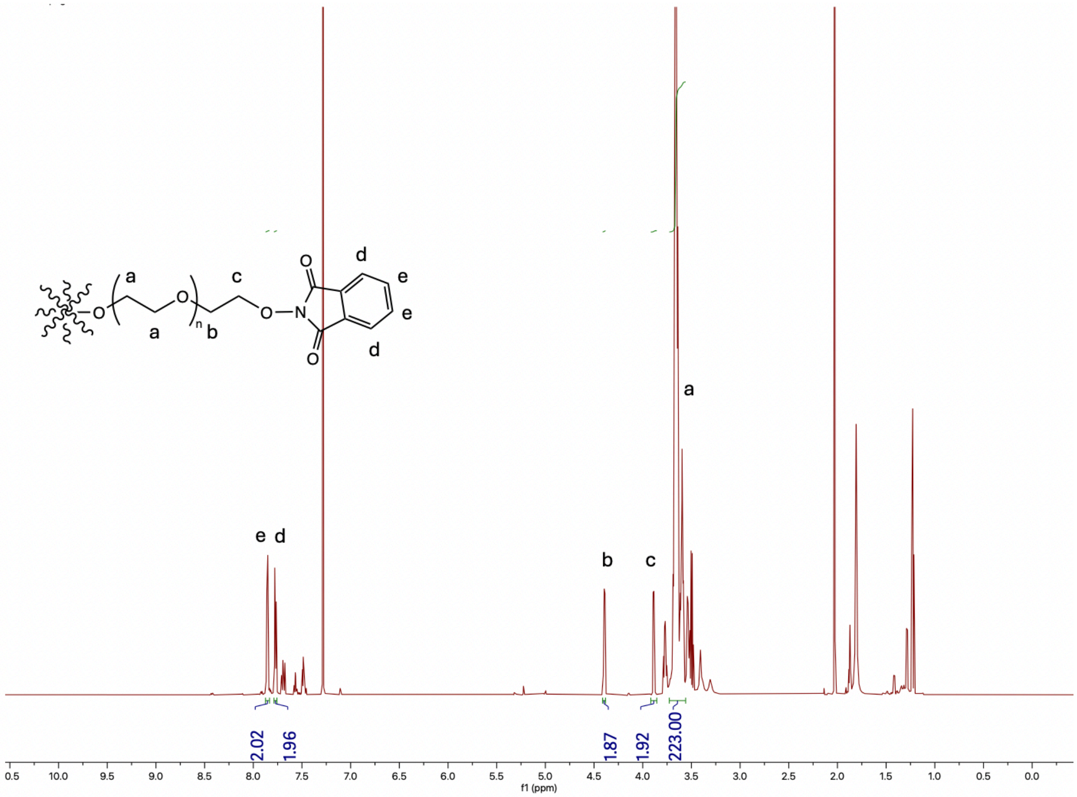

**Figure S25.** [1]H NMR of PEG-[N-hydroxyphthalimide] in $CDCl_3$.

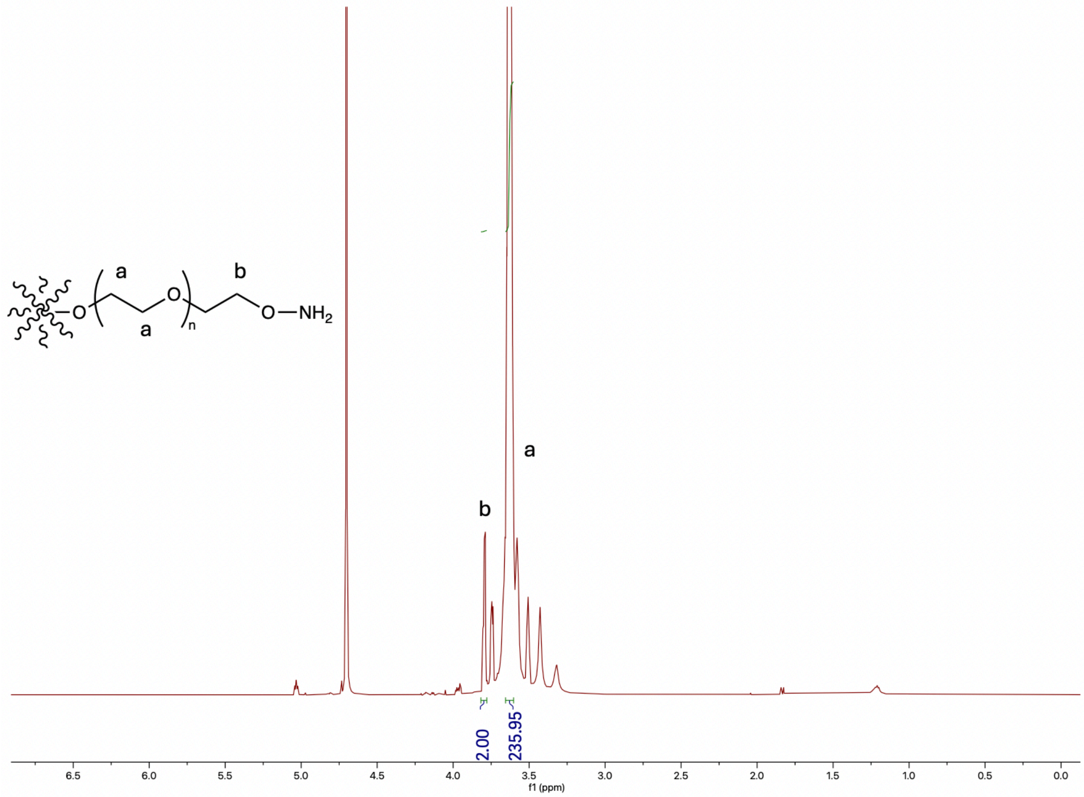

**Figure S26.** $^{1}$H NMR of PEG-alkoxyamine in $D_2O$.

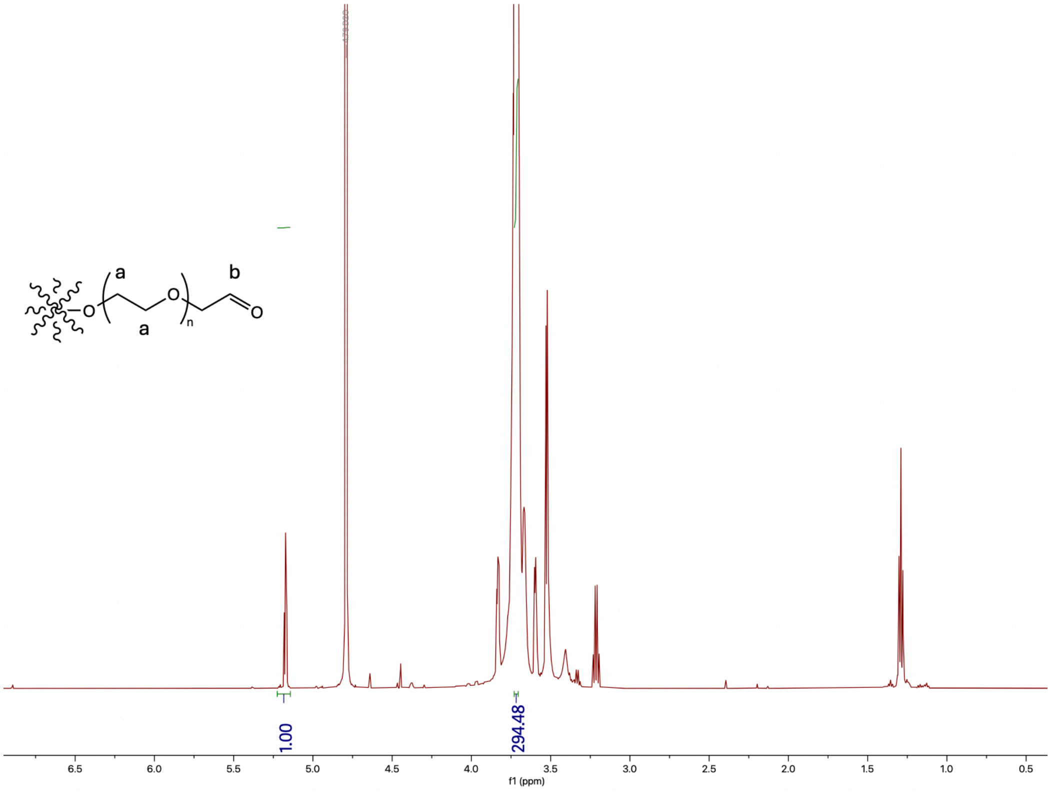

**Figure S27.** $^{1}$H NMR of PEG-aldehyde in $D_2O$.

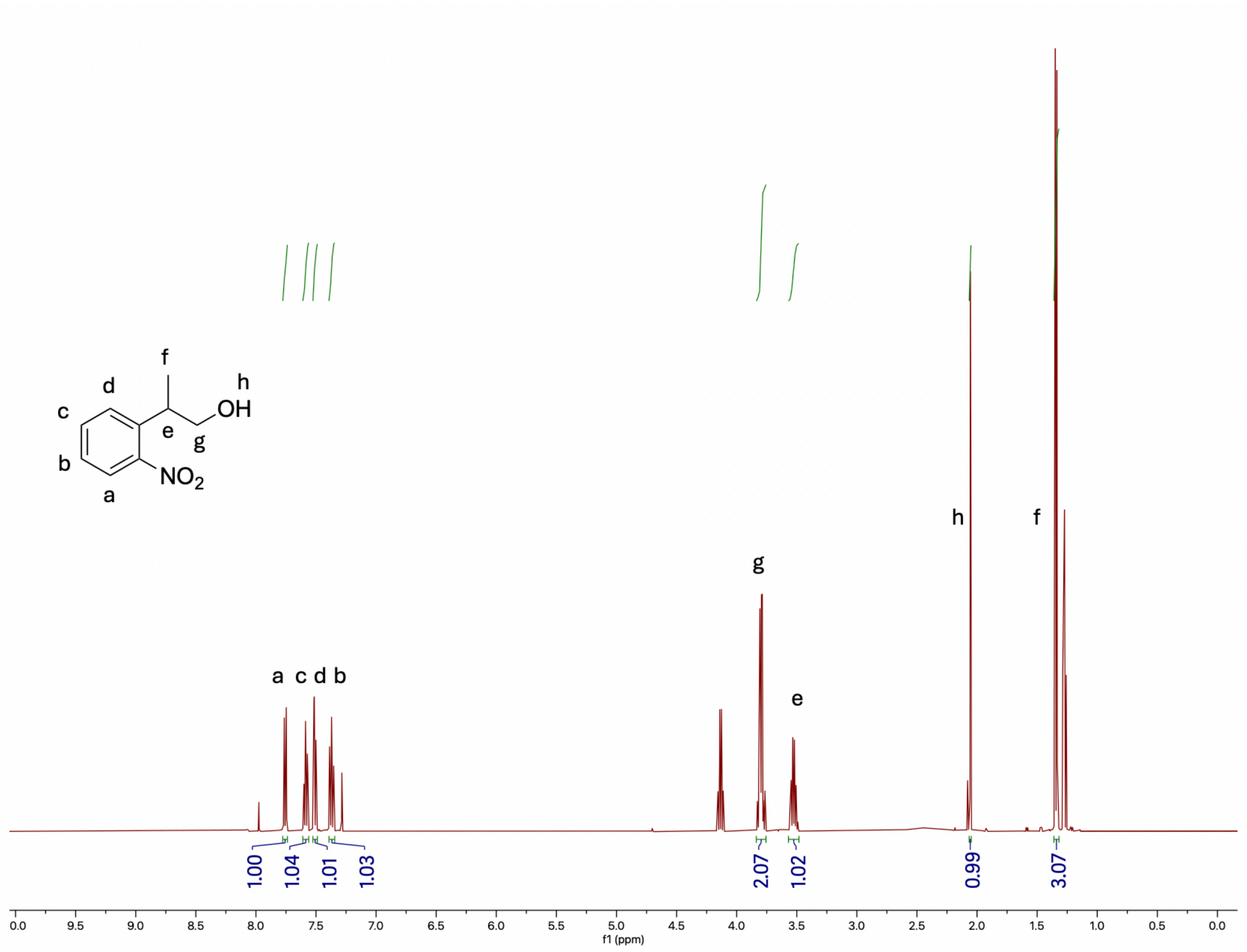

**Figure S28.** [1]H NMR of 2-(2-nitrophenyl)propan-1-ol in $CDCl_3$.

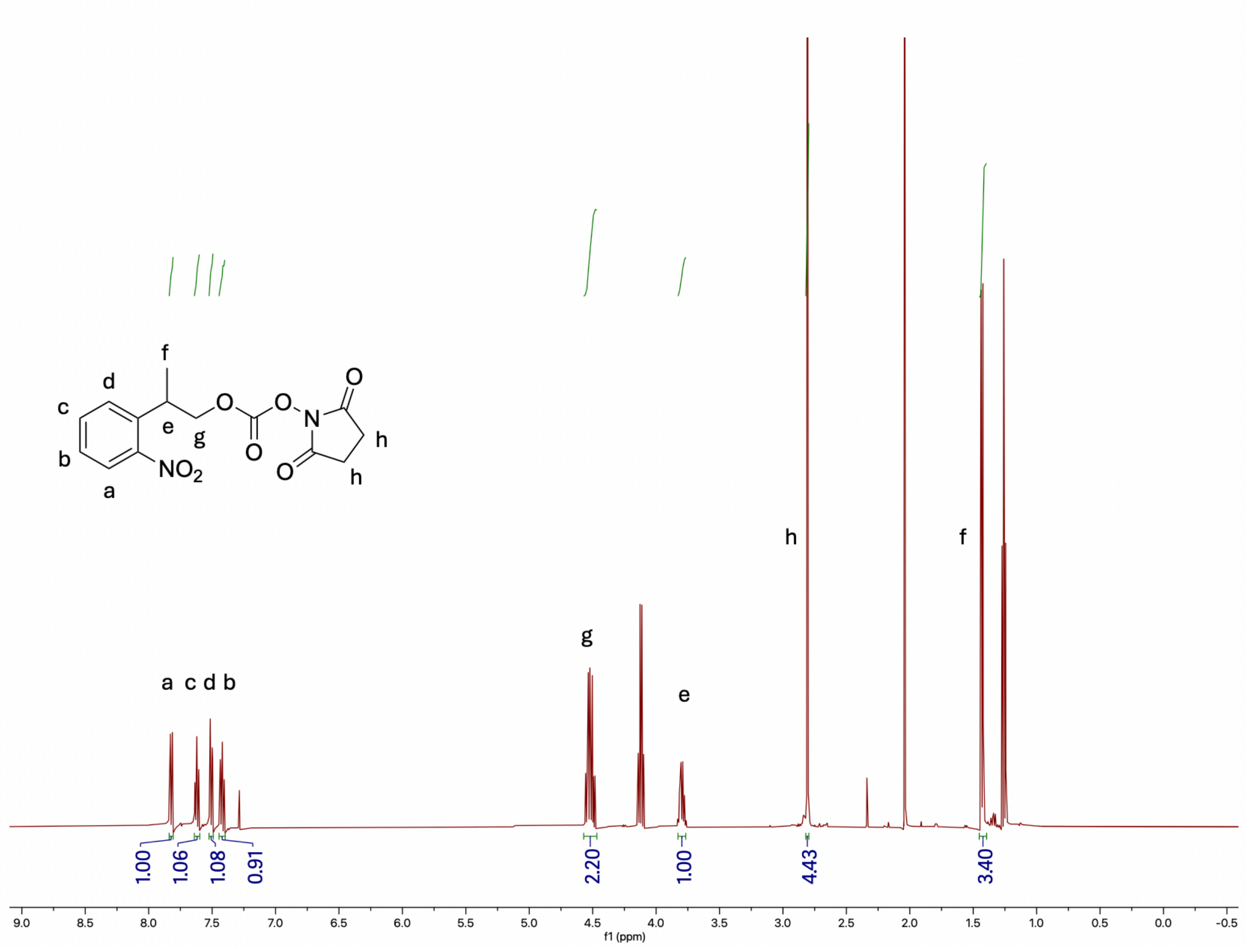

**Figure S29.** $^1$H NMR of 2,5-dioxopyrrolidin-1-yl (2-(2-nitrophenyl)propyl) carbonate (NPPOC-Osu) in $CDCl_3$.

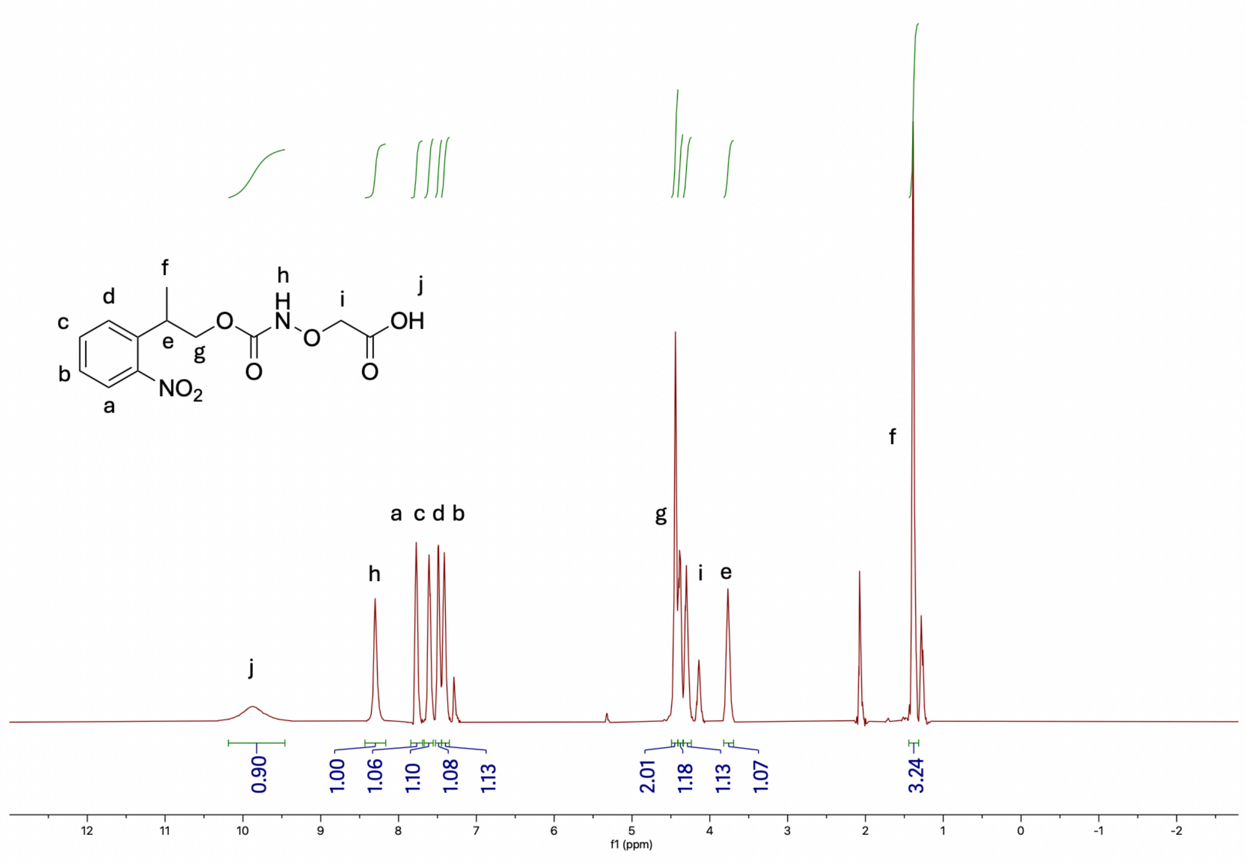

**Figure S30.** $^{1}H$ NMR of 2-((((2-(2-nitrophenyl)propoxy)carbonyl)amino)oxy)acetic acid (NPPOC-HNO-COOH) in $CDCl_3$.

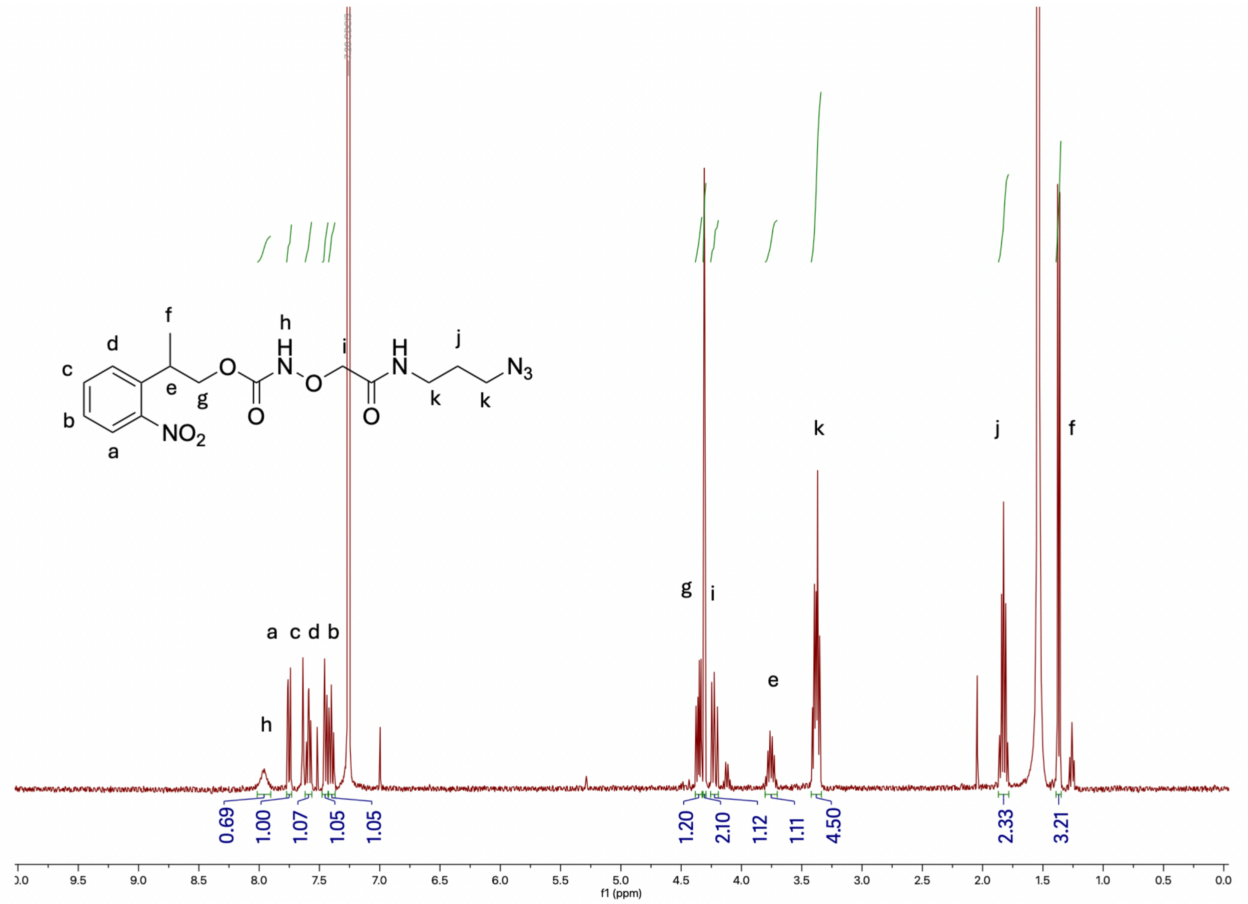

**Figure S31.** $^{1}$H NMR of 2-(2-nitrophenyl)propyl (2-((3-azidopropyl)amino)-2-oxoethoxy)carbamate (NPPOC-HNO-$N_3$) in $CDCl_3$.

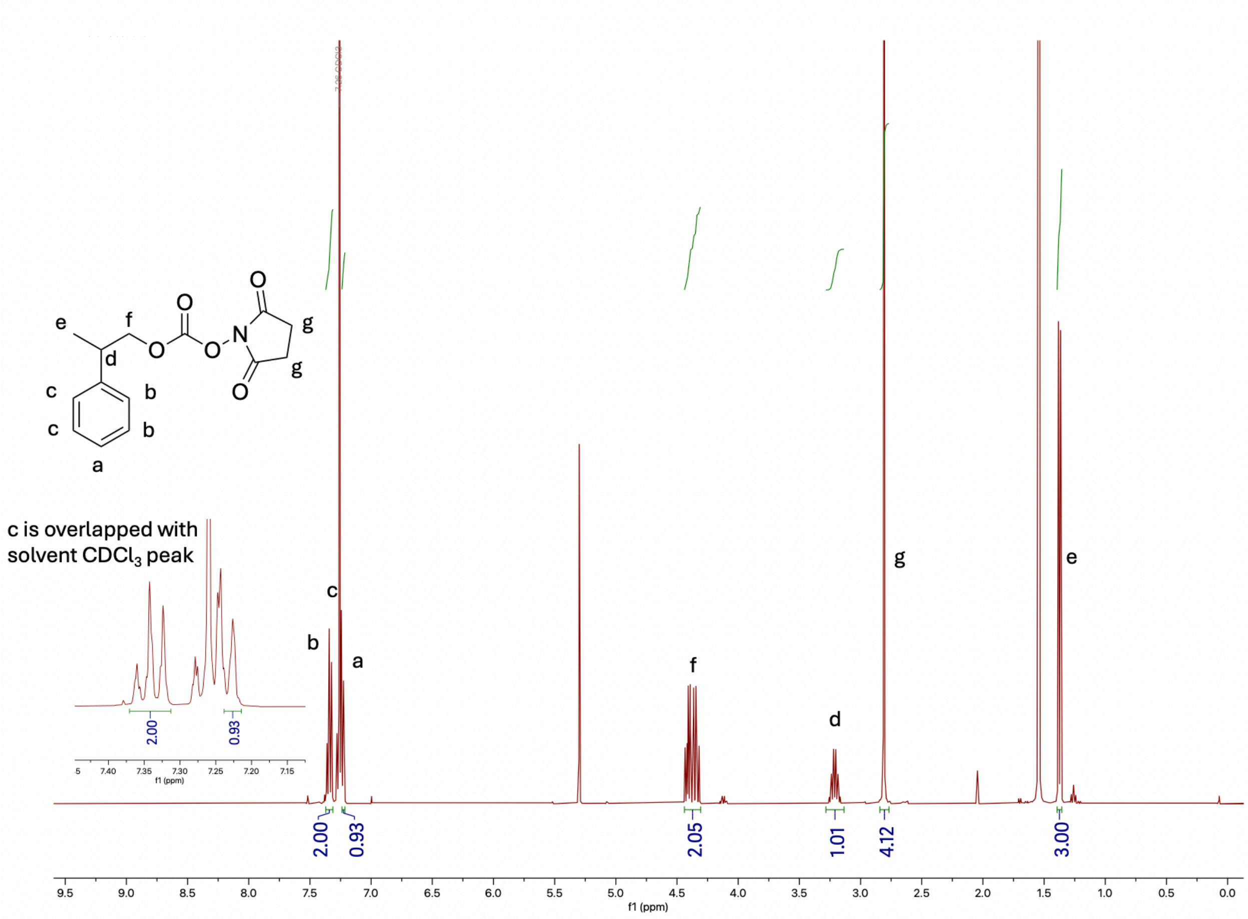

**Figure S32.** $^{1}H$ NMR of 2,5-dioxopyrrolidin-1-yl (2-phenylpropyl) carbonate (PPOC-Osu) in $CDCl_3$.

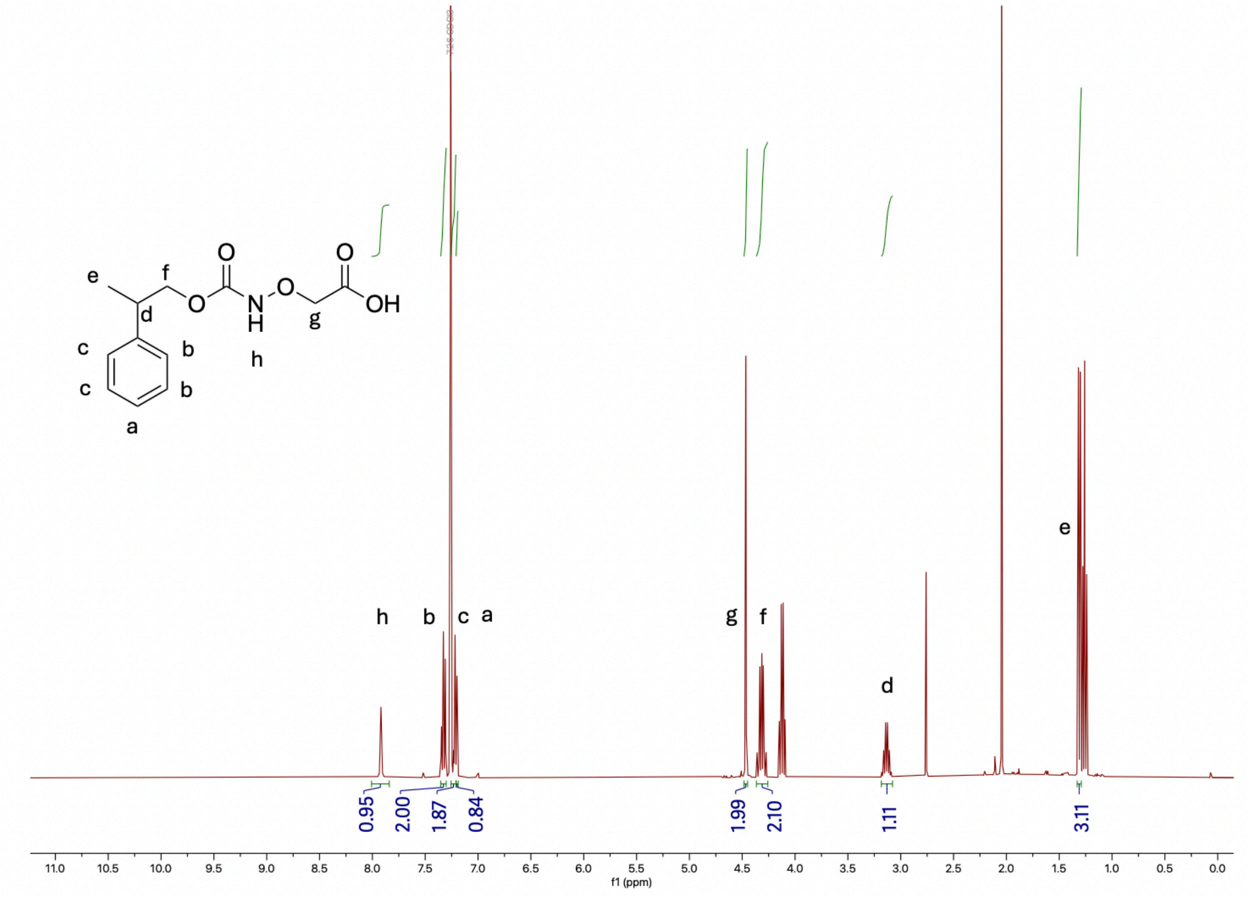

**Figure S33.** $^1$H NMR of 2-((((2-phenylpropoxy)carbonyl)amino)oxy)acetic acid (PPOC-HNO-COOH) in $CDCl_3$.

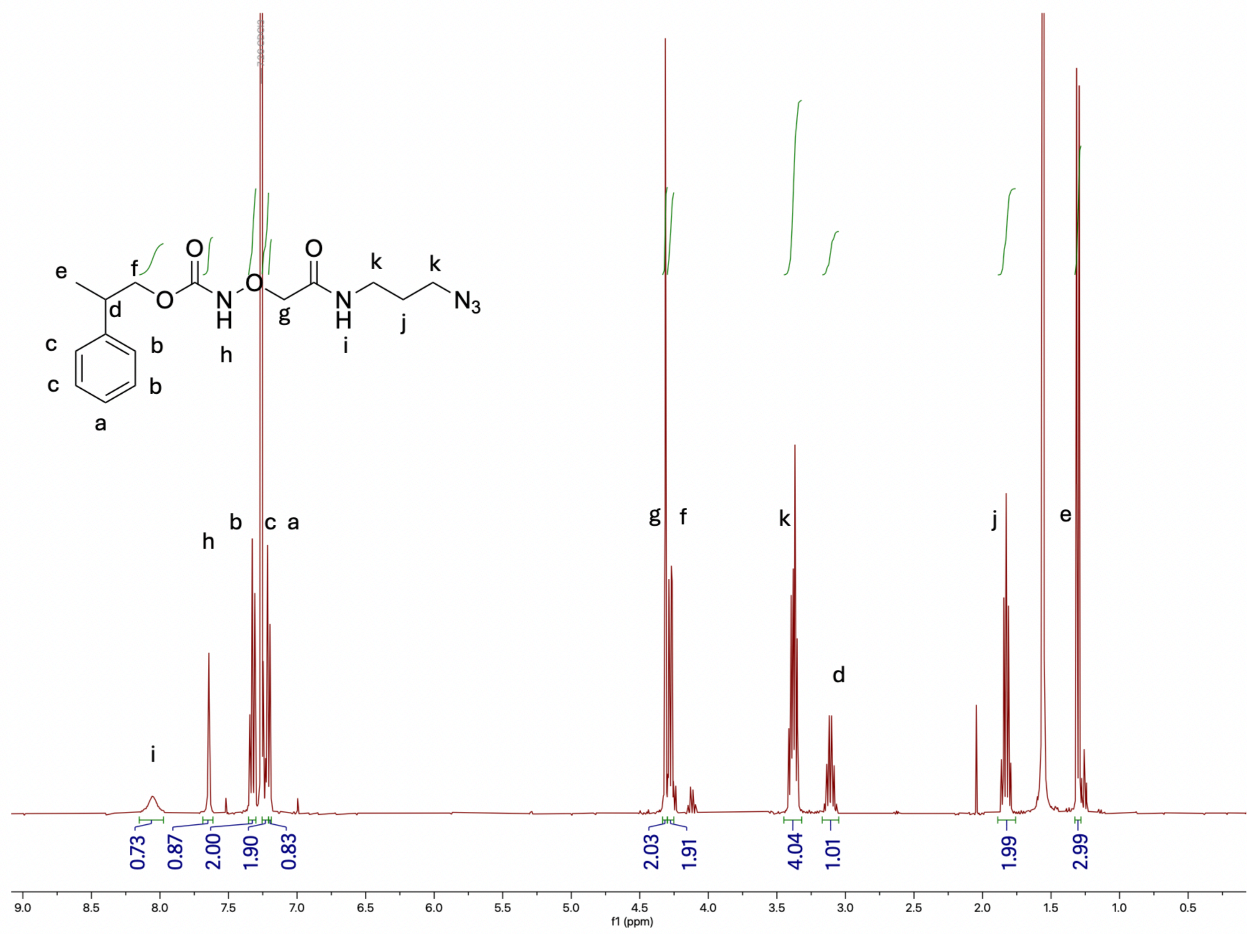

**Figure S34.** $^{1}$H NMR of 2-phenylpropyl (2-((3-azidopropyl)amino)-2-oxoethoxy)carbamate (PPOC-HNO-$N_3$) in $CDCl_3$.

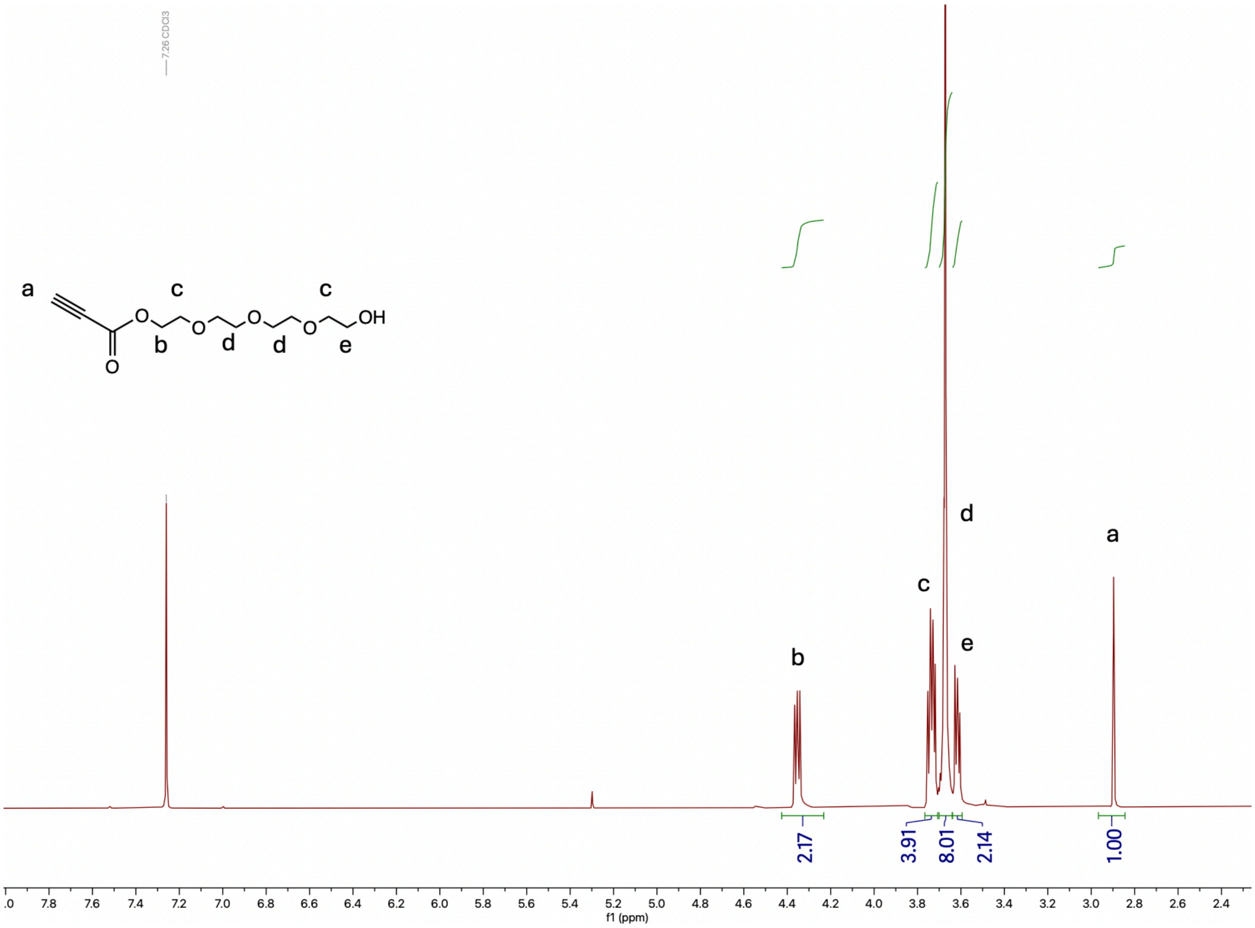

**Figure S35.** $^{1}$H NMR of 2-(2-(2-(2-hydroxyethoxy)ethoxy)ethoxy)ethyl propiolate in $CDCl_3$.

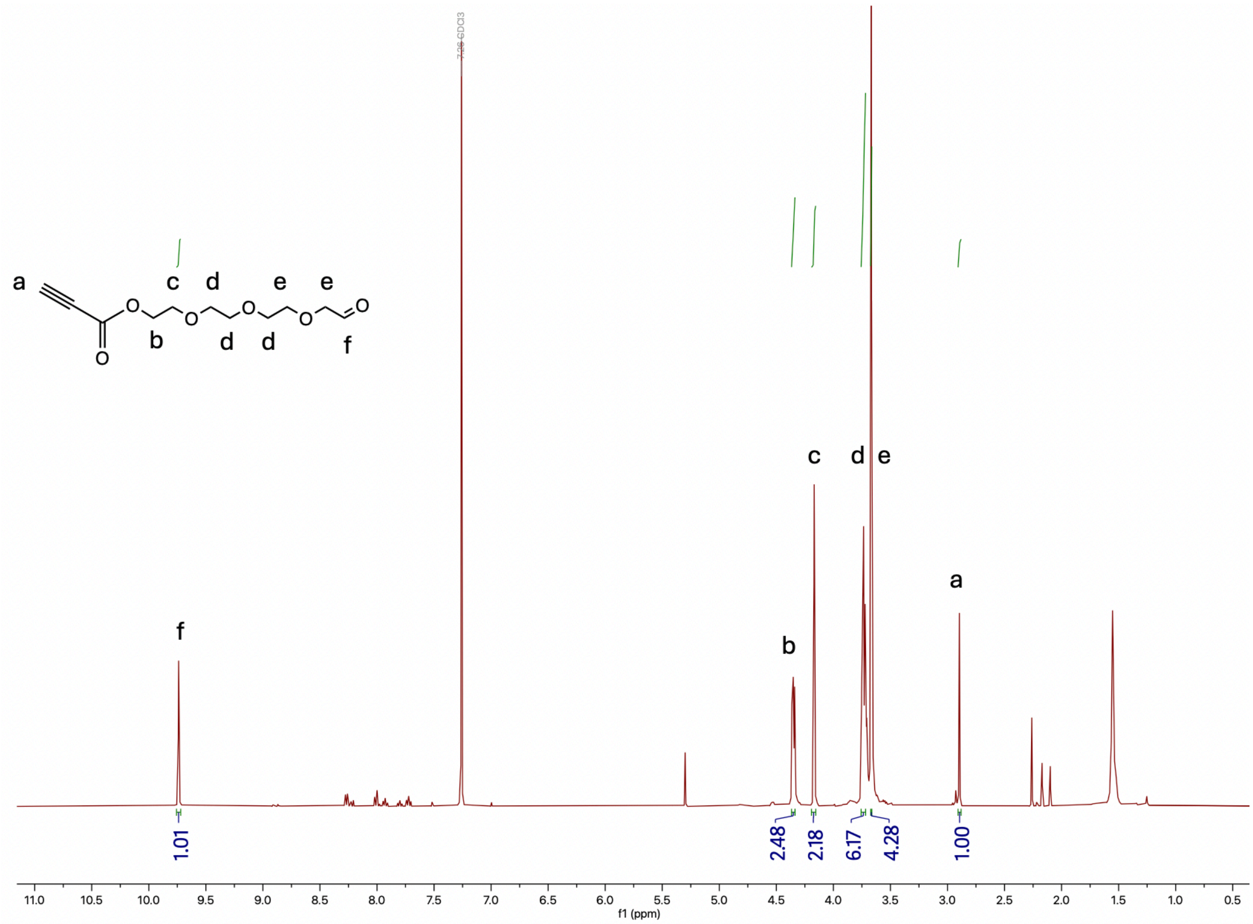

**Figure S36.** $^1$H NMR of 2-(2-(2-(2-oxoethoxy)ethoxy)ethoxy)ethyl propiolate in $CDCl_3$.

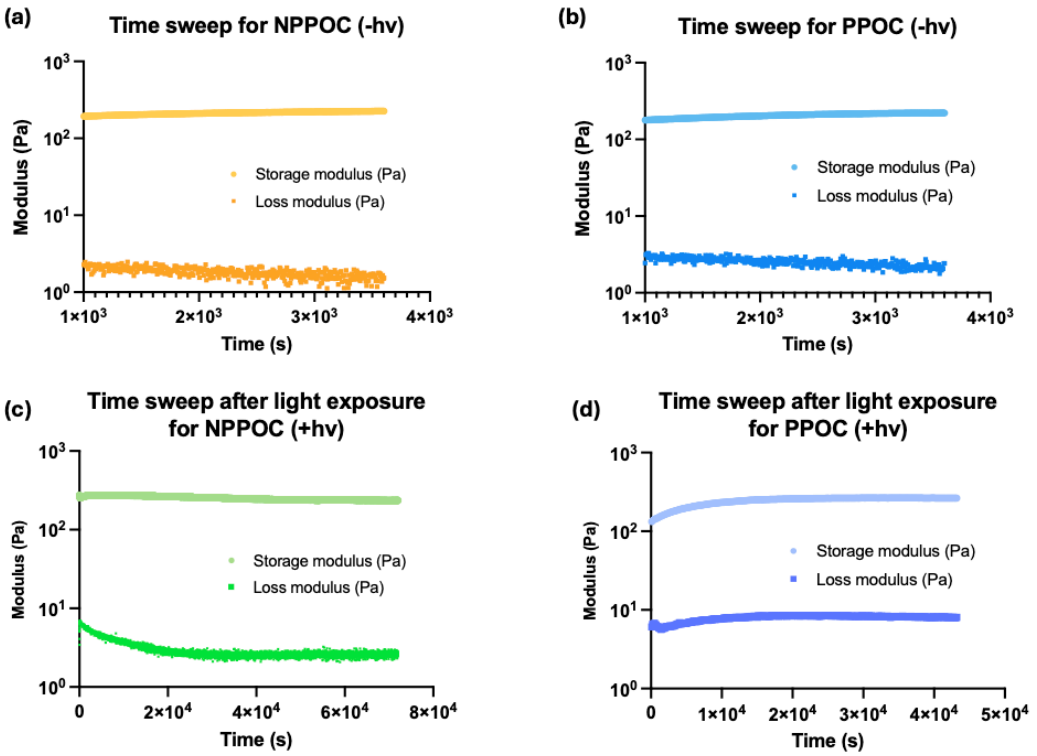

**Figure S37.** Representative time sweeps of ELP-PEG hydrogels post-gelation for (a) NPPOC- and (b) PPOC-protected PEGs without light treatment and (c) NPPOC- and (d) PPOC-protected PEGs with light treatment, showing storage moduli and loss moduli.

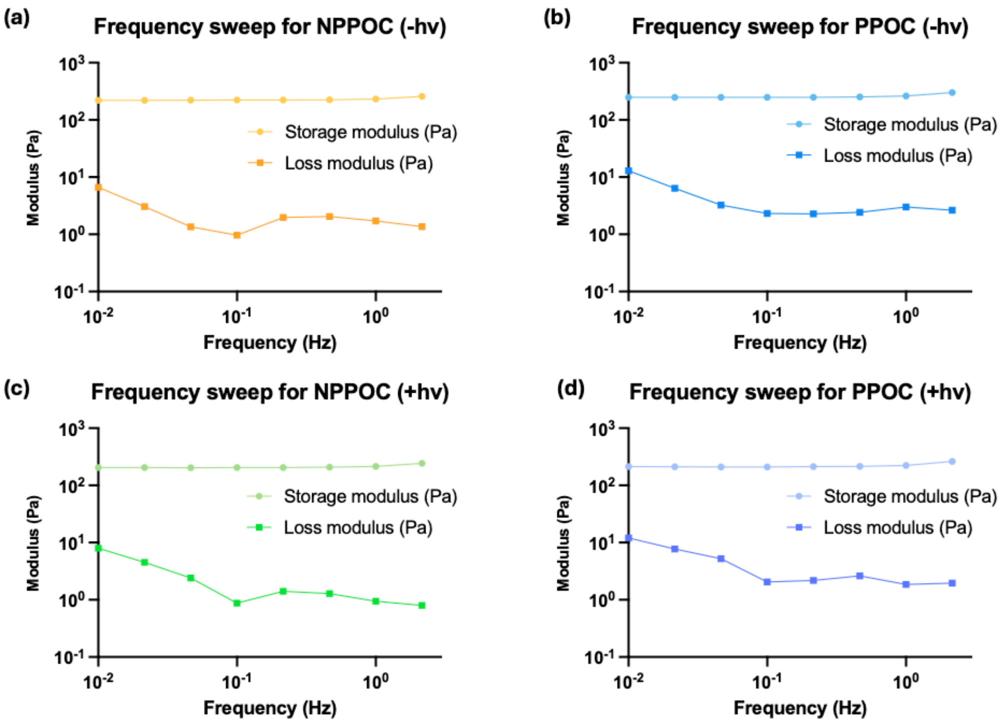

**Figure S38.** Representative frequency sweeps of ELP-PEG hydrogels for (a) NPPOC- and (b) PPOC-protected PEGs without light treatment and (c) NPPOC- and (d) PPOC-protected PEGs with light treatment.

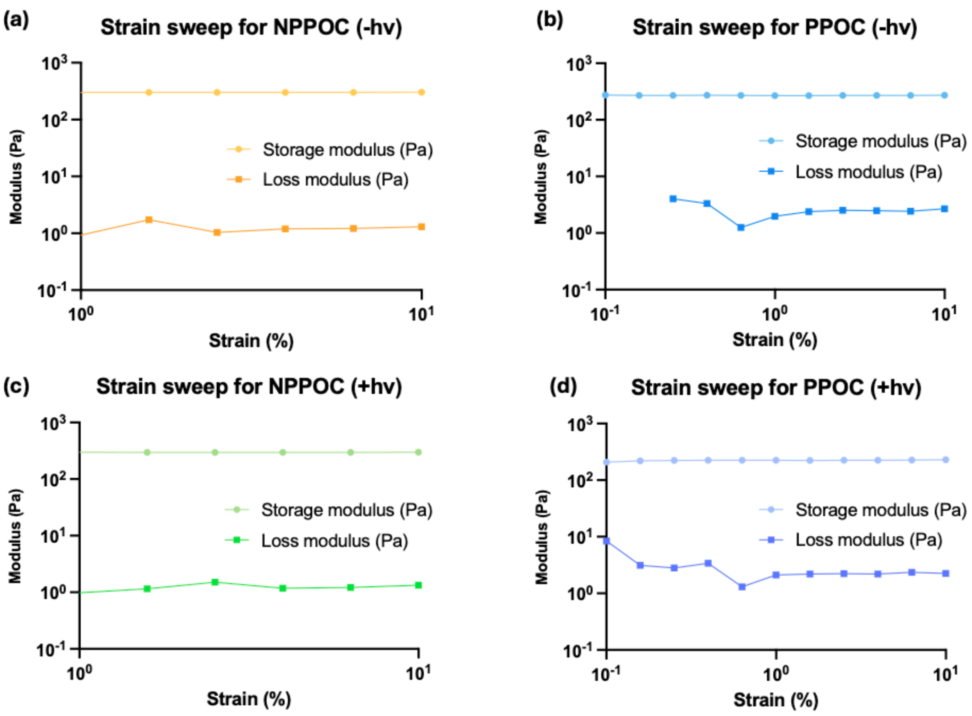

**Figure S39.** Representative strain sweeps of ELP-PEG hydrogels for (a) NPPOC- and (b) PPOC-protected PEGs without light treatment and (c) NPPOC- and (d) PPOC-protected PEGs with light treatment.

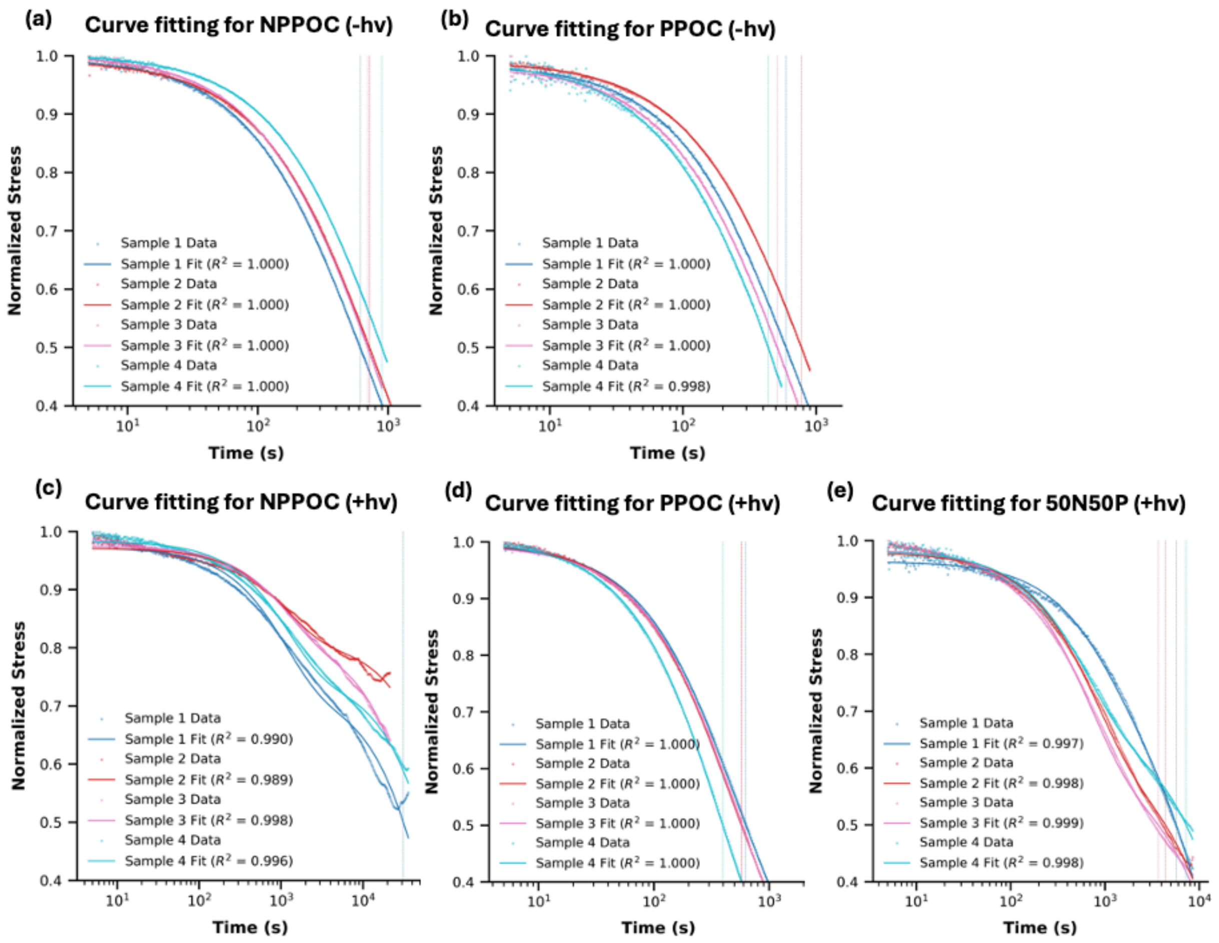

**Figure S40.** Curve fitting for stress relaxation tests of ELP-PEG hydrogels for (a) NPPOC- and (b) PPOC-protected PEGs without light treatment and (c) NPPOC-, (d) PPOC- and (e) a 1:1 blend of NPPOC- and PPOC-protected PEGs with light treatment.

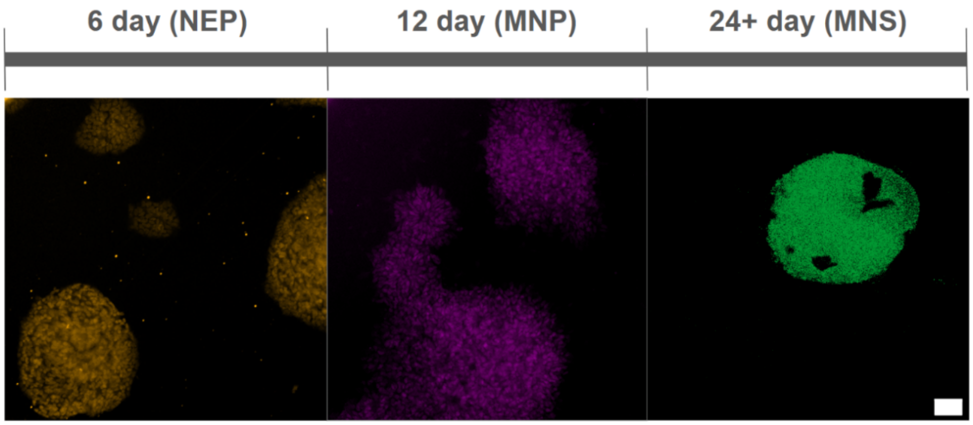

**Figure S41.** Immunofluorescence characterization of iPSC-derived motor neuron differentiation. Representative immunofluorescence images showing stage-specific marker expression during differentiation of induced pluripotent stem cells (iPSCs) into motor neurons. Neural progenitor cells (NEPs; 6 days post-differentiation) express the neural progenitor marker SOX1. Motor neuron progenitors (MNPs; 12 days post-differentiation) express the motor neuron progenitor marker OLIG2. Motor neuron spheroids (MNS; 6 days after spheroid formation) express the mature motor neuron marker MNX1, confirming successful progression through the motor neuron differentiation protocol. Scale bar = 50 μm.

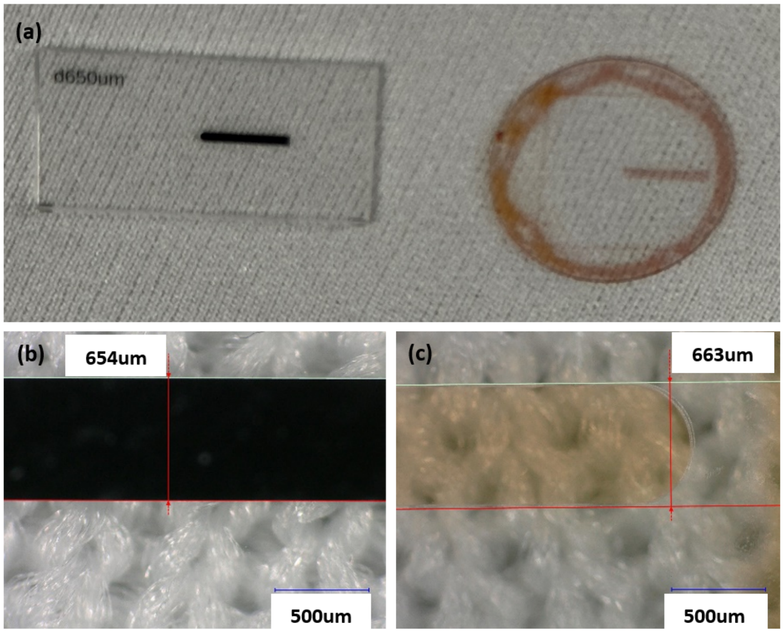

**Figure S42**. Fabricated photomask and evaluation of the resulting UV exposure pattern. (a) Photographs of the fabricated photomask and the developed AZ4620-coated quartz disk after UV exposure through the glass-bottom well plate. Optical micrographs of (b) 654 μm measured width of Cr on the photomask (c) 663 μm measured width of developed AZ4620 on the disk sample.

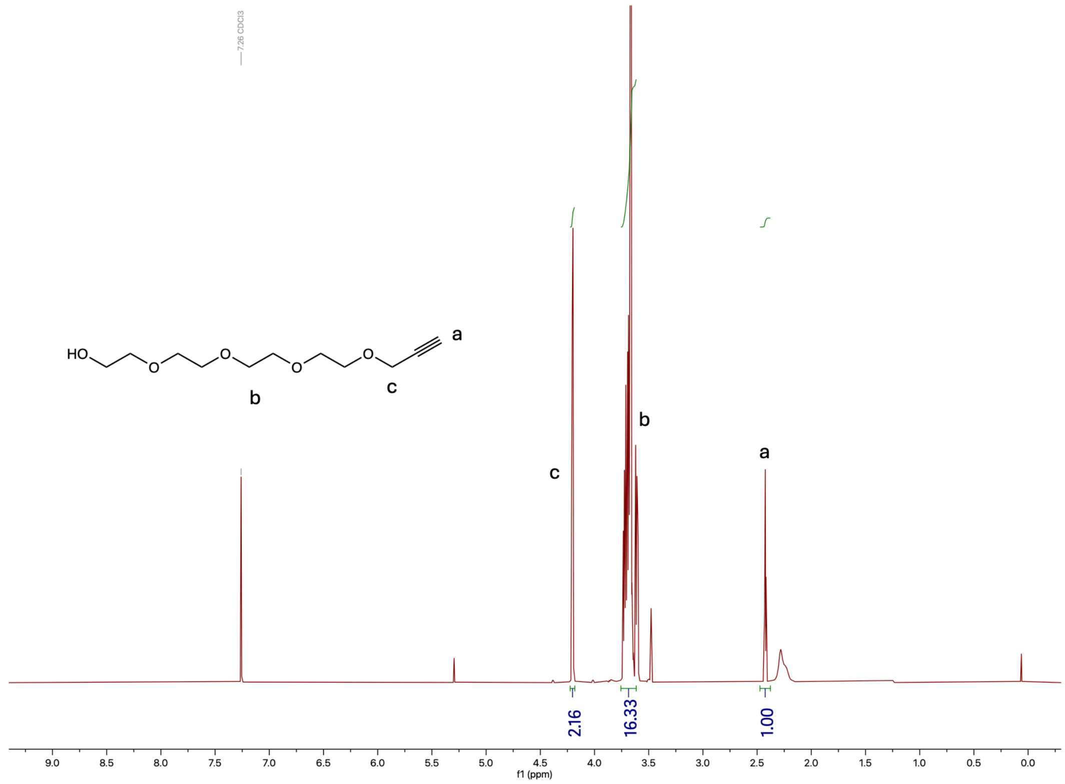

**Figure S43.** $^1$H NMR of 3,6,9,12-tetraoxapentadec-14-yn-1-ol in $CDCl_3$.

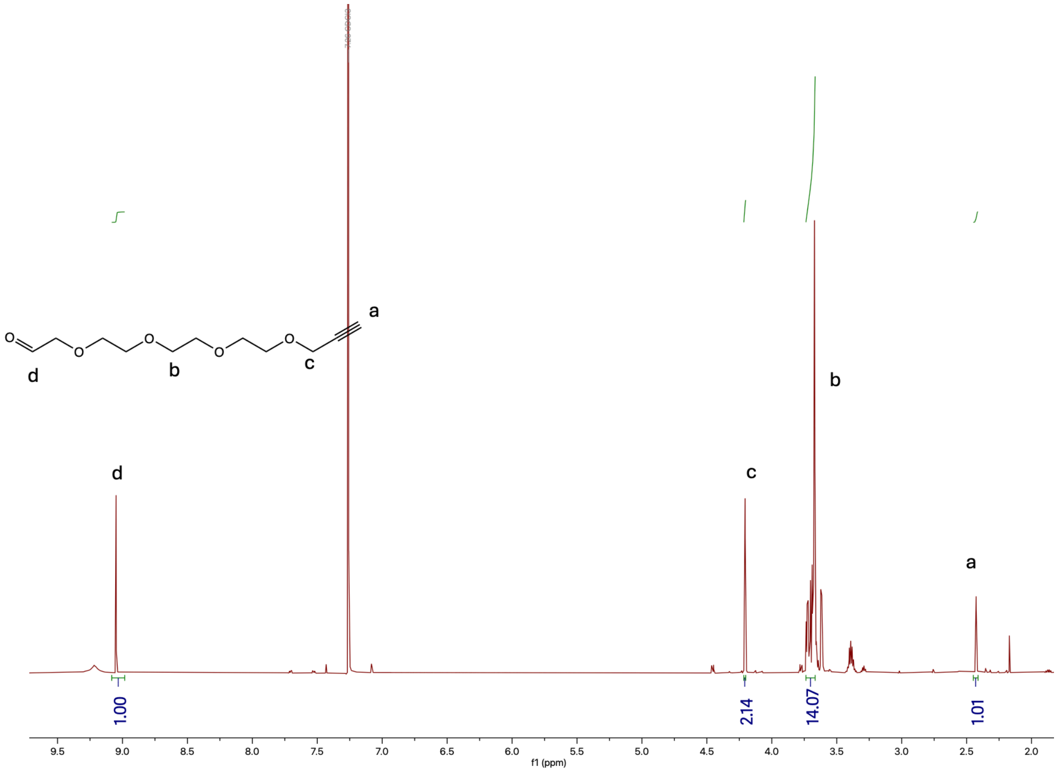

**Figure S44.** [1]H NMR of 3,6,9,12-tetraoxapentadec-14-ynal in $CDCl_3$.

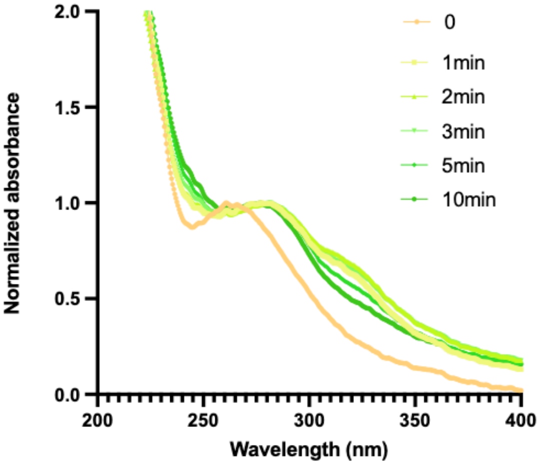

^2

**Figure S45.** UV-Vis spectra of a solution of 3.9 %(w/v) PEG-[8Hz/8HNO-NPPOC] in PBS irradiated with 365 nm light (50 mW/cm$^2$) for the indicated duration.

**Supporting Tables**

| **Bond type** | **$k_1$($M^{-1}s^{-1}$)** | **$k_{-1}$($s^{-1}$)** | **$K_{eq}$** | **Reaction conditions** | **Ref** |
|---|---|---|---|---|---|
| hydrazone | 23(5) | 2.8(6) $*10^{-4}$ | $8.1*10^4$ | 25 °C in pH 7.4 buffer containing 1% DMSO | [11] |
| hydrazone | 5.44 to 15 | $0.4*10^{-4}$ to $1.5*10^{-4}$ | ~$10^5$ | Room temperature, varied pH and with catalyst | [12] |
| hydrazone | 18.4 | $3.8*10^{-4}$ | $4.9*10^4$ | 25 μM reactant concentration at 37 °C in 0.1 M PBS buffer (pH 7.4) | [13] |
| hydrazone | 0.64 | $0.86*10^{-3}$ | 740 | Room temperature, pH 7.4 | [8] |
| oxime | 0.14 | $<10^{-9}$ | $>10^8$ | 25 °C, pH 7.4 | [14] |
| oxime | 0.12 | $1.5*10^{-7}$ | $8*10^6$ | Room temperature, pH 7.4 | [8] |

**Table 1.** Rate constants retrieved from published literature sources.

| **Target** | **Host Species (Isotype)** | **Supplier** | **Catalog Number** | **Dilution** |
|---|---|---|---|---|
| Nestin | Mouse (IgG1) | Millipore Sigma | MAB5326 | 1:600 |
| SOX2 | Rabbit (IgG) | Cell Signaling | 230645S | 1:400 |
| Ki67 | Rabbit (IgG) | Cell Signaling | 9129S | 1:400 |
| MAP2 | Rabbit (IgG) | Cell Signaling | 8707S | 1:400 |
| GFAP | Chicken (IgY) | AVES Labs | 87987979 | 1:300 |
| β-tubulin (III) | Rabbit (IgG) | Cell Signaling | 5568S | 1:500 |
| SOX1 | Goat (IgG) | R&D Systems | AF3369 | 1:200 |
| OLIG2 | Rabbit (IgG) | Millipore Sigma | AB9610 | 1:200 |
| MNX1 | Mouse (IgG1) | DSHB | 81.5C10 | 1:50 |

**Table 2.** Primary antibodies used in this study.

| Target | Conjugates | Supplier | Catalog Number | Dilution |
|---|---|---|---|---|
| Goat Anti-Mouse IgG1 | AlexaFluor 647 | Jackson ImmunoResearch | 115-605-205 | 1:600 |
| Goat Anti-Rabbit IgG (H+L) | AlexaFluor488 | Jackson ImmunoResearch | 111-545-144 | 1:400 to 1:500 |
| Goat Anti-Chicken IgY | AlexaFluor 647 | Jackson ImmunoResearch | 103-605-155 | 1:300 |
| Goat Anti-Rat IgG | AlexaFluor488 | Jackson ImmunoResearch | 112-545-167 | 1:500 |
| Donkey Anti-Mouse IgG (H+L) | Cy3 | Jackson ImmunoResearch | 715-165-150 | 1:500 |
| Donkey Anti-Goat IgG (H+L) | AlexaFluor488 | Jackson ImmunoResearch | 705-545-147 | 1:500 |

**Table 3.** Secondary antibodies used in this study.

| **Component** | **Neuronal Induction** | **MNP Patterning** | **MNP Expansion** | **MN Induction & Maturation (MN Medium)** |
|---|---|---|---|---|
| N-2 Supplement | 0.5x | 0.5x | 0.5x | 0.5x |
| B-27 | 0.5x | 0.5x | 0.5x | 0.5x |
| L-Ascorbic Acid | 100 uM | 100 uM | 100 uM | 100 uM |
| CHIR-99021 | 3 uM | 1 uM | 3 uM | -- |
| SB431542 | 2 uM | 2 uM | 2 uM | -- |
| DMH1 | 2 uM | 2 uM | 2 uM | -- |
| Retinoic Acid | 1 mM | 0.1 uM | 0.1 uM | 0.5 uM |
| Purmorphamine | 10 mM | 0.5 uM | 0.5 uM | 0.1 uM |
| Valproic acid | -- | -- | 0.5 uM | -- |
| Compound E | -- | -- | -- | 0.1 uM |
| BDNF/GDNF | -- | -- | -- | 10 ng/mL |

**Table 4.** The composition of motor neuron-related culture media.

### Supporting Video Captions

Representative widefield time-lapse microscopy recordings of fibroblast morphology on hydrogels formulated from ELP-ALD crosslinked with either photoreactive PEG-[8Hz/8HNO-NPPOC] or the photo-inert control PEG-[8Hz/8HNO-PPOC]. Samples with light treatment were irradiated with 365nm light at hour 6 for 90 seconds at a dose of 50 $mW/cm^2$. The videos highlight that, on the NPPOC hydrogel substrate, fibroblasts elongated after light irradiation, while on the PPOC substrate and on the substrates without light treatment, cell morphology was unchanged throughout the imaging duration. Images were acquired every 30 min over a total duration of 48 h (96 frames total). Scale bar: 500 μm.

**Movie S1.** PPOC-protected hydrogels without light treatment.

**Movie S2.** PPOC-protected hydrogels irradiated at t = 6 hours.

**Movie S3.** NPPOC-protected hydrogels without light treatment.

**Movie S4.** NPPOC-protected hydrogels irradiated at t = 6 hours.
